# cellGeometry: ultra-fast single-cell deconvolution of bulk RNA-Seq using a geometric solution

**DOI:** 10.64898/2026.01.24.701240

**Authors:** Rachel Lau, Cankut Çubuk, Athina Spiliopoulou, Pedro Martínez-Paz, Anna E. A. Surace, Liliane Fossati-Jimack, Soumya Raychaudhuri, Costantino Pitzalis, Myles J. Lewis

## Abstract

Single-cell analysis has rapidly expanded to produce cell atlases encompassing all human tissues. However, computational methods to deconvolute bulk samples using single-cell reference data have failed to keep pace with the increasing data size. Here we present cellGeometry, which uses non-negative geometric deconvolution (NGD), an intuitive vector projection method featuring non-negative matrix regularisation. Using matrix operations, cellGeometry scales to massive datasets and is ultrafast. Benchmarked using simulations from single-cell RNA-Seq datasets with >3 million cells, cellGeometry is more accurate than existing methods and more robust against noise simulating different sequencing chemistries. It identifies outlying residual genes which may unveil pathogenic changes in gene expression and the presence of cell types absent from the reference. cellGeometry’s flexible architecture allows merging of single-cell reference signatures to expand the range of cell types being deconvoluted. Validated against real bulk RNA blood and tissue samples, cellGeometry produces more accurate and realistic results.

## INTRODUCTION

Single-cell RNA-Sequencing (scRNA-Seq) measures gene expression in individual cells. It has become an essential foundation for understanding the web of interactions between the myriad cell subtypes that make up healthy tissues, as well as helping to unravel the complex pathogenic processes which lead to disease^1^. Advances in this technique have led to the curation of numerous large-scale datasets profiling millions of cells, such as Tabula Sapiens^2^ and the Human Brain Cell Atlas^3^, culminating in the Human Cell Atlas^4^. However, cost and stringent sample acquisition and storage requirements limit the sample size of scRNA-Seq studies. This is a real issue for clinical samples from large multicentre and/or multinational studies, where multisite sample processing for single cell data is impractical. In this scenario samples can be analysed by bulk RNA-Seq, but this lacks cell level information. With the compilation of annotated scRNA-Seq datasets in single-cell repositories^5^, deconvolution methods which use a reference single-cell dataset containing the cell types of interest have emerged as an important tool for extracting cell level information from bulk samples.

Currently, there are over 50 deconvolution methods [comprehensively reviewed by Nguyen *et al*^6^]. Deconvolution methods use a cell-specific gene signature, which is based on reference expression data, to predict cell type proportions in bulk samples. Older methods, which use a fixed, built-in reference gene signature^7,8^, limit analysis to the cell types contained within the reference, which typically only include broad hematopoietic cell types. Nguyen *et al* found that, of the newer deconvolution methods which allow a user-defined scRNA-Seq reference, MuSiC^9^, DWLS^10^ and LinDeconSeq^11^ were the top-performing methods in terms of accuracy when deconvoluting simulated pseudo-bulk RNA-Seq data. All three use weighted constrained least squares (W-CLS) for deconvolution, but with modifications. MuSiC applies gene weights based on cross-subject and cross-cell consistency. DWLS (dampened weighted least squares) applies a dampening constant to the weighting to prevent bias towards highly expressed genes or prevalent cell types with the aim of improving rare cell type identification. LinDeconSeq selects significantly highly expressed genes and assigns these to cell types through mutual linearity. DWLS and LinDeconSeq use quadratic programming to find their optimal solutions, while MuSiC’s optimisation strategy is comparable to iteratively reweighted least squares (IRLS). Other reference-based deconvolution strategies include deep neural networks^12^, support vector regression^8^ and Bayesian modelling^13,14^.

Existing deconvolution methods lack flexibility in accepting different scRNA-Seq data formats, especially the newer HDF5 anndata-based format which allows large datasets to be analysed without reading all the dataset into the programming environment, greatly reducing memory consumption and runtime.

Most current workflows do not incorporate quality check functions to assess the cell type-specific gene signatures. This is problematic when cell types are similar and therefore lack sufficiently differentiating cell-specific markers. Therefore, considering the imperfections of gene signatures and mitigating nonsensical cell abundances (such as negative values) are fundamental to the reliability of deconvolution algorithms. Failure to fully consider these issues leads to some deconvolution methods failing with real-world data despite performing well with simulated and canonical benchmarking datasets.

We present cellGeometry, which uses a geometric solution for accurate deconvolution, and whose code is highly optimised to be ultra-fast. Cell frequencies are deconvoluted by vector projection of bulk RNA-Seq gene expression against cell cluster vectors in high-dimensional gene space with non-negative compensation for spillover. Using simulations generated from diverse reference single-cell datasets, cellGeometry outperformed existing top-performing methods^6^. In real-world bulk RNA-Seq of blood and tissue samples cellGeometry removes outlying genes with extreme residuals leading to improved deconvolution accuracy.

## RESULTS

### Modular approach to deconvolution

cellGeometry deliberately splits the core components of deconvolution into stages (**Fig. 1a**). Its code has been implemented with flexible modular functions allowing users significant control over each stage of the deconvolution process. Users provide i) the count matrix of a single-cell RNA-Seq dataset, with ii) predefined cell clusters, and iii) the count matrix of bulk RNA-Seq samples to be deconvoluted. The two main stages are generation of a cell-specific gene signature matrix, followed by the deconvolution itself. Separation of the cell cluster mean gene expression computation from deconvolution means that users only have to run this step once, which is the slowest computational step. After the initial run, signatures can be rapidly modified and deconvolution parameters changed without having to regenerate the cell cluster mean expression. Users can refine the signatures through adding/subtracting genes, removing overlapping cell clusters, or even adding cell clusters by merging them from other single-cell datasets. cellGeometry provides model diagnostics, in which gene expression residuals and deconvolution standard errors can be examined.

**Fig. 1.**
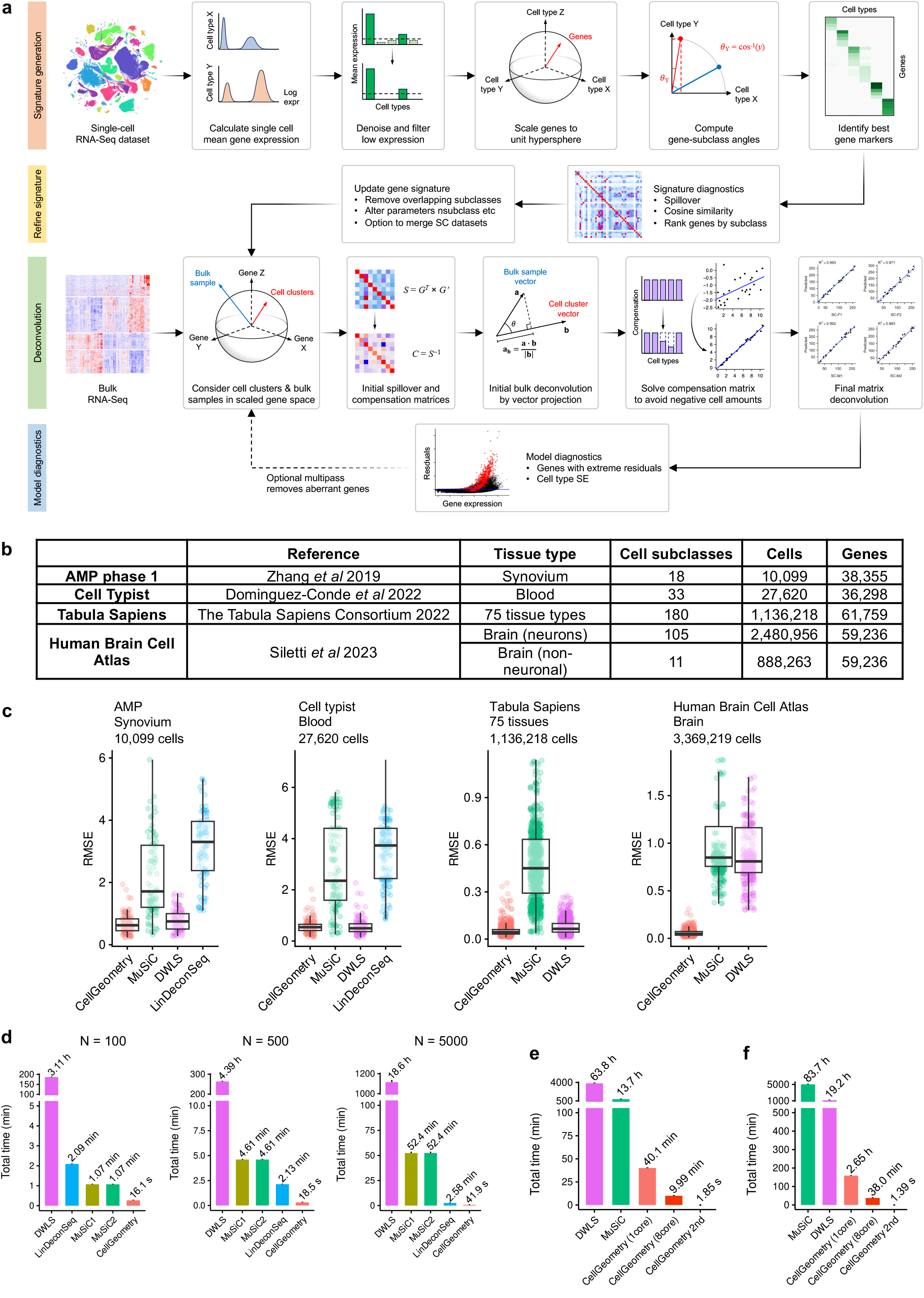
Benchmarking accuracy and speed of deconvolution of simulated pseudo-bulk data from four increasingly large reference single-cell datasets. **a**, cellGeometry generates a cell-specific gene signature matrix based on a single-cell RNA-Seq dataset by calculating the mean gene expression per cell type following denoising and filtering to remove low expressed genes. Optimal cell-specific genes are identified by considering each gene as a vector with cell clusters as dimensions and determining which genes are closest to the dimensional axis for that cell subclass. Diagnostic tests and refinement of the gene signature can be performed after signature generation prior to deconvolution. The second stage of the workflow is the deconvolution process whereby the vector projection of each bulk RNA-Seq sample is calculated against a vector representing each cell cluster in high dimensional gene marker space using the vector dot product, followed by compensation for spillover which is solved to avoid negative cell quantities. **b**, Table of reference single-cell RNA-Seq datasets used for benchmarking. **c**, Boxplots of root mean square error (RMSE) of cell subclass percentages from deconvolution of simulated pseudo-bulk datasets generated from rheumatoid arthritis synovium^15^, Cell Typist blood^18^, Tabula Sapiens^2^ and Human Brain Cell Atlas^3^. The deconvolution methods assessed were cellGeometry, MuSiC, DWLS and LinDeconSeq. Five replicates of simulated pseudo-bulk data (N = 25 each) were generated, with the exception of Human Brain Cell Atlas where 3 replicates (N = 30 each) were generated. Boxplots illustrate median, upper and lower quartiles with whiskers denoting maximal and minimal data within 1.5× interquartile range. *y* axis is cropped at the maximum whisker values to reduce plot distortion by outliers. **d**, Barplots of the average running time of deconvoluting RA synovium simulated datasets using cellGeometry, MuSiC1, MuSiC2, DWLS and LinDeconSeq with different sample sizes (range 100-5000) and 5 repeats. As MuSiC1 and MuSiC2 produced identical results, MuSiC2 was used throughout the rest of the study. **e**, Barplot of the average running time of deconvoluting Tabula Sapiens simulated dataset using cellGeometry with 1 core or 8 cores, MuSiC and DWLS. With cellGeometry’s feature to update the gene signature matrix, the running times were also compared with a 2nd run of deconvolution by cellGeometry (8 cores) with modified settings (nsubclass changed from 500 to 200). Five repeats were undertaken. **f**, Barplot illustrating the average running time of deconvoluting the simulated pseudo-bulk from the Human Brain Cell Atlas using cellGeometry with 8 cores, MuSiC and DWLS. 2^nd^ run of deconvolution (8 cores) was also undertaken with different settings (nsubclass changed from 500 to 200). Three repeats were undertaken.

### Vector angles identify cell-specific gene markers

cellGeometry first generates a cell type-specific gene expression signature matrix by calculating the mean gene expression in each cell cluster in the user-defined reference scRNA-Seq dataset. Each gene is considered as a vector in high-dimensional space with cell types as dimensions (**Supplementary Fig. 1a-b**). Since gene expression is non-negative, the gene vectors occupy the non-negative orthant. The most specific gene markers for a specific cell type are genes whose vector lies closest to the dimensional axis for that cell type of interest. Thus, genes can easily be ranked in order of cell type specificity by this angle. If each gene is scaled to the unit hypersphere in cell type dimensional space, the specificity angles are easily computed by applying arc cosine to the unit sphere-scaled gene expression matrix.

This is illustrated as a simplified example in **Supplementary Fig. 1c** using the Accelerated Medicines Partnership (AMP) scRNA-Seq study of rheumatoid arthritis (RA) synovial tissue^15^ where the mean gene expression for three cell types are visualised: B cell (*z* axis), monocyte (*x* axis) and fibroblast (*y* axis). *CD22, MS4A1* (CD20) and *CD79A* are highly expressed in B cells but show very low expression in monocytes and fibroblasts; hence, each of these genes, which are known B-cell specific markers, are closest to the B cell axis (*z* axis).

With gene signature curation, a low expression filter is applied to remove unreliable scRNA-Seq genes exhibiting apparent cell specificity purely by chance, i.e. with very rare counts which happen to only occur in one cell type. For real bulk deconvolution, an optional noise filter can also be applied which denoises the scRNA-Seq reference signature matrix.

### Non-negative geometric deconvolution

After the generation and diagnostics of the gene expression signature matrix (example in **Supplementary Fig. 2**), bulk RNA-Seq data is deconvoluted using vector projection, which is calculated using the dot product. In essence, the gene expression in each bulk sample is represented as a vector in gene expression space, and projected against each cell cluster vector, representing the mean gene expression per cell (**Supplementary Fig. 1d-e**). Here, *genes* are dimensions, in contrast to signature generation where *cell types* are dimensions. Sphere scaling of genes means that each gene has equal weighting in the vector projection, so that the process is not dominated by the most highly expressed genes.

As genes are almost never 100% cell specific, the cell type vectors are non-orthogonal which is similar to the problem of spillover in fluorescence activated cell sorting (FACs)^16^. Hence the spillover in vector projection (**Supplementary Fig. 1e**) is adjusted by applying a compensation matrix (**Supplementary Fig. 3**), which is solved to prevent negative cell abundance. This variable compensation, controlled on a per-cell type basis, preserves the relative relationship for each cell cluster to their respective gene signatures so that increased gene marker expression corresponds to linearly increased cell type abundance (further details in the Methods) (**Supplementary Fig. 4**). As NGD deconvolutes the bulk samples together rather than independently, NGD preserves linear proportionality between the cell-specific gene expression and the cell subclass abundance across bulk samples, which is desirable for end users. Hence even if there are problems (experimental noise/bias) between SC and bulk matrices, NGD will still yield results for each subclass that are sensible by being proportional to each cell-specific signature.

While our method has a geometric derivation, and hence we refer to it as non-negative geometric deconvolution (NGD), we show in the Methods that it is equivalent to constrained generalised Tikhonov regularisation with a novel heuristic solution for the regularisation matrix driven by non-negativity of cell counts rather than cross-validation.

### cellGeometry demonstrates optimal accuracy with simulated data

We benchmarked cellGeometry against reference based-deconvolution methods that were considered top performers by Nguyen *et al*^*6*^: MuSiC^9,17^, DWLS^10^ and LinDeconSeq^11^. We constructed simulated pseudo-bulk datasets with known true cellular subclass proportions for benchmarking using three scRNA-Seq and one single-nucleus RNA-Seq (snRNA-Seq) dataset. These were selected to cover complex healthy tissues including blood immune cells (Cell Typist^18^), all human tissues (Tabula sapiens^2^) and brain (Human Brain Atlas single-nucleus^3^), as well as disease tissue with inflammatory cell infiltration (RA synovial tissue^15^) (**Fig. 1b**). These datasets contain a wide range of cell amounts (10,000-3,000,000 cells) and cell type numbers (18-180 subclasses). DWLS and LinDeconSeq are not capable of handling matrices with >2^31^ elements for scRNA-Seq input, whereas the datasets Tabula Sapiens and the Human Brain Cell Atlas are massively larger than 2^31^. To stay within the 2^31^ matrix size limit affecting DWLS and LinDeconSeq, we downsampled the cell subclasses randomly in the reference datasets, while ensuring that cell subclasses with fewer cells were sufficiently sampled. This reduced these >1 million cell datasets to ~20,000-30,000 cells. DWLS successfully processed both subsetted scRNA-Seq datasets. But LinDeconSeq was only able to process the Brain Atlas non-neuronal snRNA-Seq dataset, since it failed at the signature generation step for Tabula Sapiens and the Brain Atlas neuron datasets.

Accuracy measured by root mean square error (RMSE) (**Fig. 1c**), the coefficient of determination (R^2^) (**Supplementary Fig. 5**) and Lin’s Concordance Coefficient (CCC) (**Supplementary Fig.6, Supplementary Table 1**) was highest for cellGeometry across all four datasets and comparable to DWLS for RA synovium, Cell Typist blood and Tabula Sapiens. On the other hand, MuSiC and LinDeconSeq performed significantly less well. Scatter plots of the predicted and observed cell subclass proportions from one repeat for each deconvolution method are shown for RA synovium (**Supplementary Fig. 7**), Cell Typist blood (**Supplementary Fig. 8**), Tabula Sapiens (**Supplementary Fig. 9**), Human Brain Cell Atlas neurons (**Supplementary Fig. 10**) and Human Brain Cell Atlas non-neuronal cells (**Supplementary Fig. 11**).

cellGeometry also showed consistently high accuracy across all subclasses and cell groups of the RA synovium and Cell Typist blood dataset and the distributions of the estimated cellular proportions were similar to the ground truth for the two datasets (**Supplementary Fig. 12-13**). Notably, MuSiC failed to detect several T cell subsets or granulocyte-monocyte progenitors in Cell Typist blood, 16 subclasses in Tabula Sapiens and 63 neuron subclasses in the Human Brain Cell Atlas. MuSiC did not output these subclasses or reported zero cell proportions despite these subclasses being present in the simulated pseudo-bulk data. LinDeconSeq also failed to detect 3 subclasses in Cell Typist blood.

For the atlas-scale datasets (Tabula Sapiens, Human Brain Cell Atlas), we re-ran MuSiC using randomly downsampled cell subclasses in the reference datasets, but this did not significantly improve its accuracy (**Supplementary Table 1**). Also, we tested deconvolution tools purportedly designed for large-scale sc/snRNA-Seq studies (Bisque^19^ and InstaPrism^14^) on the atlas-level datasets and found that they did not perform as well as cellGeometry (**Supplementary Fig. 9-11, Supplementary Table 1**).

In addition, cellGeometry was consistently more accurate at deconvolution than other popular mathematical methods, including non-negative least squares (NNLS), non-negative elastic net regression (glmnet) with varying alpha (including both LASSO and ridge regression), and non-negative matrix factorisation (NMF) using the Brunet, Kullback-Leibler, Lee and DeBruine algorithms^20,21^ (**Supplementary Fig. 14**).

### cellGeometry is ultra-fast compared to other methods

With increasing sample size of RA synovium^15^ simulated data, cellGeometry was substantially faster than the other deconvolution methods because it uses matrix operations where all bulk samples are deconvoluted simultaneously (**Fig. 1d**). DWLS, LinDeconSeq and MuSiC deconvolute samples one by one; so as the bulk sample size increases the deconvolution runtime increases linearly. With 5000 samples, cellGeometry was ~1600 times faster than DWLS (41.9 seconds vs 18.6 hours).

The speed increase with cellGeometry is clearer still with larger scRNA-Seq datasets. Using Tabula Sapiens^2^ simulated data, cellGeometry (8 cores) is ~80 times faster than MuSiC (9.99 minutes vs 13.7 hours) and ~380 times faster than DWLS (9.99 minutes vs 63.8 hours (**Fig. 1e**). Even using a downsampled scRNA-Seq dataset, the signature curation alone by DWLS took 54.6 hours. As previously mentioned, users can refine the gene signatures in cellGeometry without having to rerun the rate-limiting step of cell cluster gene means calculation. The 2^nd^ run of cellGeometry with updated settings, i.e. the deconvolution step alone, was substantially faster (1.85 seconds) than the initial run.

With the Human Brain Cell Atlas, cellGeometry (8 cores) took a total of 38.0 minutes to deconvolute a combined simulated pseudo-bulk dataset of neurons and non-neuronal cells (**Fig. 1f**). In comparison, MuSiC, which is a one-step process that does not allow for parallelisation, took a total of 79.6 hours to deconvolute neurons plus 4.15 hours to deconvolute non-neuronal cells (~130 times slower overall than cellGeometry). DWLS took a total of 18.1 hours to deconvolute neurons plus 1.07 hours to deconvolute non-neuronal cells. The 2^nd^ run of deconvolution by cellGeometry with updated settings was considerably faster (1.39 seconds) than the initial run.

The main reasons for cellGeometry’s considerable speed gains are: i) during signature generation, computation of mean gene expression per cell cluster across large, highly sparse scRNA-Seq matrices is optimised through the use of DelayedArray subroutines^22^; ii) fork-based parallelisation during gene signature generation; and iii) deconvolution is performed on all bulk samples and all cell clusters simultaneously so that computations are executed entirely through large matrix operations such as R’s matrix cross-product which harness optimised basic linear algebra subroutine (BLAS) libraries^23^.

### Cell subclass similarity imposes a limit on deconvolution accuracy

With Tabula Sapiens^2^, cellGeometry had much greater accuracy across all subclasses and cell groups compared to MuSiC (**Fig. 2a, Supplementary Fig. 15**). DWLS had similar deconvolution accuracy to cellGeometry. Some cell clusters defined in Tabula Sapiens have high similarity with each other, as shown by a cosine similarity heatmap (**Fig. 2b**). For example, related subtypes “CD4^+^ aβ T cell” and “naive thymus-derived CD4^+^ aβ T cell” show high similarity, “fibroblast” and “thymic fibroblast type 2” are highly similar, and “monocyte”, “intermediate monocyte” and “classical monocyte” also show high similarity with each other. These overlapping cell clusters tend to have relatively lower deconvolution accuracy with cellGeometry (**Fig. 2c**). We observed a reciprocal relationship (pseudo-R^2^=0.64) between deconvolution error as measured by RMSE and (1 – max similarity). This observed trend is consistent with the algorithm’s assumption that each cell subtype is distinct and mutually exclusive.

**Fig. 2.**
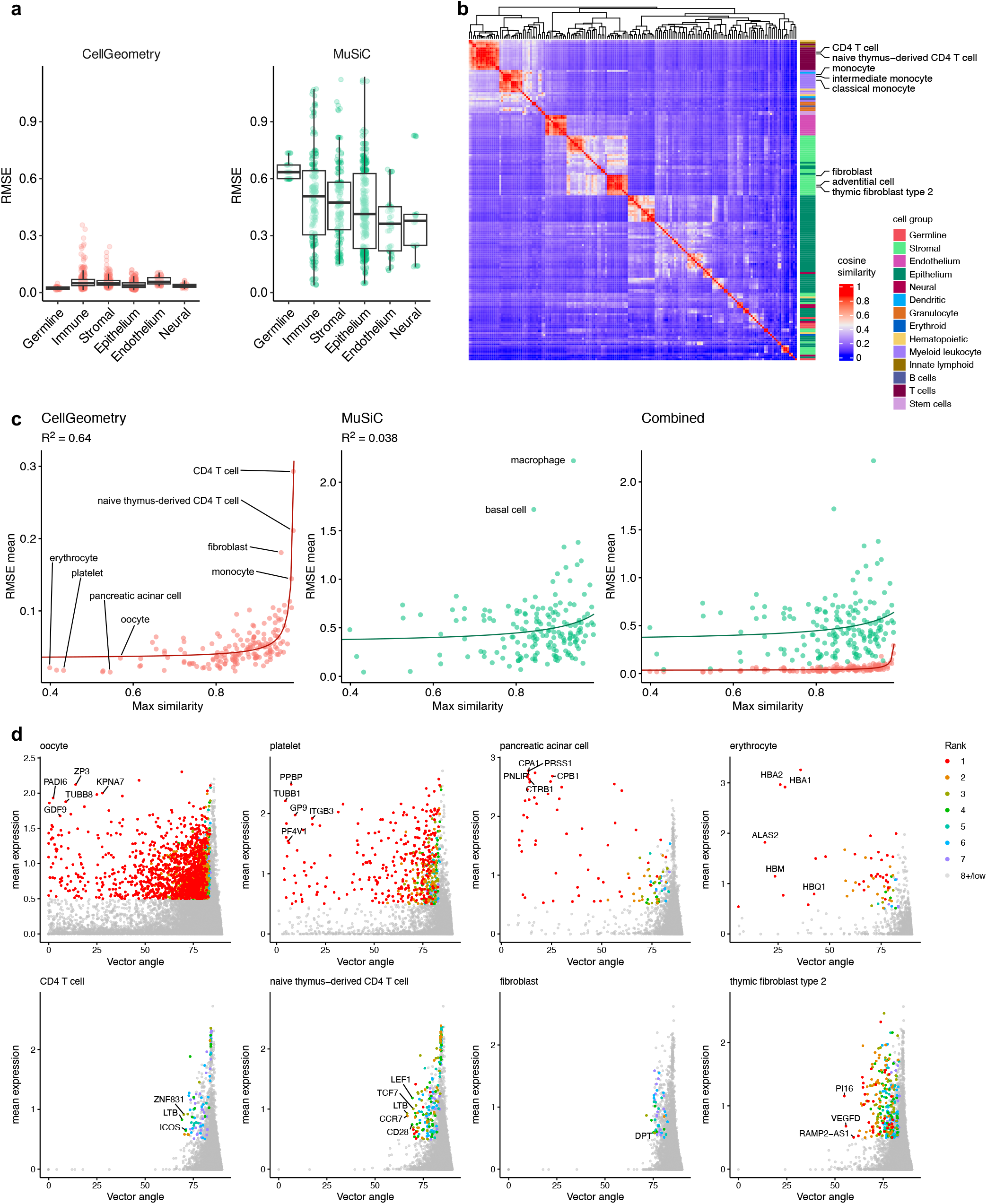
Deconvolution can be improved by removing overlapping cell clusters with insufficiently dissimilar gene signatures. **a**, Boxplots of root mean square error (RMSE) for cell subclass percentages from deconvolution using cellGeometry, MuSiC and DWLS on 5 replicates of Tabula Sapiens^2^ simulated pseudo-bulk data (N = 25). Boxplots illustrate median, upper and lower quartiles with whiskers denoting maximal and minimal data within 1.5× interquartile range. *y* axis is cropped at the maximum whisker values to reduce plot distortion by outliers **b**, Heatmap showing cosine similarity of cell subclasses. **c**, Mean RMSE per cell subclass from deconvolution of 5 replicates of Tabula Sapiens simulated data by cellGeometry and MuSiC plotted against the maximum cosine similarity for each cell subclass. **d**, Vector angle specificity plots of mean gene expression (*y* axis) and vector specificity angle (*x* axis) (see Methods). Colours show ranking of each gene by mean gene expression across cell subclasses.

Where cell types are strongly distinct from one another and have recognisably distinct cell-specific markers, the vector method used by cellGeometry identifies these cell-specific markers with ease using vector angle specificity plots. These are shown for oocyte, platelet, pancreatic acinar cell and plasma cell in Tabula Sapiens (**Fig. 2d**, upper row). For instance, *GDF9* is an oocyte-specific factor critical for ovarian folliculogenesis^24^; *GP9* is a platelet receptor for von Willebrand factor and *GP9* mutations result in giant platelets^25^; *PRSS1* is a digestive enzymes that is synthesised and secreted by pancreatic acinar cells^26^; and *HBA1/2* are haemoglobin α subunits expressed in erythrocytes to transport oxygen.

In contrast, it is harder for the algorithm to identify suitably specific gene markers for similarly named and clustered cell subclasses (**Fig. 2d**, lower row). So although Tabula Sapiens considers the different tissue sampling in the naming and clustering of the different cell types, the reality is that some cell types may lack differentiating unique cell markers between tissues and may be difficult to reliably deconvolute in bulk data.

Therefore, how the reference scRNA-Seq clustering and resultant cell subtypes are defined needs to be strongly considered. Cell types that overlap too closely with another can be removed from the signature matrix in the workflow of cellGeometry.

### Ability to merge single-cell reference signatures

The Human Brain Cell Atlas is a good example of a challenging snRNA-Seq dataset to deconvolute due to the close similarity in gene expression between neuron subtypes. A unique feature of cellGeometry is the ability to merge two different single-cell cluster signatures and redefine the optimal gene signatures. This is demonstrated through the Human Brain Cell Atlas^3^ which contains neurons and non-neuronal cells in separate files. Simulated pseudo-bulk samples containing random mixtures of neurons and non-neuronal cells were generated (**Fig. 3a-c**). MuSiC, DWLS and LinDeconSeq do not have this feature and so the predicted percentages can only consider neuron and non-neuronal separately, which is poorly representative of the actual percentages of the simulated data. Consequently, cellGeometry had substantially greater accuracy than the other methods (**Fig. 3d-e, Supplementary Fig. 16-17**). LinDeconSeq was unable to generate a signature for Human Brain Cell Atlas neurons so could only deconvolute the non-neuronal cells.

**Fig. 3.**
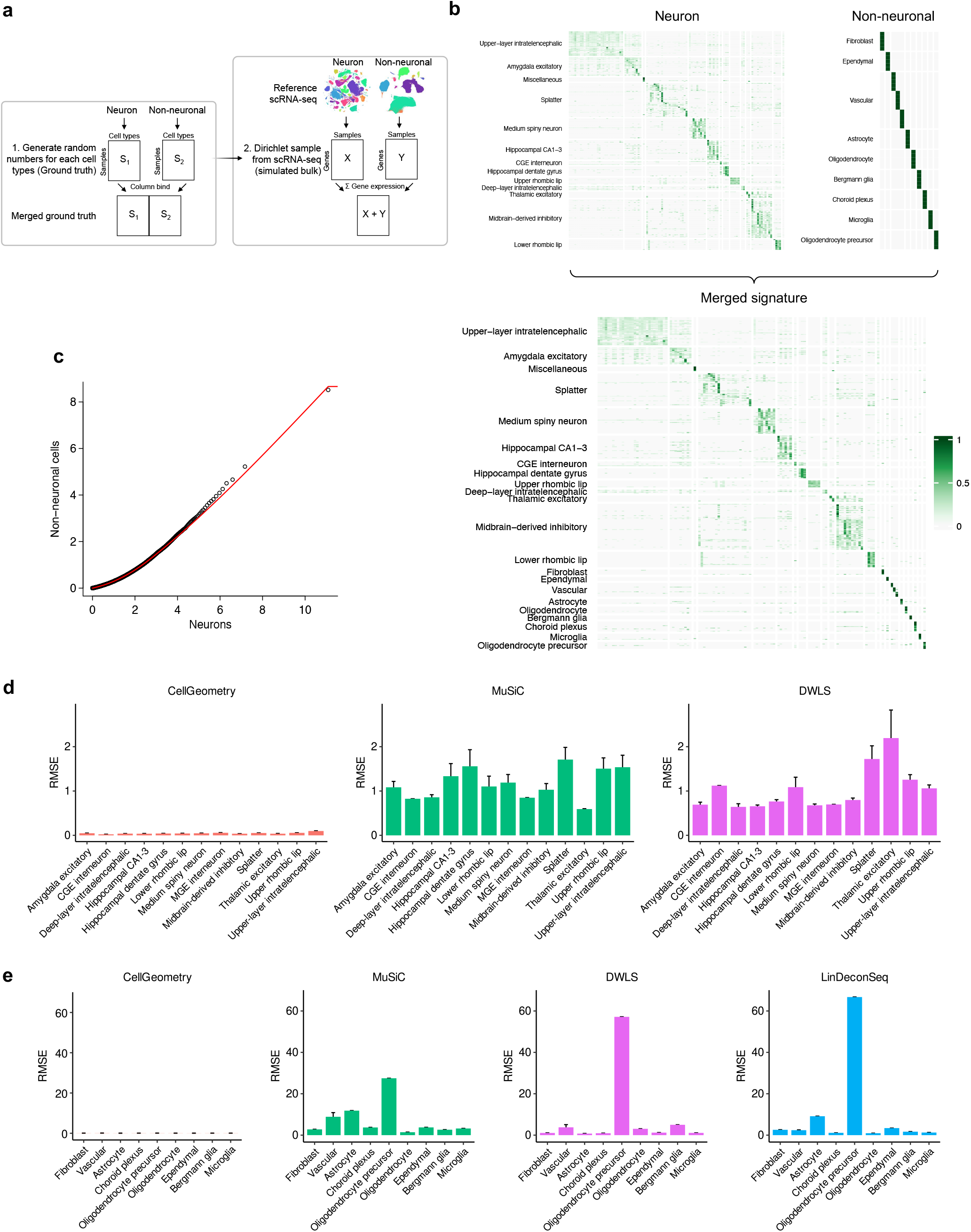
Merging single-cell reference signatures enables deconvolution of complex tissues such as brain. **a**, Flow diagram showing the generation of random cell numbers (ground truth) and sampled pseudo-bulk RNA-Seq counts based on Human Brain Cell Atlas^3^ neuron and non-neuronal single-cell datasets, which are stored in separate files, so ground truth cell numbers were merged through column binding. Simulated pseudo-bulk RNA-Seq was generated through Dirichlet based sampling of each scRNA-Seq dataset separately. The final pseudo-bulk count matrix was combined by summation. **b**, Gene signature curation workflow by cellGeometry whereby neuron and non-neuronal gene signatures from Human Brain Cell Atlas were merged. **c**, Quantile-quantile plot on log_2_+1 scale of neuron and non-neuronal mean gene expression per cell type. **d, e**, Barplot of the mean root mean square error (RMSE) for deconvoluted cell subclass percentages from pseudo-bulk simulation of the Human Brain Cell Atlas showing **(d)** neurons and **(e)** non-neuronal cells. Three replicates of simulated bulk data (N = 30 samples) were generated. Error bars represent the standard error of the mean (SEM).

### Testing robustness of deconvolution methods to experimental noise

The earlier simulations were generated from reference single-cell datasets without introduction of bias. But technical noise in scRNA-Seq and technical differences between scRNA-Seq and bulk RNA-Seq are both inevitable and may affect deconvolution accuracy. To further evaluate the robustness of cellGeometry, different types of noise were applied to simulated pseudo-bulk data derived from RA synovium scRNA-Seq^15^ (**Fig. 4a**): Gaussian noise, log noise, square root (sqrt) noise and shift noise (details in Methods). Importantly, sqrt noise is similar to the technical noise in scRNA-Seq replication experiments conducted by Brennecke *et al*^27^. Shift noise simulates the differences between sequencing chemistries, causing randomly selected whole genes to be detected at higher or lower abundance. Different standard deviations (SD) of noise were applied for each noise type (**Supplementary Fig. 18a**).

**Fig. 4.**
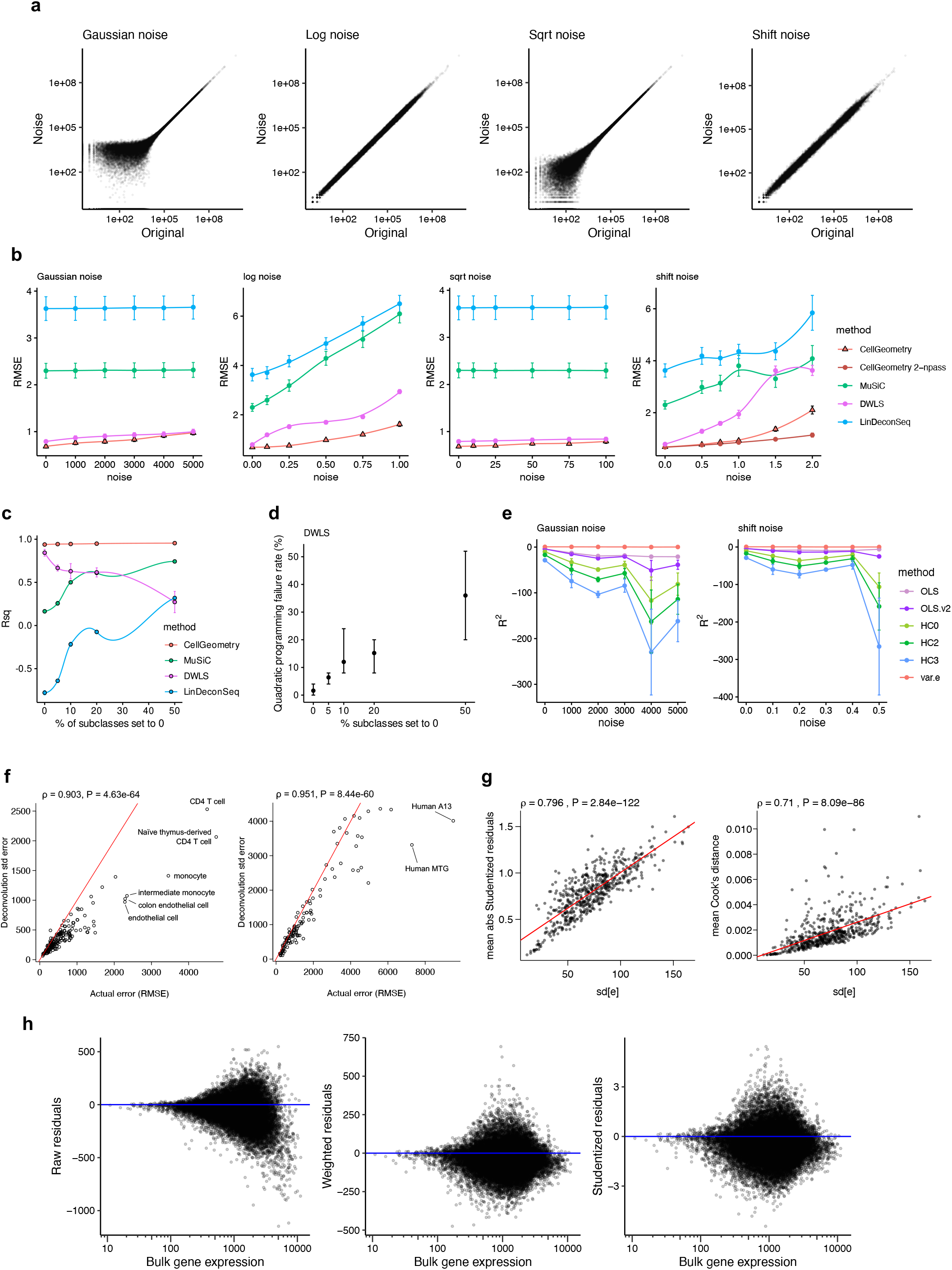
Non-negative geometric deconvolution (NGD) accurately predicts deconvolution error and exhibits improved handling of experimental noise. **a**, Scatter plots comparing gene counts from simulated pseudo-bulk data from rheumatoid arthritis synovium scRNA-Seq^15^ with the addition of different types of noise: simple addition of Gaussian noise to counts (Gaussian noise); counts are converted to log2+1 scale before the addition of Gaussian noise and counts are converted back to count scale (Log noise); counts are square rooted before addition of Gaussian noise and then transformed back to count scale (Sqrt noise); genes are randomly selected and multiplied by a random amount (shift noise). Sqrt noise resembles experimental noise observed by Brennecke et al^27^. Shift noise mimics systematic differences in sequencing chemistries between single-cell and bulk. **b**, Root mean square error (RMSE) of cell subclass percentages from the deconvolution of the simulated data from RA synovium with varying standard deviation (SD) of noise for each noise type, comparing cellGeometry, MuSiC, DWLS and LinDeconSeq. Five simulations (N = 25 samples) were tested. *x* axis represents the SD of Gaussian noise applied inside the relevant transformation. **c**, Mean coefficient of determination (R^2^) of cell subclass percentages from the deconvolution of the AMP simulated pseudo-bulk with varying proportions of cell subclasses that are randomly set to zero in each sample. Error bars represent the standard error of the mean (SEM). **d**, Plot of the quadratic programming failure rate in DWLS for **(c)**. Points show mean failure rate, whiskers show range. **e**, Mean coefficient of determination (R^2^) comparing deconvoluted cell frequency standard error and actual error (RMSE) when deconvoluting simulated pseudo-bulk data (N = 50 samples) from Tabula Sapiens^2^ with varying amounts of noise. Standard errors of cell counts were determined by analogy to ordinary least squares (OLS), OLS using non-negative modified compensation matrix (OLS.v2), heteroscedasticity-consistent SE (HC0, HC2, HC3) and row variance of residuals matrix per gene (var.e) [see Methods]. Error bars represent SEM. **f**, Scatter plot showing correlation between cellGeometry deconvolution cell frequency standard error and actual error (RMSE), when deconvoluting the simulated pseudo-bulk data from Tabula Sapiens (left) or the Human Brain Cell Atlas^3^ (right). P and ρ values were calculated using Spearman’s test. The red line shows the line of identity. **g**, Scatter plot showing that the square root of the row variance of the weighted residuals per gene (sd[e]) correlates with the mean Studentized residuals per gene (left) and mean Cook’s distance (right) following deconvolution of Cell Typist^18^ simulated data by cellGeometry. P and ρ values were calculated using Spearman’s test. **h**, Scatter plot of raw gene expression residuals, weighted residuals or Studentized weighted residuals following deconvolution of Cell Typist simulated pseudo-bulk data by cellGeometry, showing reduced heteroscedasticity following equal weighting of genes.

All methods were robust against standard Gaussian noise and sqrt noise but cellGeometry outperformed the other methods for log noise and shift noise (**Fig. 4b**; **Supplementary Fig. 18b**). Moreover, the completion rate of DWLS is adversely affected by log noise and shift noise due to the breakdown in the quadratic programming solution and resultant inability to deconvolute all simulated samples. DLWS failure rate ranged from 12-24% for log noise (SD=1) and 0-92% for shift noise (SD=2) (**Supplementary Fig. 18c**), mirroring similar failure rates with real bulk RNA samples.

### cellGeometry accuracy is less affected by absent cell types than other methods

The accuracy of cellGeometry was further assessed by having cell counts randomly set to zero for varying proportions of cell subclasses per sample prior to sampling to generate simulated pseudo-bulk data from RA synovium scRNA-seq^15^ (**Fig. 4c; Supplementary Fig. 19**). This mimics real life where some cell subclasses found in the scRNA-Seq may be completely absent from selected bulk samples. cellGeometry remained consistently accurate in the face of increasing proportions (0-50%) of subclass cell counts being set to zero. Both MuSiC, LinDeconSeq and DWLS performed less well compared to cellGeometry, and erroneously predicted the presence of certain cell subclasses despite the ground truth counting zero cells for those subclasses. The quadratic programming failure rate of DWLS increased proportionally with the rate of missing cell subclasses, suggesting that the underlying algorithm is sensitive to the absence of specific cell subclasses (**Fig. 4d**).

### Standard errors calculated using cross-sample variance of weighted residuals accurately predict deconvolution error

Due to the mathematical relationship between cellGeometry’s geometric algorithm and both ordinary least squares (OLS) and weighted least squares (WLS) (details in Methods), we extended the mathematical formulae for calculating standard error (SE) of regression coefficients for OLS and WLS to calculate the SE of the cell counts. The deconvolution must occur in count space as this is where gene expression and cells are both additive. However, this means that errors in gene expression typically break the assumption that modelling errors are homoscedastic. Thus we tested SE based on simple OLS (but adjusted for weighting), OLS using the modified inverse Gram matrix (OLS.v2), three variants of heteroscedasticity-consistent SE (HC0, HC2, HC3^28,29^), and a novel SE measure (var.e) based on the row (gene) variance of the residuals matrix across all bulk samples.

With large-scale simulated datasets (Tabula Sapiens^2^ and The Human Brain Cell Atlas^3^), the mean R^2^ comparing the actual error against the deconvoluted cell frequency SE using var.e was consistently higher than the other methods and is stable against noise (**Fig. 4e, Supplementary Fig. 20**). We observed strong correlation between the actual error and the deconvoluted SE for each cell subclass (**Fig. 4f**), showing that the var.e SE method reliably predicts deconvolution error (Tabula Sapiens ρ=0.903, CCC=0.668; The Human Brain Cell Atlas ρ=0.951, CCC=0.832). Logically, the cell subclasses with high predicted SE are those previously observed to have high similarity with another cell subclass (**Fig. 2b-c**).

With the accuracy of cell frequency estimates being partly subject to the genes defined in the gene signature, identification and removal of genes that behave inconsistently due to noise in either scRNA-Seq or bulk RNA-Seq may improve deconvolution accuracy. cellGeometry can compute Studentized residuals and Cook’s distance, as well as the var[e] measure. In large-scale scRNA-Seq datasets, the square root of var[e], i.e. sd[e], correlated strongly with both the mean absolute Studentized residuals per gene and Cook’s distance (**Fig. 4g**). Analysis of raw, weighted and Studentized gene residuals showed that the raw gene residuals exhibit marked heteroscedasticity as expected, but that the spherically scaled equal weighting of genes provides considerable correction of heteroscedasticity (**Fig. 4h**). This may explain why the heteroscedasticity-consistent SE methods HC0-3 were less accurate than OLS-based SE for determining deconvolution SE. Conversion of the weighted residuals to Studentized residuals confirms the improvement in heteroscedasticity, and standardises the scale of the residuals, allowing easier interpretation of whether certain genes are acting as true outliers.

### Extreme residuals flag problematic genes and yield biologically explainable reasons for deconvolution failure

With the ability to identify and remove outlying genes, cellGeometry can undergo multiple passes of deconvolution where genes with extreme residuals are removed with each iteration. This is demonstrated when deconvoluting real-world whole blood samples from RA patients in the PEAC study^30^ using the Cell Typist blood dataset^18^ (**Fig. 5a, Supplementary Fig. 21**). With 4 passes of deconvolution, there is detection of outlying genes at each pass and the range of studentized residuals decreases with the removal of these genes. 13 outlying genes were originally incorporated into the gene signatures (**Fig. 5b**). These genes are markers of granulocytes and red blood cells, which are cell types missing from Cell Typist blood^18^ as it only contains peripheral blood mononuclear cells (PBMC). Removal of these strongly genes decreased the deconvolution SE as well as increasing the compensation amount for each cell subclass (**Fig. 5c**). Therefore, missingness in cell type information in the reference scRNA-Seq can lead to major distortions of estimated cell type proportions.

**Fig. 5.**
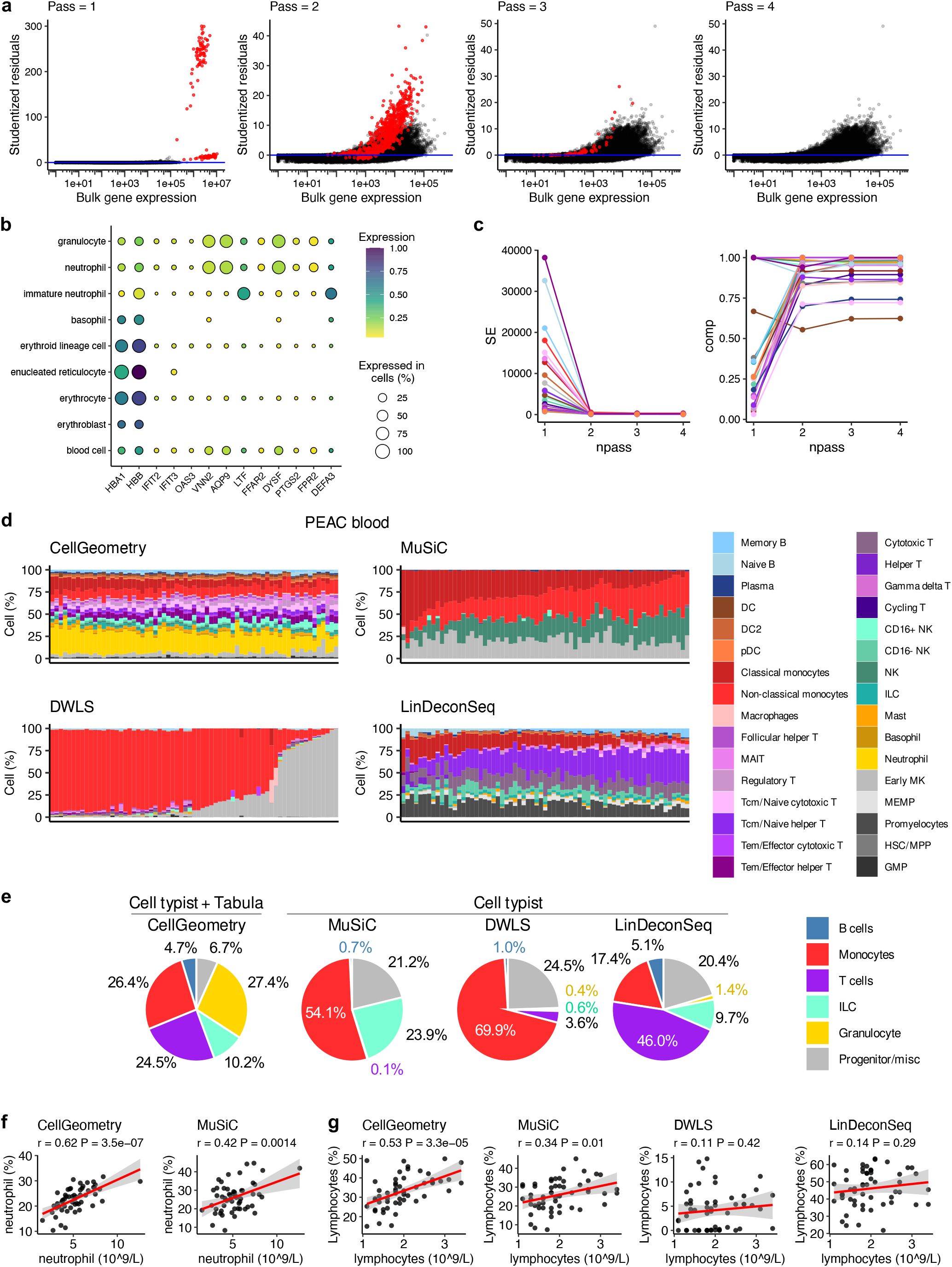
Extreme residual-based detection and removal of outlying genes along with neutrophil signature incorporation improves deconvolution of real whole blood RNA-Seq. **a**, Scatter plots of bulk gene expression and Studentized residuals against the number of deconvolution passes by cellGeometry. With each pass, outlying genes with extreme outlying residuals (red) are removed. Whole blood RNA-Seq from RA patients (n=67) in the PEAC study^30^ was deconvoluted with the Cell Typist blood^18^ dataset. Each point represents the residual gene expression error for an individual gene per bulk sample (black, with outliers in red). Beyond 4 passes, the algorithm automatically stopped as no further outliers were detected. **b**, Outlying genes that were removed from the gene signature after 4 passes of deconvolution were datamined in CZ CELLxGENE Discoverer. Bubble plots show scaled gene expression and percent of expressed cells in granulocytes and red blood cells. **c**, Standard error (SE) of cellGeometry deconvoluted cell counts (var.e method) and compensation values across cell subclasses improve following sequential deconvolution passes to remove outlying genes with extreme residuals. **d**, Stacked barplots of cell subclass proportions predicted by cellGeometry, MuSiC, DWLS and LinDeconSeq in whole blood RNA-Seq from RA patients in the PEAC study. cellGeometry deconvoluted with the Cell Typist blood dataset merged with basophils and neutrophils from Tabula Sapiens blood^2^. Other deconvolution methods used the Cell Typist blood dataset alone. **e**, Pie charts of the mean blood cell type group proportions predicted by cellGeometry, MuSiC, DWLS and LinDeconSeq in whole blood RNA-Seq from RA patients in the PEAC study. **f, g**, Scatter plots comparing deconvoluted cell percentages and Coulter counter measurements for **f**, neutrophils and **g**, lymphocytes (B cells, T cells and ILC) from RA blood samples. P value and r coefficient were calculated using Pearson’s correlation.

It is informative that the genes flagged as deconvolution outliers based on residual gene expression included interferon response genes *IFIT2, IFIT3* and *OAS2*. Although these genes were selected during signature matrix creation as being cell-specific, interferon response genes are highly labile, and their expression increases several-fold in response to interferon stimulation. Since the single-cell data was generated in healthy individuals, while the bulk blood data was generated in actively inflamed RA patients who are known to frequently show a type I interferon response blood signature^31^, this suggests that residual gene expression left over after cellGeometry deconvolution could be biologically relevant and this represents a useful new way to extract residual gene expression that takes into account differences in cell proportions.

### Accurate deconvolution of whole blood bulk RNA-Seq

Using cellGeometry’s unique ability to merge gene signatures from different single-cell datasets, the Cell Typist PBMC dataset^18^ gene signature was merged with single-cell signatures for basophils and neutrophils from Tabula Sapiens peripheral blood^2^ (**Supplementary Fig. 22**) using quantile-quantile mapping of the signature matrices to adjust for differences in the overall level of gene expression (**Supplementary Fig. 23**). Tabula Sapiens blood alone is less useful because it does not have the immune cell subset resolution akin to Cell Typist blood which has extensive B and T cell subsets. cellGeometry was benchmarked with the previously assessed deconvolution methods. The other deconvolution methods do not have the ability to merge gene signatures and so were deconvoluted with Cell Typist blood dataset alone.

CellGeometry deconvoluted whole blood RNA-Seq data using the merged blood reference dataset (**Fig. 5d-e, Supplementary Table 2**) and its relative neutrophil and lymphocyte percentages correlated well with Coulter counter measurements (**Fig. 5f-g**). In contrast, other deconvolution methods show marked biases in the cellular proportions to particular cell subclasses (**Fig. 5d-e, Supplementary Table 2**) and the lymphocyte percentages did not correlate as well with Coulter counter measurements compared to cellGeometry (**Fig. 5g**). MuSiC identified excess proportions of monocytes, natural killer (NK) cells and megakaryocytes (**Fig. 5d**). However, MuSiC reported zero proportions for 13 subclasses, including multiple B/T cell subsets (**Supplementary Table 2**), much like the simulated data where MuSiC had poor accuracy with T cell subsets. T cells make up ~16% (reference range 6-31%) of white cells in whole blood^32^. Despite DWLS performing well with simulated data, DWLS identified excess proportions of monocytes and megakaryocytes in whole blood and showed minimal detection of B, T and NK cell subsets (**Fig. 5d-e**). Conversely, LinDeconSeq showed less extreme bias compared to MuSiC and DWLS, but was dominated by Tcm/Naive helper and cytotoxic T cells. It also detected a strangely high proportion of promyelocytes, which had only 4 cells in the single-cell reference. Overall, LinDeconSeq was unable to detect 9 subclasses (**Supplementary Table 2**). We also tried re-running MuSiC, LinDeconSeq and DWLS whilst removing single-cell clusters with <10 cells in the reference. As a whole, this hardly altered their deconvolution results.

In addition, we have tested the effect of manually removing the same outlying genes identified by cellGeometry from the gene signatures of the other deconvolution methods to see if this improved their functionality, but this led to only slight improvements in their deconvolution accuracy.

MuSiC was also investigated for deconvolution using Tabula Sapiens blood subclasses alone, but it failed to detect 10 B/T-cell and myeloid cell subclasses (**Supplementary Fig. 24, Supplementary Table 3**). The neutrophil percentage predicted by MuSiC showed considerably weaker correlation with Coulter counter measurements than CellGeometry (**Fig. 5f**).

### Validation of cellGeometry deconvolution against synovial immunohistology

Bulk RNA-Seq of 183 synovial biopsies from RA individuals from the R4RA randomised clinical trial^33^ were deconvoluted using the AMP RA synovium single-cell dataset^15^, which contains 18 cellular subclasses including 4 fibroblast, 4 monocyte, 4 B cell and 6 T cell subsets (**Fig. 6, Supplementary Table 4**). Problematically, DWLS only managed to deconvolute 36/183 samples (80.3% failure rate), reportedly due to failure of the quadratic programming solution (**Supplementary Fig. 25a**).

**Fig. 6.**
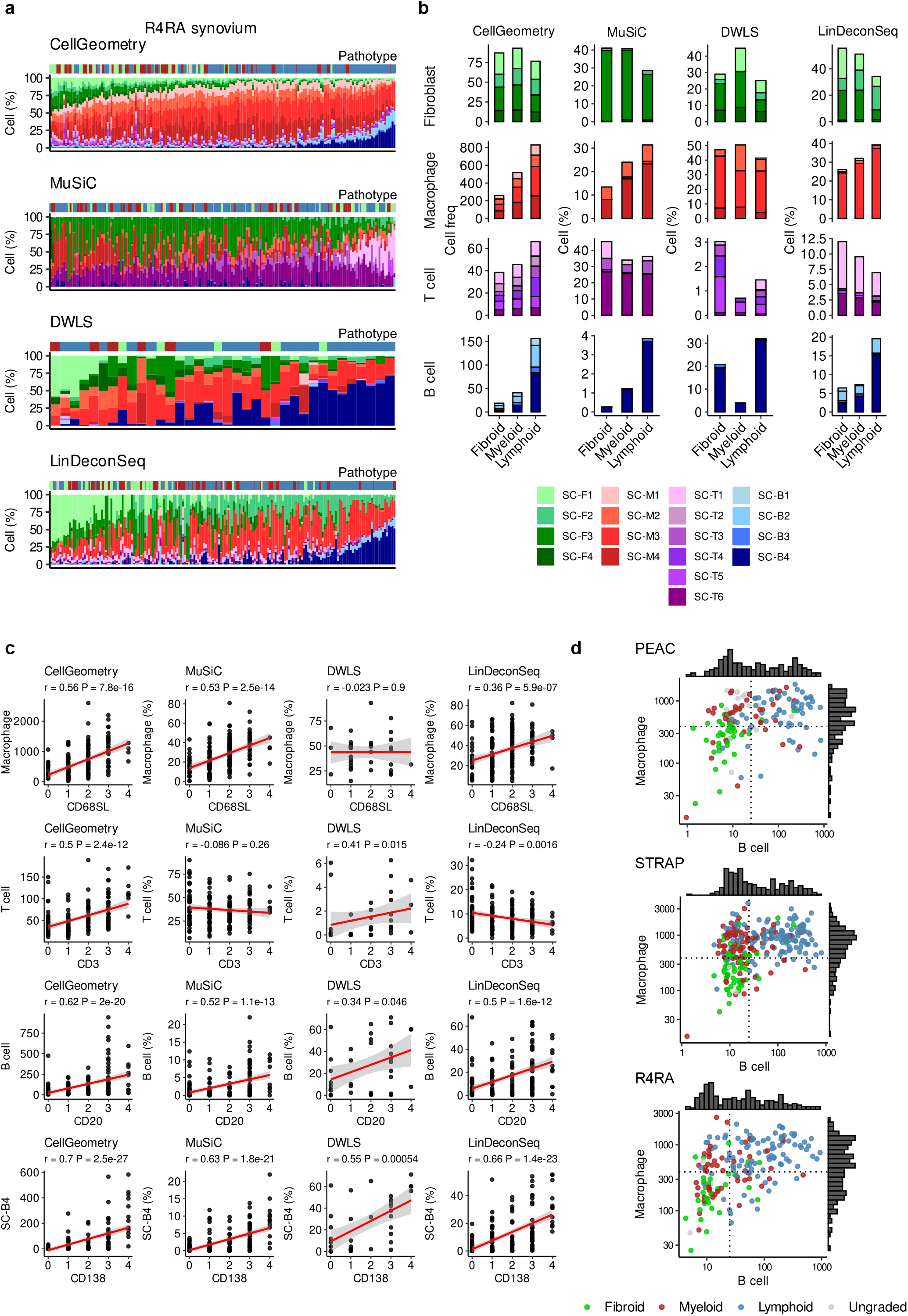
Cellular deconvolution of rheumatoid arthritis synovial bulk RNA-Seq improves upon immunohistology classification of tissue endotypes. **a**, Stacked barplots of rheumatoid arthritis (RA) synovium cell subclass proportions in bulk RNA-Seq of synovial biopsies from RA patients from the R4RA trial^33^ as deconvoluted by cellGeometry, MuSiC, DWLS and LinDeconSeq using synovium scRNA-Seq^15^ as reference. Each stacked column corresponds to an individual synovial biopsy. The coloured track above shows the corresponding histology-based pathotype class (green = fibroid, red = myeloid, blue = lymphoid, grey = ungraded). **b**, Stacked barplots of the average relative cell subclass abundance or proportion for each cell group (fibroblast, macrophage, T cell, B cell) with respect to histology-based pathotype (fibroid, myeloid and lymphoid). Average relative cell subclass abundance was used for cellGeometry whereas average cell subclass proportion was used for MuSiC, DWLS and LinDeconSeq. **c**, Correlation between the different cell type (Macrophage, T cell, B cell) or SC-B4 (plasmablast) levels with their respective histology cell surface markers (CD68SL, CD3, CD20 and CD138). Relative cell subclass abundance was used for cellGeometry whereas cell subclass proportion was used for MuSiC, DWLS and LinDeconSeq. P value and r coefficient were calculated according to Spearman’s correlation test. **d**, Deconvolution of RA synovium from the PEAC^30^, STRAP^37^ and R4RA^33^ trials were undertaken by cellGeometry using the AMP phase 1 RA synovium scRNA-Seq dataset. Scatter plot with overlaid histograms of B cell and macrophage abundance with points coloured by the original histology-based pathotype (Green = fibroid, red = myeloid, blue = lymphoid, grey = ungraded). B cell and macrophage low/high status were defined using thresholds of ≥25 and ≥388 cells respectively.

Previous studies have demonstrated cellular composition heterogeneity in synovium in RA patients and have identified endotypes based on histopathology and gene expression data^30,34,35^. In the R4RA study, biopsies were classified histologically into three pathotypes (pathological endotypes): lymphoid (high lymphocyte and myeloid cells), myeloid (macrophage rich) or fibroid (low immune cells), based on immunohistology stains for CD3 (T cells), CD20 (B cells), CD68 (synovial lining and sub-lining macrophages) and CD138 (plasma cells). With histology-based definitions, there may be discrepancies between the thin tissue sections used for staining and the pooled biopsies used for transcriptomic analysis. This is highlighted by correlating histology markers with known cell-specific gene markers: *MS4A1, MS4A7* and *CD3E* for B cell, macrophage and T cell, respectively (**Supplementary Fig. 25b**). Generally, there is a significant positive correlation but there are instances when there is low staining but high expression of the cell-specific gene marker, suggesting the presence of these cells in the RNA sample. In spite of the use of up to 3 sections at progressively deeper cutting levels^36^, 3µm tissue histology sections will be less representative of the tissue as whole compared to the typical 6 needle biopsies pooled for an RNA sample.

Accepting these limitations, the immunohistology of the biopsies provides a validation of the deconvolution as an independent method for semi-quantitative measurement of cell abundance in tissue samples. cellGeometry detected all 18 subclasses in the synovium. There were generally high B cell levels in lymphoid biopsies and high macrophage levels in lymphoid and myeloid biopsies while low immune cell levels were observed in fibroid biopsies (**Fig. 6a-b, Supplementary Fig 25c**).

However, other deconvolution methods were dominated by specific subclasses across all samples (**Supplementary Table 4**) and did not completely reflect the pathotype classification (**Fig 6b**). MuSiC detected a higher proportion of SC-F4 and T cell subclasses, particularly SC-T6, but inappropriately low B cell subclass proportions. Although MuSiC predicted high levels of B cells in lymphoid, its output was swamped by plasmablasts (SC-B4), which dominated the total B cells across the pathotypes (**Fig. 6b**). Also, DWLS identified a minimal proportion of T cell subclasses, and the fibroid synovium, which should have low immune cell infiltration, had similar macrophage proportions to myeloid and lymphoid synovium. LinDeconSeq was unable to detect SC-T2 and SC-T4.

cellGeometry’s realistic deconvolution output is also demonstrated through the significant positive correlations of the relative abundance of macrophage, T cell, B cell and plasmablasts (SC-B4) with their respective histology markers (**Fig. 6c**). Conversely, MuSiC and LinDeconSeq deconvolution estimates of T cells failed to correlate with CD3 histology and DWLS macrophage estimates failed to correlate with CD68.

### cellGeometry redefines pathotypes to delineate the cellular heterogeneity of rheumatoid arthritis synovium

Using histological pathotypes as reference, we sought to redefine the RA synovium pathotypes according to the estimated abundance of synovial B cells and macrophages from deconvolution of bulk RNA-Seq using cellGeometry (**Fig. 6d**). As well as the R4RA trial (n=183 biopsies)^33^, deconvolution was performed on RA synovial biopsies from the PEAC study (n=160)^30^ and the STRAP randomised controlled trial (n=273)^37^. Decision tree-based prediction of histological pathotypes from cellGeometry estimated B cell and macrophage cell counts was used to define thresholds. Final thresholds were derived by averaging thresholds from repeated subsampling of the data (10 folds with one fold left out each time, 15 repeats) (**Supplementary Fig. 26a**). This was also used to assess stability of thresholds. Based on deconvolution estimates, the lymphoid pathotype was redefined as estimated B cells ≥25 cells; myeloid pathotype was redefined as low B cells (<25), but macrophages ≥388 cells; and fibroid was redefined as low B cell and macrophage status. These redefined pathotype groupings consistently reflect the original histology-based pathotype across all studies (**Supplementary Fig. 26b-c**).

The redefined pathotypes were more precise in terms of relative cell subclass abundances and proportions compared to the original histology-based pathotype (**Supplementary Fig. 27a-c**). With the redefined myeloid pathotype, the relative B/T cell levels decreased whilst the relative macrophage levels increased compared to the original pathotype (**Supplementary Fig. 27b**). Furthermore, the pathotype redefinition highlights the differences in cellularity to a greater extent, where fibroid samples have a low number of total cells per biopsy whilst myeloid and lymphoid samples have a higher number of cells due to immune infiltration (**Supplementary Fig. 27d**). As a result of the refined cellularity estimates, the high fibroblast percentage in the fibroid pathotype is starker and clearer with the redefined pathotypes than with the original histology-based pathotypes (**Supplementary Fig. 27c**). Thus, deconvolution of bulk RNA-Seq using cellGeometry produced enhanced classification of RA synovium pathotypes across three sizeable cohorts compared to histology. This is important for future downstream applications such as prediction of clinical response to therapy.

## DISCUSSION

Although previous reference-based deconvolution methods typically perform well with simulated pseudo-bulk data, these methods commonly fail to deconvolute real-world bulk RNA-Seq data, producing unrealistic, nonsensical cell abundances, or fail outright with a large percentage of samples. Existing methods are slow or incapable of handling large reference datasets and not fully compatible with modern matrix systems such as anndata and DelayedArray^22^. cellGeometry performs ultra-fast deconvolution of bulk RNA-Seq from a user-provided reference scRNA-Seq dataset. The speed gain derives from the extensive use of large matrix operations which are data type optimised: DelayedArray subroutines are used for sparse single-cell count matrices; while R’s matrix cross-product, which implements BLAS subroutines, is used for deconvolution itself.

The underlying method implemented by cellGeometry, which we term non-negative geometric deconvolution (NGD), has an important advantage over other methods. Commonly, scRNA-Seq and bulk RNA-Seq differ in chemistry. scRNA-Seq is often 3’ or 5’ biased, whereas bulk RNA-Seq may cover the full length of transcripts. Depletion of superabundant transcripts (e.g. globin or ribosomal genes) is another source of transcriptome variability. This means there may be systematic (but unknown and unpredictable) differences in gene expression between single-cell and bulk. Thus, cell-specific genes may be expressed at consistently higher or lower levels in bulk than anticipated from the scRNA-Seq signature. In NGD, variable compensation preserves the relative relationship, so that increased expression of cell-specific genes corresponds to linearly increased cell type abundance. Other methods ignore systematic errors and may yield negative cell quantities or fail to generate a solution. cellGeometry has been tested for robustness in multiple real-world deconvolution situations and can handle large-scale datasets (>3 million cells). Outlying genes with extreme residuals can optionally be removed from the gene signature to increase the deconvolution accuracy. This may also flag problems with the single-cell reference.

A limitation of our study is that it lacks true cross-platform studies performed on matching specimens. These would improve our understanding of the biases introduced by differences between bulk and single-cell sequencing chemistries, which could lead to more accurate deconvolution algorithms in future. Similarly, single-nucleus RNA-Seq appears to have advantages over single-cell RNA-Seq for certain tissues, notably brain, where fragile or elongated cells such as neurons or glia are damaged or activated by disaggregation techniques^38^. Simulations suggest that snRNA-Seq yielded more reliable deconvolution of bulk brain samples than scRNA-Seq^39^. However, this study lacked experimentally confirmed ground truth. To truly determine whether snRNA-Seq or scRNA-Seq is optimal for deconvolution of real bulk brain samples would require direct comparison of these methods alongside bulk RNA-Seq on matching samples.

The modular aspect of cellGeometry’s workflow enables diagnostics of the gene signatures, which is critical, since deconvolution accuracy is dependent on accurately defined cell type-specific reference gene signatures. Similarly named cell clusters may be redundant, as seen in Tabula Sapiens^2^, so deconvolution can be improved if overlapping subclasses are removed. Another unique feature of cellGeometry is the ability to merge two scRNA-Seq signatures, as demonstrated through deconvolution of the Human Brain Cell Atlas^3^ which contains neurons and non-neuronal cells in separate datasets. **Supplementary Table 5** comprehensively compares the different features between deconvolution methods.

Applied to RA synovium^30,33,37^, cellGeometry captures the cellular composition of bulk samples with its relative cell type abundance significantly correlating with cell quantification by immunohistology. Therefore, cellGeometry facilitates enhanced pathotype classification of the synovium based on RNA-Seq for downstream analysis, which may potentially be useful for inferring cell type-specific mechanisms behind RA progression and drug response.

Deconvolution of whole blood RNA-Seq showed that deconvolution may be hampered by missing cell types in the reference single-cell data. Currently, there is no single scRNA-Seq dataset which covers all blood cell types and immune cell subsets. This is due to technological limitations, such as cells being lost during cryopreservation which is particularly the case for neutrophils^40,41^. Irrespective, by merging the Cell Typist blood signature with neutrophil and basophil signatures from Tabula Sapiens, cellGeometry produced reliable estimates of cell proportions in whole blood RNA.

In summary, cellGeometry takes full advantage of large-scale scRNA-Seq datasets to efficiently deconvolute complex cell abundances in bulk samples. By harnessing annotated single-cell atlases, cellGeometry bridges the gap between single-cell and bulk RNA studies to illuminate cell type specific-mechanisms in disease and health.

## METHODS

### Implementation

cellGeometry is implemented as an R package which can be installed from CRAN (https://cran.r-project.org/package=cellGeometry). Two comprehensive vignettes, which include full instructions for installation and deconvolution of peripheral blood using the Cell Typist scRNA-Seq dataset and simulated brain pseudo-bulk using the Human Brain Atlas datasets, are included with links on the CRAN webpage. Atlas. Source code is available on Github at https://github.com/myles-lewis/cellGeometry.

Users provide the following input data: i) the count matrix of a single-cell RNA-Seq dataset, along with ii) predefined cell clusters, and iii) the count matrix of bulk RNA-Seq samples to be deconvoluted. The package includes functions to generate simulated pseudo-bulk counts from the scRNA-Seq data to simulate deconvolution.

### Angle method for identification of cell-specific genes

Single-cell gene expression is first transformed to be less skewed, by a log_2_ +1 transformation and the mean expression per cell calculated for each single cell cluster. We consider a high dimensional space consisting of dimensions for each cell cluster or subtype. During the gene selection process, we initially assume that each cell subtype is mutually distinct to search for perfectly cell-specific gene markers. Each gene represents a point in space with cell types as dimensions in *n* dimensional space for *n* cell subclasses, ℝ^n^, specifically the non-negative orthant. We consider the hypothetical perfect gene marker to be only expressed in one cell type and to have zero expression in all other cell types. A classic example is *FOXP3* which is only expressed in regulatory T cells and shows zero (or near zero) expression in all other cells. Genes which are specific for a particular cell type lie closest to the dimensional axis for that cell type and can be ranked by their vector angle.

If each gene vector is scaled to the unit hypersphere, the angle between that gene’s vector and the axis corresponding to each cell cluster is given by arc cosine of the magnitude of the scaled expression along that axis. In matrix terms, if *G* is an *m* × *n* nonnegative matrix of mean gene expression on log_2_ +1 scale with *m* genes in rows and *n* cell clusters or subclasses in columns,

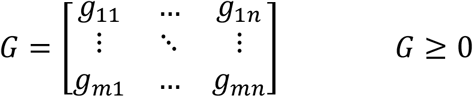

we think of each gene as a vector *G*_*i*,∗_ derived from each row of matrix *G*, with cell subclasses as dimensions and elements *g*_*ij*_ denoted by

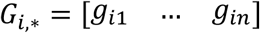

Then we can define matrix *H* with elements *h*_*ij*_ whose elements have been scaled by the Euclidean norm for the vector for each gene *i*

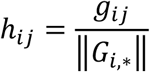

or in matrix terms

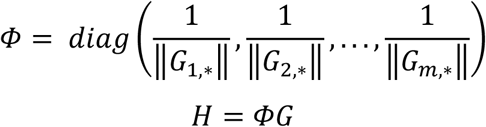

Applying arc cosine to *H* yields a matrix containing angle values between every gene *i* and every cell cluster *j*.

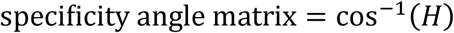

The most cell-specific markers approach an angle of zero for that cell cluster with a perfect marker achieving an angle of zero for that cell type and an angle of 90° for all other cell types. This allows genes to be easily ranked from lowest angle to highest based on their cell cluster vector angle.

### Vector angle specificity plot

We developed a novel visualisation to compare cell type-specificity of all genes for a particular cell type. In these plots, the specificity of each gene is measured as the cell specificity vector angle is shown on the *x* axis, with small angles representing greater specificity, while mean gene expression is shown on the *y* axis, since genes with higher expression are generally more reliable in the real world. Genes with perfect specificity, i.e. exclusively expressed in one cell cluster alone, have a vector angle of zero. Points are coloured to show ranking of each gene by mean gene expression across different subclasses, so that genes which are optimal markers are usually ranked first.

### Optimal signature matrix mean gene expression per cell

Log_2_ transformation of the data ensures that the mean gene expression is considerably less sensitive to outlying cells with very high counts which skew the mean. It should be noted that although we recommend the application of log_2_ to the data, for simulations, cells for each cell subclass are randomly sampled from the original scRNA-Seq dataset and gene expression in counts is summed to generate the pseudo-bulk counts. Therefore the mean gene expression per cell across a particular cell subclass for simulated data will converge on the arithmetic mean for that subclass and not the geometric mean. cellGeometry can calculate both mean of log_2_(counts+1) as well as the arithmetic mean, so that deconvolution of simulated pseudo-bulk can be optimally achieved using the arithmetic mean of counts as the gene signature matrix. Users can use either option, or a third method, the trimmed arithmetic mean of counts, which is also provided.

### Expression filtering

One problem which can occur is that some low expressed genes might only be expressed in a particular cell cluster. So, a low expression filter is applied to remove genes whose maximum mean log_2_ expression falls below the level of the filter. Another problem which often occurs is that due to the amplification applied to single cell RNA, highly expressed genes sometimes exhibit low level expression in other cell types which might be due to contamination. This low-level expression of the gene in other cell types, which can be thought of as noise, tends to shift the angle of that gene away from zero for the main cell cluster for which the gene is most specific. Thus, an optional noise filter can be applied which reduces low level gene expression in non-specific cell types. Gene expression below the level of the filter in low expressing non-specific cell types is reduced to zero as a form of noise reduction. Empirically we observed that to be effective the noise threshold needed to be proportional to the maximum log gene expression for each gene. Thus, we use a simple ratio with a typical value of 0.1 to 0.25 to determine the threshold for shrinking noisy low-level expression for each gene to zero. In simulations, if the arithmetic mean is used, the gene signature is already perfect, so the noise filter is only recommended for use with real bulk samples.

### Deconvolution by vector projection with variable compensation

To deconvolute bulk RNA-Seq, we restrict that dataset to only gene expression of the cell-specific subset of genes in each sample, using raw counts as input. We visualise a single bulk RNA-Seq sample as a vector in high dimensional space, this time with *genes* as dimensions in *m*-dimensional vector space ℝ^m^, specifically the non-negative orthant. Each cell cluster is represented as its own vector of mean gene expression per cell. Thus, to determine the quantity of cells for each cell cluster, gene expression from the sample of interest is projected along the vectors for each cell cluster using the vector dot product to quantify the scalar amount of projection. The scalar projection of vector **a** in the direction of **b** is:

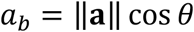

where *θ* is the angle between vectors **a** and **b**. This can be rewritten:

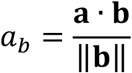

The quantity of cells in the bulk RNA-Seq is related to *a*_*b*_ namely the projection of the bulk gene expression vector along the vector **b** representing the vector of mean gene expression for the specific cell cluster in *m*-dimensional vector space ℝ^m^, specifically the non-negative orthant. To convert *a*_*b*_ to an amount of cells rather than in terms of the length of **b**, we need to redefine **b**, the signature vector as mean gene expression per cell for a cell cluster, to have length 1 to represent 1 cell. Thus the number of cells is:

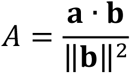

This formula calculates the projected cell abundance for a single bulk sample and single cell cluster. But the same process can easily be computed using matrices instead of vectors. Thus we start with a nonnegative matrix of bulk RNA-Seq counts *Y* with *m* genes in rows and *k* samples in columns matching the gene signature matrix *G* with genes in *m* rows and clusters in *n* columns, also on a count scale.

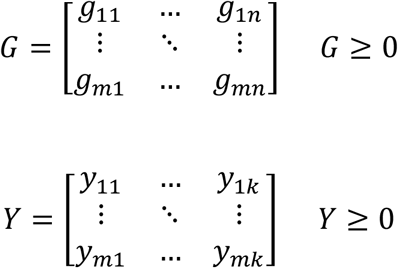

Here the mean gene expression in the gene signature matrix *G* has been converted back to count space. The algorithm allows for different methods of mean gene expression to be calculated. While using gene means calculated from mean of log_2_(counts + 1) as described above works best for identifying the best gene markers, for deconvoluting simulated bulk data, the arithmetic mean of raw (unlogged) counts is optimal (unbiased).

We first scale the columns of *G* by dividing by each cell cluster column by ‖*G*_∗,*j*_‖^2^ where *G*_∗,*j*_ are the vectors for each cell cluster *j* represented by columns from *G* for *j* = 1 to *n*,

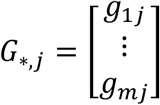

to create column scaled matrix *G’*.

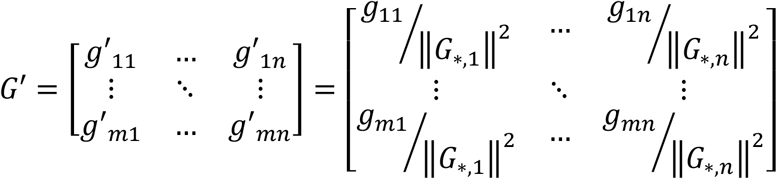

Or in matrix terms

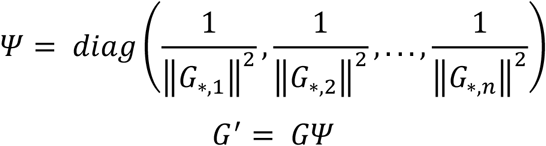

The quantitation in terms of numbers of cells is:

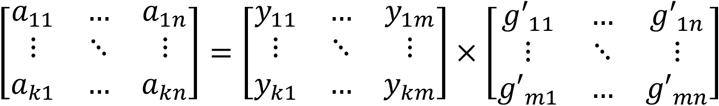

Or in matrix notation

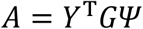

which yields a matrix of cell numbers *A* with *k* rows for samples and *n* columns for cell clusters. The transformation can be performed either in (unlogged) count space or in log_2_(counts +1) space.

### Equal weighting of genes by unit hypersphere scaling

These first equations leave the relationship between signature genes untouched. However, this means that the most highly expressed genes dominate the deconvolution of each subclass. A modification on this is to scale each of the signature genes to the unit hypersphere so that the vector for each gene has length = 1. This is the same scaling vector as was used for the specificity angle calculation. To preserve the relationship between genes in the bulk RNA-Seq to be deconvoluted, each gene in the bulk matrix is scaled accordingly to align with the amount of scaling applied to the gene signature matrix. The scaling vector **e** is calculated as:

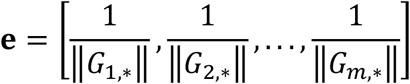

and note that

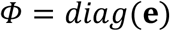

where *G*_*i*,∗_ are vectors for each gene represented by rows from *G* for *i* = 1 to *m*. The rows of both *G* and *Y* are scaled by each element of **e**, yielding *H* and *Y*_*e*_ respectively as shown below. The geometric interpretation of this is that both *H* and *Y*_*e*_ have been transformed into a new gene space in which the vectors for each gene in *H* lie on the unit hypersphere in cell subclass space, while crucially the interrelationships between genes in *H* and *Y*_*e*_ have been preserved. Thus each signature gene in *H* is given equal weighting for each cell subclass vector for the cell subclass deconvolution. Formally, *H*_∗,*j*_ represents the columns of *H*, such that the algorithm can be summarised as

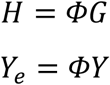

In order to calculate the weighted deconvolution, we substitute *H* for *G* in the earlier equations.

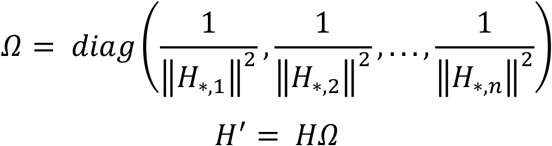

Therefore the equal gene weighted deconvolution becomes

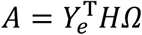

### Correcting spillover with a compensation matrix

However, as the basis vectors for each cell cluster are not completely specific, i.e. non-orthogonal, they show spillover from one cell cluster vector into another. The spillover matrix can be calculated from the gene signature projection onto itself, where *G*^*T*^ is the transpose of *G*, and, as before, the vector upon which we project is normalised to unit length to reflect mean expression in a single cell from that cell type.

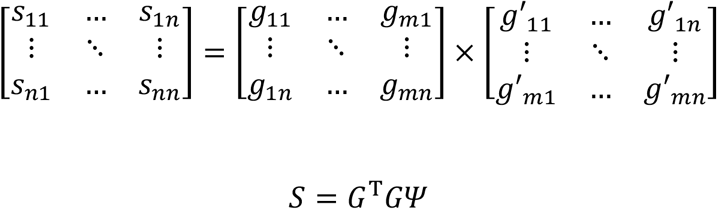

or for the equally weighted version

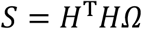

As a spillover matrix, *S* is a positive square *n* × *n* matrix whose diagonal values are 1 and whose other values represent the spillover of one cell cluster’s vector as projected into another. A perfect gene signature with perfect markers in every cell cluster would have zero spillover and would thus be the identity matrix. The smaller the vector angle for each gene selected as specific for a particular cell type, the more orthogonal that cell type vector will be to other cell type vectors during deconvolution. Thus, the vector angle method is designed to choose genes that minimise the spillover matrix and reduce the compensation burden.

To compensate for spillover, we use a similar system to flow cytometry where signal from one fluorochrome commonly bleeds into another. Compensation can be solved by determining the inverse matrix for the spillover matrix. *A*, the apparent cell quantities, will be affected by spillover unless the cell cluster vectors are perfectly specific (i.e. in vector terms they are all perfectly orthogonal). In reality, cell-specific genes are only very rarely 100% specific. If *T* is a matrix of true cell numbers for each cell cluster then:

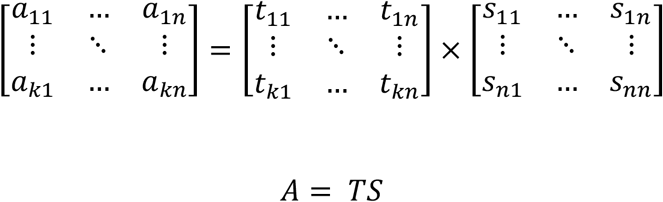

Where *S* is a square *n* × *n* matrix of spillover values. This can be solved as:

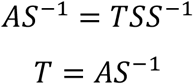

Where *S*^−1^ is the inverse matrix of *S* and can be understood to represent the compensation matrix as found in flow cytometry analysis. The spillover matrix can be visualised as a heatmap which can help to show the cell types whose cell cluster vectors bleed most into other cell types. This can be understood as cell clusters whose cell markers are not sufficiently specific to that cell cluster alone, with spillover representing the amount of expression in other cell types. Putting this all together we have:

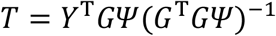

or with equal weighting’

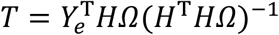

### Non-negative tuning of compensation

One problem encountered in real life is that full 100% compensation can lead to negative values in the quantified cell amounts. This is mostly due to systematic errors being amplified in the compensation matrix. We propose that the compensation for each cell type is varied from 1 → 0 to determine the threshold of the maximum amount of compensation which can be tolerated which avoids any of the outputted cell quantities becoming negative. This is solved by varying each element of a weight vector **w** between 1 and 0 until all the output results for that cell cluster are non-negative, by finding min(output)^2^ = 0. The optimisation method uses a combination of golden section search and successive parabolic interpolation.

To enable this, the compensation process is modified to include a vector of weight coefficients **w** for each cell cluster ranging from 0 to 1, which allows the amount of compensation for each cell type to range from 0 to 100%. In practice we solve *C’*, the weighted compensation matrix in the following manner:

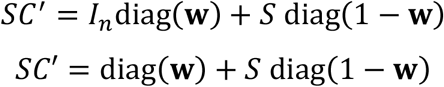

*I*_n_ is the identify matrix matching the length of **w**, so that columns of the right-hand side output matrix vary smoothly from identity matrix to the spillover matrix depending on the individual weights in **w**. This allows each compensation weight coefficient in **w** to be varied and affect only one cell cluster independently without affecting the results of the other cell clusters. Thus:

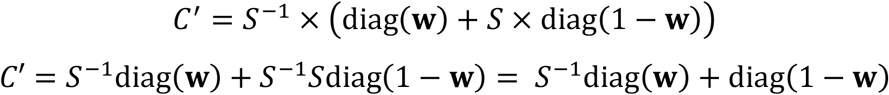

So the final true cell abundance *T* with samples in *k* rows and cell clusters in *n* columns is:

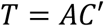

In cellGeometry, the compensation weightings for each subclass, i.e. each element of **w**, can be set manually, or the compensation weightings can be automatically tuned, on the basis that we know that the final true cell abundance must be always positive or zero and can never be negative. So in *p* subclasses where, if full compensation is applied this leads to *T*_∗,*p*_ for that subclass yielding negative cell counts for that cell cluster, then we solve these selected *p* elements of **w** so that *T*_∗,*p*_ is non-negative.

Formally, if *T*_∗,*j*_ are the columns of *T* with *j* = 1 to *n*, for any *p* in *j* where min(*T*_∗,*p*_) < 0, the *p*th element of **w**, *w*_*p*_ is solved for min(*T*_∗,*p*_) = 0 by varying *w*_*p*_ within the bounds 0 to 1 and finding *w*_*p*_ which yields the minimum of min(*T*_∗,*p*_)^2^.

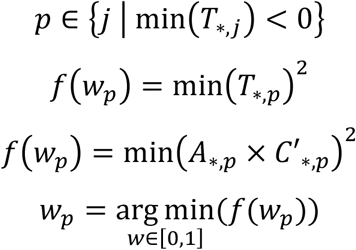

If *w*_*p*_ = 0, this guarantees that column *T*_∗,*p*_ is non-negative as the associated compensation matrix column *C*′_∗,*p*_ becomes the *p*th column of the identity matrix and no compensation occurs for that column, hence *T*_∗,*p*_ = *A*_∗,*p*_ and all values in *A* are > 0. Thus there always exists values of *w*_*p*_ which ensure non-negativity of *T*. cellGeometry checks *T* for the presence of negative cell counts and warns users if this is detected. If negative cell counts are detected in specific cell clusters, then the compensation weighting for each cell cluster in turn is automatically determined to prevent negative cell counts in that cell cluster, unless users override this and set compensation weightings manually.

Examples of the coefficient paths with varying **w** are shown in **Supplementary Fig. 4**. A critical aspect of the method is that each element of **w** controls each cell subclass independently, and that all bulk samples are considered simultaneously. This means that proportionality between the signature gene expression and deconvoluted cell counts for a particular cell subclass is always preserved (but at the expense of reducing spillover compensation).

### Compact algorithm

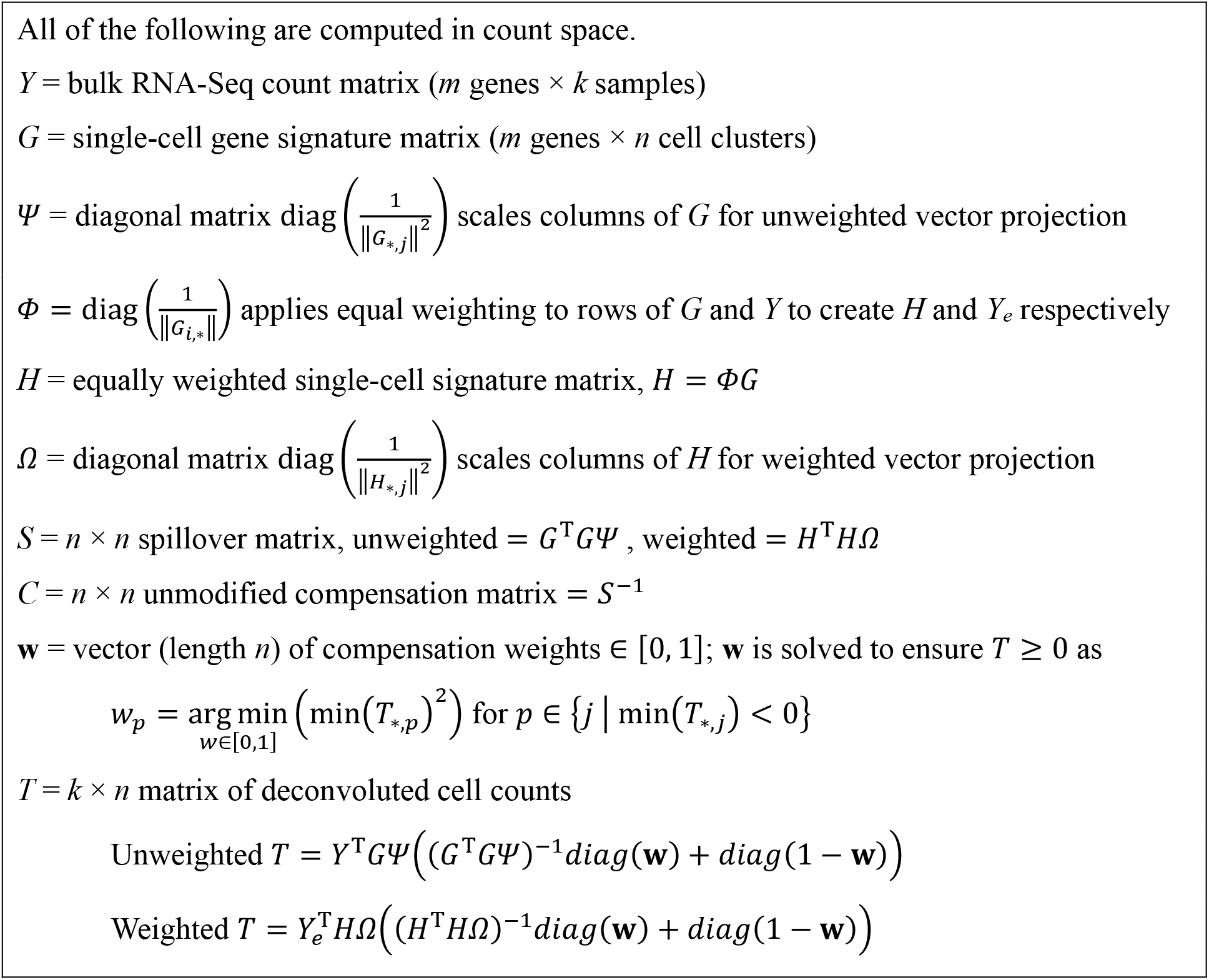

### Fast matrix operations and parallelisation

Note that thus far almost all operations can be performed as matrix operations and thus the deconvolution of large numbers of samples against large numbers of cell clusters can be performed in a single series of matrix manipulations, making extensive use of R’s matrix cross-product, so that the overall deconvolution computation is extremely fast. The only exception is the automatic compensation weighting, which is performed on each cell cluster in turn. However, this only needs to be performed on selected cell clusters that require it. In fact the slowest part of the method remains the calculation of averaged gene expression in each cell cluster due to the imposition of handling very large sparse matrices containing the raw count data for individual cells, optimised through the use of sparse matrix row functions and DelayedArray matrix functions for the largest scRNA-Seq datasets, as well as fork-based parallelisation.

### Relationship to ordinary least squares

The geometric solution to deconvolution described above has a partial relationship to ordinary least squares (OLS). The matrix notation description of a general multivariate linear model is:

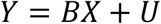

The maximum likelihood solution for the matrix of model coefficients *B* is:

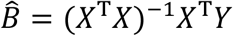

We can consider *X* to be the gene signature matrix, previously referred to as *G* with *m* genes as samples and *n* cell types as regressors. Then *Y* is a matrix of the gene expression for bulk RNA-Seq samples whose expression is to be deconvoluted with samples in columns and genes in rows. Then *B* is a matrix of model coefficients representing the quantity of each cell type which is to be determined in this single sample of bulk RNA-Seq. However, the two main differences are that:

i. the model is a non-negative regression, i.e. *Y* ≥ 0, *X* ≥ 0 and *B* ≥ 0 and
ii. we deliberately remove the intercept so that when gene expression is zero for any cell-specific genes, there are no cells of that type.

Equal weighting of the genes in the gene signature matrix by transforming them to the unit hypersphere means that both *X* and *Y* are transformed row-wise, where *X*_*i*,∗_ represents rows of *X* for *i =* 1, …, *m*. This would be equivalent to a weighted least squares (WLS) solution, where the weights are given by the inverse of the magnitude of each gene vector (data point) with the expression of the gene in different cell types, here expressed as matrix *Φ*.

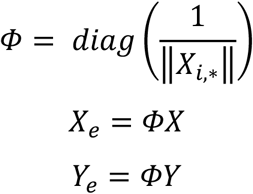

From here on, to simplify the equations, we write *X* and *Y* in place of the equal-weighted *X*_*e*_ and *Y*_*e*_. The equal-weighted *X* is further transformed column-wise as part of the vector projection, using *Ψ*:

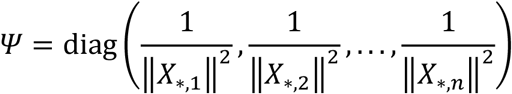

In NGD, the rows of the compensation matrix are modified to prevent non-negative *B* using the following formulation:

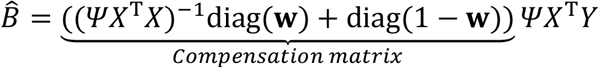

which can be rewritten

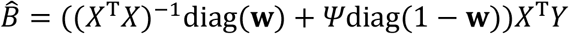

where **w** is a vector of weights bounded from 0 to 1 for each row of *B*. This allows each row of *B* to be tuned independently to be non-negative. When all **w** = 1, this resolves to OLS. When **w** = 0, this resolves to:

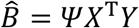

which makes sense as it is the original geometric projection without spillover. This also explains how NGD ensures non-negativity as *Ψ, X* and *Y* are all positive.

It can be interpreted that **w** varies the gram matrix smoothly between (*X*^T^*X*)^−1^ and *Ψ*. It is worth noting that *X*^T^*X* and *Ψ*^−1^ have the same diagonal elements. *Ψ* is the inverse of *X*^T^*X* if all the non-diagonal (i.e. covarying) elements of *X*^T^*X* are set to 0, i.e. *Ψ* is the idealised (*X*^T^*X*)^−1^ if *X* is orthonormal. This defines a sensible upper limit on *B*, based on projection with no spillover. Whereas L1/2 regularisation tends to shrink *B* towards 0, NGD accepts expansion of *B*, while defining an upper limit based on *Ψ*, i.e. that *B* is never greater than the basic projection with *Ψ* instead of (*X*^T^*X*)^−1^. Importantly, in NGD, *B* is not solved one sample at a time; all samples to be deconvoluted are solved simultaneously. Note also that genes which are not informative (for example constant across all cell clusters) have already been removed through the angle-based gene selection method and filtering process which removes low expressed genes.

This estimator equation for 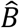 is the solution to the following optimisation:

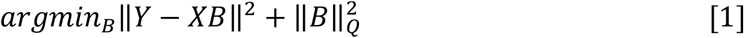

for *B* ≥ 0, where 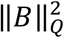 represents the weighted norm squared *B*^*T*^*QB* with

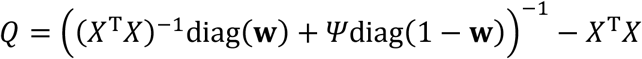

Thus, 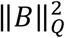 are Lagrange multipliers for the constraints of *B* having positive values.

Proof:

Expanding [1]

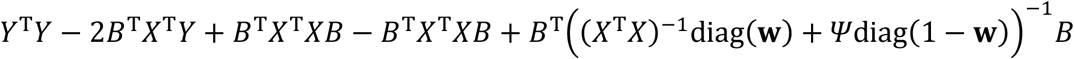

Differentiating this with respect to *B* and setting to zero to find minimum:

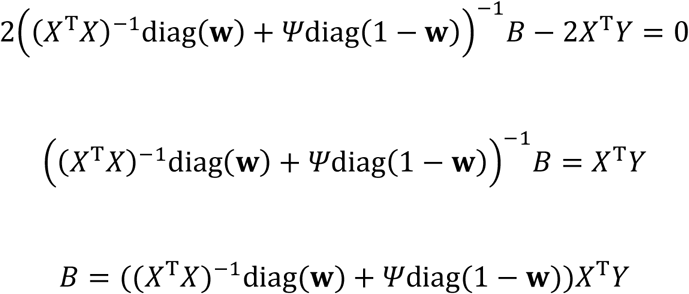

If all **w** = 1, the term 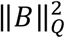 disappears and the optimisation reduces to OLS. If **w** = 0, the optimisation reduces to:

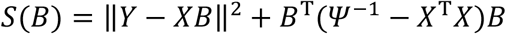

### Relationship with ridge regression

In Tikhonov regularisation, the regression model is formulated in matrix terms as:

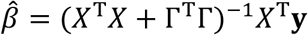

where for ridge regression the Tikhonov matrix Γ is chosen to be a multiple of the identity matrix *αI*.

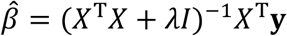

The ridge parameter *λ* is ≥ 0, so that *λ* serves as a constant shifting the diagonals of the moment matrix. In contrast, in NGD, the rows of the compensation matrix are varied between (*X*^T^*X*)^−1^ and *Ψ* smoothly according to **w**, in effect variably reducing the compensation for each row of *B* (i.e. each cell cluster), with the constraint that all values of *B* are >0. When *λ* = 0, ridge regression resolves to OLS; similarly, if **w** = **1**_n_, NGD resolves to WLS. Conversely, whereas increasing *λ* eventually shrinks all the coefficients to zero, reducing elements of **w** towards 0 allows rows of coefficients in *B* to inflate to an upper limit based on *Ψ* (**Supplementary Fig. 4**), which is useful in the non-negative setting. In comparison, while ridge regression increases the diagonal elements of *X*^T^*X* prior to inversion, in NGD, mixing with *Ψ* shrinks the non-diagonal elements of (*X*^T^*X*)^−1^ towards 0. Thus, NGD can be viewed as related to generalised Tikhonov regularisation. However, a major difference between the two approaches is that we choose to determine **w** heuristically, since we believe the main problem in real-world datasets is almost always due to excessive correction of spillover, whereas in ridge regression, the choice of *λ* is typically data-driven by cross-validation. By solving *B* for multiple bulk samples simultaneously and allowing rows of *B* with negative values to inflate, NGD preserves the proportional relationship between cell-specific genes and the bulk subclass cell counts *across samples*. This is a highly desirable feature, which is a major advantage over methods, which deconvolve each bulk sample individually, thus destroying the linear relationship of the cell subclass signature to the subclass counts between bulk samples. In contrast, the excessive application of the ill-conditioned inverse Gram matrix in other methods, will often lead to large and erroneous negative values. Even with a non-negative constraint, the most likely outcome is a poor overall solution which often erroneously underestimates certain cell populations especially those with high apparent compensation values.

Ridge regression (and Tikhonov regularisation more generally) have advantages over ordinary least squares with ill-posed problems, giving more reliable estimates of parameters when fitting models with large numbers of closely correlated independent variables, such as occurs with deconvolution of closely related cell types. In this situation, which is common in deconvolution, the spillover matrix is typically ill-conditioned and the inverse (the compensation matrix) is prone to error. This analogy may explain how the use of variable compensation in NGD is more effective and less likely to break down even with closely correlated cell types in comparison to W-CLS based methods, as well as being more tolerant of systematic differences between single-cell and bulk samples introduced by different sequencing methodologies.

### Standard errors of deconvolution

Viewing NGD as related to OLS/WLS means that it is possible to estimate standard errors on the deconvoluted cell counts. Based on OLS, the standard errors of 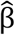 can be calculated as:

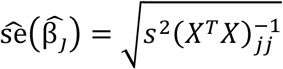

where *s*^2^ is the estimate of the sample variance. A correction to the exact OLS form, involves using *W* the modified version of (*X*^*T*^*X*)^−1^ derived from the compensation matrix which has been regularised to prevent non-negative 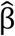:

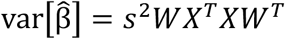

Bulk RNA-Seq is count data, which breaks the OLS assumption of homoscedasticity. Hence the third method available is White’s heteroscedasticity-consistent SE (HC0)^28^:

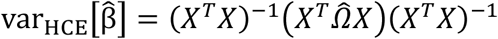

where

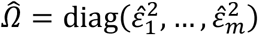

except that *W* replaces (*X*^*T*^*X*)^−1^. White’s method estimates the regression error variances as 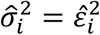. Refinements to White’s method have been proposed with newer heteroscedasticity-consistent methods referred to as HC1, HC2 and HC3^29^.

However, all of the above methods for estimating the variance of 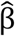 coefficients view each bulk sample separately. We note that the variance of the regression errors of β can be estimated directly from the variance of the cross-sample residuals for each gene. Rather than view the deconvolution as a collection of individual linear models with one model per bulk sample, the deconvolution is viewed as a general multivariate regression model *Y* = *XB* + *U*. In our case, the set of linear equations are considered as follows, without the intercept:

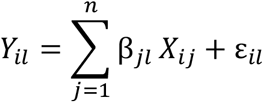

for observations (genes) indexed as *i* = 1, …, *m*, independent variables (cell types) indexed as *j* = 1, …, *n* and dependent variables (bulk samples) indexed as *l* = 1, …, *k*. Then, 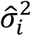, the variance of errors *e*_*i*_ for each observation in *Y* (i.e. each gene) is simply estimated as the variance of residuals for each gene across bulk samples:

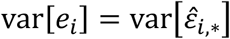

where 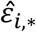 is row *i* of the weighted residuals matrix *ε* corresponding to the residuals for a single gene across different bulk samples. We calculate the standard error for each cell type as

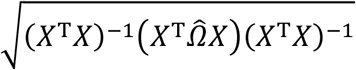

where

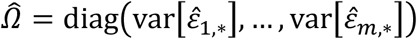

This appears to be an example of Stein’s phenomenon^42^: since many error estimators are being considered simultaneously across multiple bulk samples, inclusion of the variance of the residuals for each gene across samples improves the estimation of the variances of the errors compared to handling samples separately.

### Studentized residuals

Residuals are calculated as the difference between the actual measured bulk gene expression across samples and the fitted gene expression from the gene signature and fitted cell counts. Gene weights are applied first if they have been used.

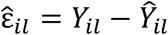

Standardized residuals are calculated based on the hat matrix

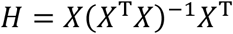

where *X* is the weighted gene signature matrix and (*X*^T^*X*)^−1^ is derived from the compensation matrix. Leverage *h*_*ii*_ = diag(*H*) is used to standardize the residuals

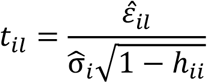

Studentized residuals are based on excluding the *i* th case (gene), calculated as follows

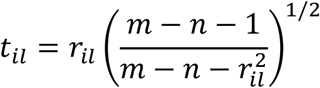

where *r*_*il*_ are standardized residuals and *n* is the number of predictors (cell subclasses; the intercept is not included). This corresponds to refitting the regression, but without recomputing the non-negative compensation matrix. Cook’s distance is calculated as

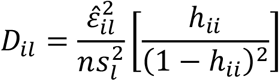

where 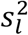 is the mean squared error for each bulk sample.

### Multipass removal of outlier genes based on extreme residual gene expression

Multipass deconvolution is activated by a user specified argument ‘npass’ for the number of passes. Each pass detects and removes genes which behave inconsistently due to noise originating from either the single-cell or bulk datasets as detected by extreme residual gene expression. Deconvolution during each pass is performed as usual, including compensation tuning unless the user has explicitly deactivated this. The var.e method calculates the variance of the residuals across bulk samples for each gene. Genes whose variance of residuals are outliers based on Z-score standardisation with |*Z*| > 4 are removed during successive passes. If no outliers are identified, the algorithm stops, so users can set a high number of passes and in our experience the algorithm will automatically stop after 3-4 passes unless the Z cut-off is set too low. Alternatively, users can opt to identify outlying genes based on Studentized residuals or Cook’s distance, which have default absolute outlier thresholds of >10 or >1 respectively.

### Simulation of pseudo-bulk RNA-Seq from scRNA-Seq

Included with cellGeometry are functions to simulate pseudo-bulk from single-cell RNA-Seq datasets. Simulated samples were generated with random cell counts for every cell type up to a maximum of the mean cell number per cell subclass x3-5. Thus cells per subclass ranged from ~0-8,700 for RA synovium; ~0-20,900 for Cell Typist blood; ~0-19,900 for Tabula Sapiens; ~0-29,000 for Human Brain Cell Atlas. Total cells per simulated bulk sample ranged from 49,300-101,300 cells for RA synovium; 49,300-101,300 cells for Cell Typist blood; 1.6-million cells for Tabula Sapiens; 1.4-1.9 million cells for Human Brain Cell Atlas. Simulations contained 30-50 random samples and each simulation was repeated 3-5 times. To simulate pseudo-bulk counts, for each sample we directly sample at random with replacement from the cells in the original scRNA-Seq dataset and calculate the sum of the total counts per gene. Rather than sample cells uniformly, which is problematic if the number of cells to sample exceeds the number of cells in a cluster (as is common), cell sampling is performed according to the Dirichlet distribution of order *K*, where *K* is the number of cells in each cluster. Dirichlet distribution sampling avoids uniformity of sampling when there is oversampling.

### Curation and deconvolution of simulation data

Simulated pseudo-bulk data was constructed by Dirichlet based sampling with known cell counts for each subclass from the respective sc/snRNA-Seq datasets (RA synovium^15^, Cell Typist blood^18^, Tabula Sapiens^2^, Human Brain Cell Atlas single-nucleus^3^). Pseudo-bulk RNA-Seq data were deconvoluted using cellGeometry using the unfiltered arithmetic mean for the gene signature; MuSiC^9^ (MuSiC1 version 0.2.0; MuSiC2^17^ version 1.0.0), DWLS^10^ (version 0.1.0) or LinDeconSeq^11^ (version 0.1) using default settings. Initially both MuSiC1 and MuSiC2 were tested. But as both versions produced identical results across all 4 scRNA-Seq datasets and showed identical times in benchmark speed tests, MuSiC2 was used throughout the rest of the study. We also tested two more recent deconvolution tools which have been applied to atlas-level datasets: Bisque which uses constrained least squares and is published as BisqueRNA (version 1.0.5) and InstaPrism (version 0.1.6). Comparisons were also made against deconvolution by non-negative least squares using nnls (version 1.4) and elastic net regression with non-negative constraint using glmnet^43^ (version 4.1-3) with the setting lower.limits=0 across a range of alpha from 0 (ridge) to 1 (LASSO). NMF based methods investigated used NMF^20^ (version 0.28) using the Brunet, Killback-Leibler (KL) or Lee algorithms, and RcppML^21^ (version 0.5.6). This was undertaken with 5 replicates, with the exception of the Human Brain Cell Atlas data where 3 replicates were used.

Simulated data of the Human Brain Cell Atlas was generated by generating the cell counts and pseudo-bulk RNA-Seq for neuron and non-neuronal separately. The cell counts were merged through column binding, whereas the pseudo-bulk RNA-Seq were merged through summing the gene expression (**Fig. 3a**). The gene signatures for neuron and non-neuronal cells were generated separately and merged using the mergeMarkers function of cellGeometry in which optimal gene signatures for the merged cell subclasses are recalculated (**Fig. 3b**). Since the two datasets were sequenced and reported by the same group^3^, the two datasets did not require transformation when merging the gene signatures as illustrated by the quantile-quantile plot where the curve is sufficiently close to the line of identity (**Fig. 3c**).

### Downsampling of single-cell datasets

As DWLS, LinDeconSeq and Bisque are not programmed to handle matrices with >2^31^ elements, the larger scRNA-Seq datasets (Tabula, Brain) were downsampled to smaller matrices within the 2^31^ size limit, which were used as the scRNA-Seq reference inputs for both methods. Downsampling was conducted to maintain sufficient representation of all cell subclasses, so that for subclasses with 10-100 cells all cells were preserved. Subclasses with <10 cells were discarded. Downsampling was also tested on MuSiC.

### Simulation of experimental noise and absent cell types

Different types of noise with varying standard deviation were applied to the simulated pseudo-bulk data derived from the AMP RA synovium scRNA-Seq dataset. ‘Gaussian noise’ is the simple addition of Gaussian noise to bulk counts, so that low expressing genes are affected, but highly expressed genes are hardly affected. For ‘Log noise’, counts are converted to log_2_+1 scale before the addition of Gaussian noise and then transformed back. Log noise affects all genes irrespective of expression level. For ‘Sqrt noise’, counts are square rooted before the addition of Gaussian noise and then transformed back to count scale. Sqrt noise has a stronger effect on moderately expressed genes compared to standard Gaussian noise and the distribution of expression from the original simulated data compared to the simulated data with Sqrt noise is similar to that of the technical noise in scRNA-Seq experiments conducted by Brennecke *et al*^27^. For ‘Shift noise’, a proportion (50%) of genes are selected at random and each gene is multiplied by a random amount varying according to 2^random number drawn from a normal distribution. Note that whereas the other methods apply randomly different amounts of noise to each sample, in Shift noise, the per gene effect affects all samples in the same way. This simulates differences in chemistry between scRNA-Seq and bulk RNA-Seq, causing randomly selected whole genes to be detected at higher or lower levels.

Absence of cell types was simulated in pseudo-bulk data derived from scRNA-Seq datasets by selecting a varying proportion (0%, 5%, 10%, 20% and 50%) of cell subclasses at random, whose cell counts were set to zero prior to sampling from the scRNA-Seq reference.

### Evaluation metrics

With true cell type proportions denoted by *p* and estimated proportions denoted by 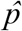, deconvolution methods were evaluated by the following metrics:

i. Coefficient of determination: 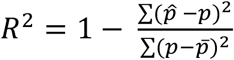
ii. Root Mean Square Error: 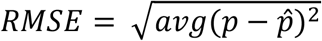

Lin’s Concordance Correlation Coefficient (CCC) was calculated using the DescTools package (version 0.99.60).

### Benchmarking

When measuring the running time of the deconvolution, deconvolution using the RA synovium dataset was undertaken on Intel Xeon W-2145 processor, 256 GB RAM. Cell Typist blood, Tabula Sapiens and the Human Brain Cell Atlas deconvolution were undertaken on an AMD Ryzen Threadripper PRO 3975WX workstation, 256 GB RAM, both running Ubuntu Linux. MuSiC, DWLS and LinDeconSeq are single-core methods. cellGeometry was run using 1 core for AMP, and both 1 and 8 cores were compared for Tabula Sapiens and Human Brain Cell Atlas. DWLS, LinDeconSeq and Bisque are unable to handle large matrices with >2^31^ elements. In order to compare these methods scRNA-Seq matrices were downsampled to smaller matrices.

### Deconvolution of blood samples

Whole blood bulk RNA-Seq samples used for benchmarking deconvolution were from the Pathobiology of Early Arthritis Cohort (PEAC) observational study^30^. The study received ethical approval from the UK Health Research Authority (REC 05/Q0703/198, National Research Ethics Service Committee London – Dulwich). Following written informed consent from patients, RNA from whole blood samples was stored, processed and bulk RNA-Sequenced as described previously^30^.

cellGeometry deconvoluted the whole blood samples using Cell Typist single cell blood dataset^18^ merged with basophils and neutrophils single cell gene signatures from the Tabula Sapiens dataset restricted to blood^2^. Gene signature cellMarker objects for each dataset were curated separately using default settings and merged through quantile-quantile mapping of the signature matrix gene expression distributions for each single-cell dataset. Deconvolution of the whole blood samples using Cell Typist blood dataset^18^ or Tabula Sapiens blood dataset^2^ alone by MuSiC, DWLS and LinDeconSeq.

Subclass proportions were visualised as stacked barplots and the mean total of subclass proportions according to cell groups were visualised as pie charts. Peripheral blood lymphocytes and neutrophils were quantified by Coulter counter and correlated with the lymphocyte and neutrophil proportions from deconvolution by Pearson’s correlation.

### Deconvolution of rheumatoid arthritis synovium biopsies

Synovium biopsies used for deconvolution originated from the PEAC study^30^, the Stratification of biological therapies for RA by Pathobiology (STRAP) randomised controlled trial^37^ and the R4RA trial^33^. The STRAP and R4RA trials were approved by the UK ethics committee (MREC 14/WA/1209 and 12/WA/0307 respectively). All patients provided written informed consent. For STRAP and R4RA, synovial biopsies were obtained through ultrasound-guided or arthroscopic procedure and were processed as described previously^33,37^. For PEAC, synovial biopsies were obtained through ultrasound-guided procedure. For consistency with STRAP and R4RA, PEAC RNA-Seq data were re-mapped to Gencode release v29 GRCh38 and quantified using Salmon. Deconvolution was performed using AMP RA synovium scRNA-Seq^15^ as reference with cellGeometry, MuSiC, DWLS or LinDeconSeq with default settings.

Subclass proportions were visualised as stacked barplots per patient and with respect to histological pathotype. SC-B4 and B cell, T cell and macrophage levels were correlated using Spearman correlation test against semi-quantitative scoring of the respective immunohistology cell surface marker, namely CD20, CD3 and CD68 respectively. Semi-quantitative histology scoring was performed using a histology atlas for each marker^30,33,44^.

### Determining thresholds for pathotype redefinition

Decision trees were generated by rpart (version 4.1.24) to predict the histology pathotypes, lymphoid vs non-lymphoid and myeloid vs non-myeloid, based on cellGeometry results on B cell and macrophage cell frequencies respectively. This analysis was undertaken by sampling 90% of the data (divided into 10-folds, one fold removed each time) with 15 repeats of the 9/10-folds (total 150 samples).

## CODE AVAILABILITY

cellGeometry is publicly available as an R package which can be installed from CRAN (https://cran.r-project.org/package=cellGeometry). Source code is available on Github at https://github.com/myles-lewis/cellGeometry. Scripts used for benchmarking and figure generation are available from https://github.com/EMR-bioinformatics/cellGeometry_manuscript/

## DATA AVAILABILITY

Single-cell datasets are available from https://www.immport.org/shared/study/SDY998 (AMP phase 1 RA synovium), https://cellxgene.cziscience.com/collections/62ef75e4-cbea-454e-a0ce-998ec40223d3 (Cell Typist), https://cellxgene.cziscience.com/collections/e5f58829-1a66-40b5-a624-9046778e74f5 (Tabula Sapiens) and https://cellxgene.cziscience.com/collections/283d65eb-dd53-496d-adb7-7570c7caa443 (Human Brain Cell Atlas). PEAC study blood and synovium bulk RNA-Seq, R4RA and STRAP clinical trial bulk synovium RNA-Seq datasets are available from ArrayExpress under accession codes E-MTAB-6141, E-MTAB-11611 and E-MTAB-13733.

## Acknowledgements

This work acknowledges the support of the National Institute for Health Research Barts Biomedical Research Centre (NIHR 203330). The PEAC study was supported by funding from the UK Medical Research Council (MRC) (grant number G0800648). The STRAP study was supported by MRC and Arthritis Research UK (ARUK) by their joint funding of Maximizing Therapeutic Utility in Rheumatoid Arthritis (MATURA) (grant numbers MR/K015346/1 and 20670 respectively). The R4RA study was supported by the UK National Institute of Health Research (grant reference: 11/100/76). Core work associated with this project was supported by Versus Arthritis (Experimental Arthritis Treatment Centre, grant number 20022). Other support includes Barts and The London School of Medicine and Dentistry charity (grant number 523/819), NIHR EME (grant 131575) and MRC TRACT-RA (MR/ V012509/1). A.S. is supported by a Career Development Fellowship from Arthritis UK (MT/23270). The views expressed are those of the authors and do not represent those of the NHS, NIHR or our funding bodies.

## Author contributions

M.J.L. created the cellGeometry R package. R.L. tested and benchmarked the package and generated results. P.M.P. & A.E.A.S. performed additional testing of the software on other datasets and helped with debugging. C.C. processed the RNA-seq data and helped with debugging. L.F.J. oversaw processing of RNA samples. C.P. provided bulk and scRNA-Seq samples through overall supervision of clinical studies (PEAC) and randomised clinical trials (R4RA, STRAP). M.J.L. and R.L. wrote the manuscript with input from S.R.. M.J.L. & A.S. reviewed the mathematical formulation. All authors participated in the revision of the final paper.

