## Supplementary Material for "cellGeometry: ultra-fast single-cell deconvolution of bulk RNA-Seq using a geometric solution"

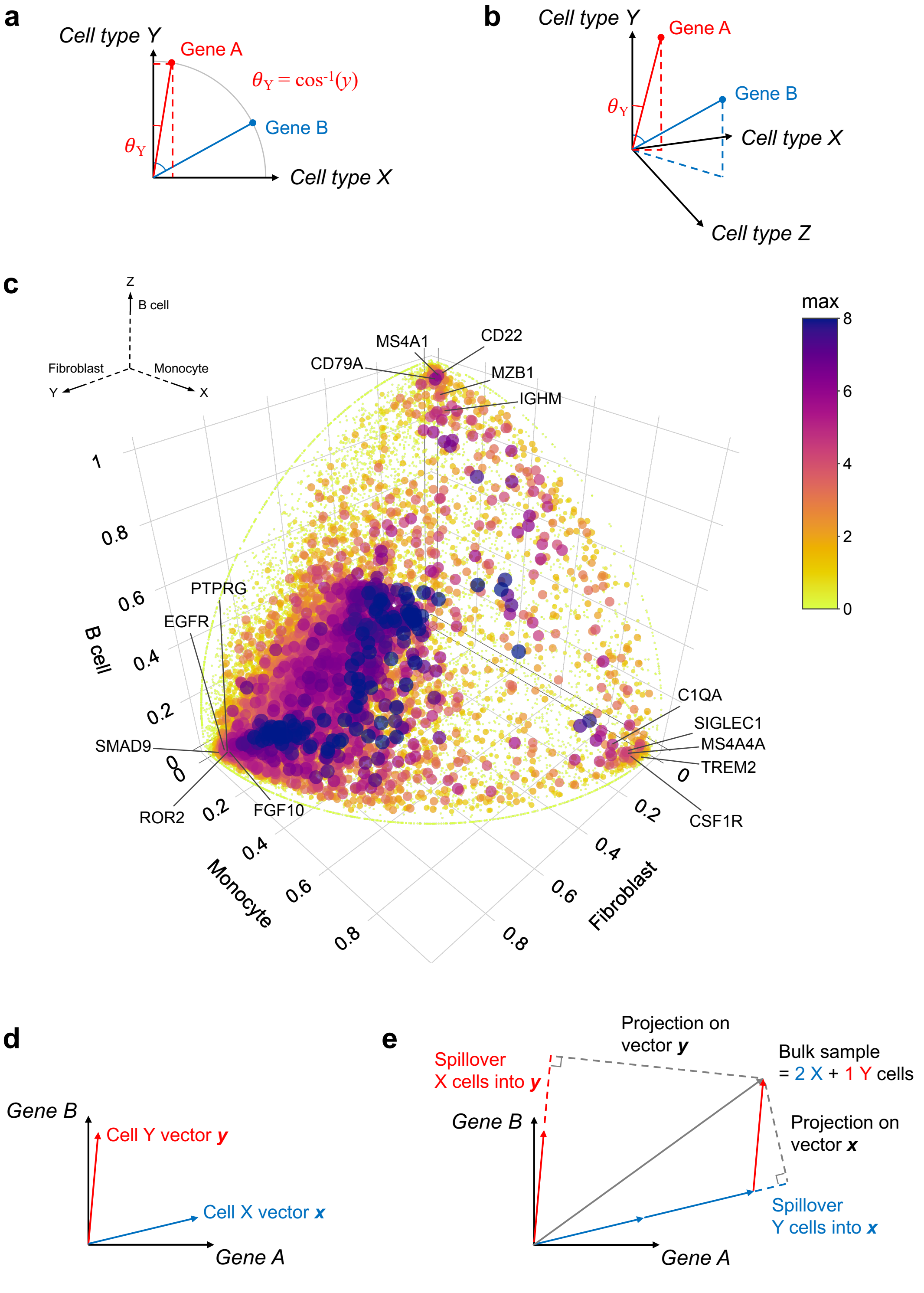


**Supplementary Fig. 1. Different geometries are used for identifying cell-specific genes and performing deconvolution**
**a & b,** Geometry of the angle method for identifying cell specific genes showing simplified examples with only 2 cell types **(a)** or 3 cell types **(b)**. Each gene is represented as a point in space with coordinates as the mean gene expression in each cell type with *cell types* as dimensions in high dimensional space. Each gene vector is scaled to the unit hypersphere. Arc cosine determines the angle for each gene with the axis for that cell cluster. The most cell-specific genes have the lowest angle relative to that cell cluster axis. **c,** 3D plot of fibroblast (*y* axis), monocyte (*x* axis) and B cell (*z* axis) mean gene expression of all genes from a rheumatoid arthritis synovium scRNA-Seq dataset[1] to illustrate how cellGeometry selects the most cell-specific gene markers. In this illustrative example only 3 cell types are used. The most cell-specific genes are those closest to the dimensional axis for that cell type. Genes known to be highly specific for each cell type are labelled. Point colour shows maximum mean $\log_2$ gene expression per cell type. **d & e,** Geometric method of deconvolution. **d,** The gene signature for each cell cluster is represented as a vector in high dimensional *gene* space. **e,** In this simplified example in only 2 dimensions with only 2 signature genes, the bulk sample is represented as a vector. The amount of X cells in the bulk sample is calculated by projecting the bulk sample vector on the gene signature vector for the X cell type. The illustration shows how spillover occurs when genes are not 100% specific for a cell type. Spillover is corrected using a compensation matrix (see Methods).

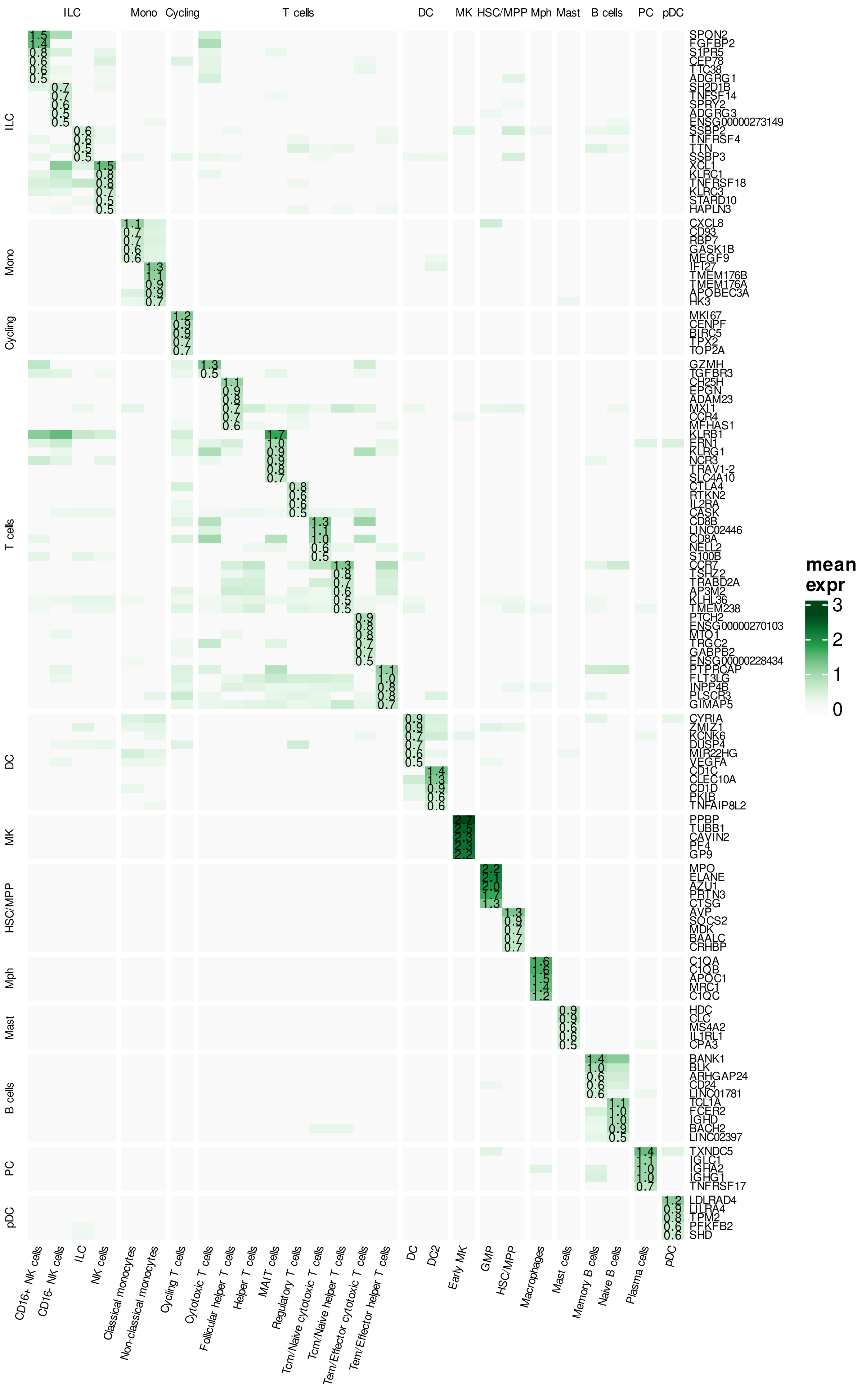

**Supplementary Fig. 2. see overleaf**

**Supplementary Fig.2. Gene signature produced by cellGeometry on the Cell Typist blood dataset**
Heatmap of the gene signature expression matrix generated by cellGeometry for the Cell Typist blood dataset[2] showing the top 5 genes for each cell subclass ranked by specificity angle and expression.

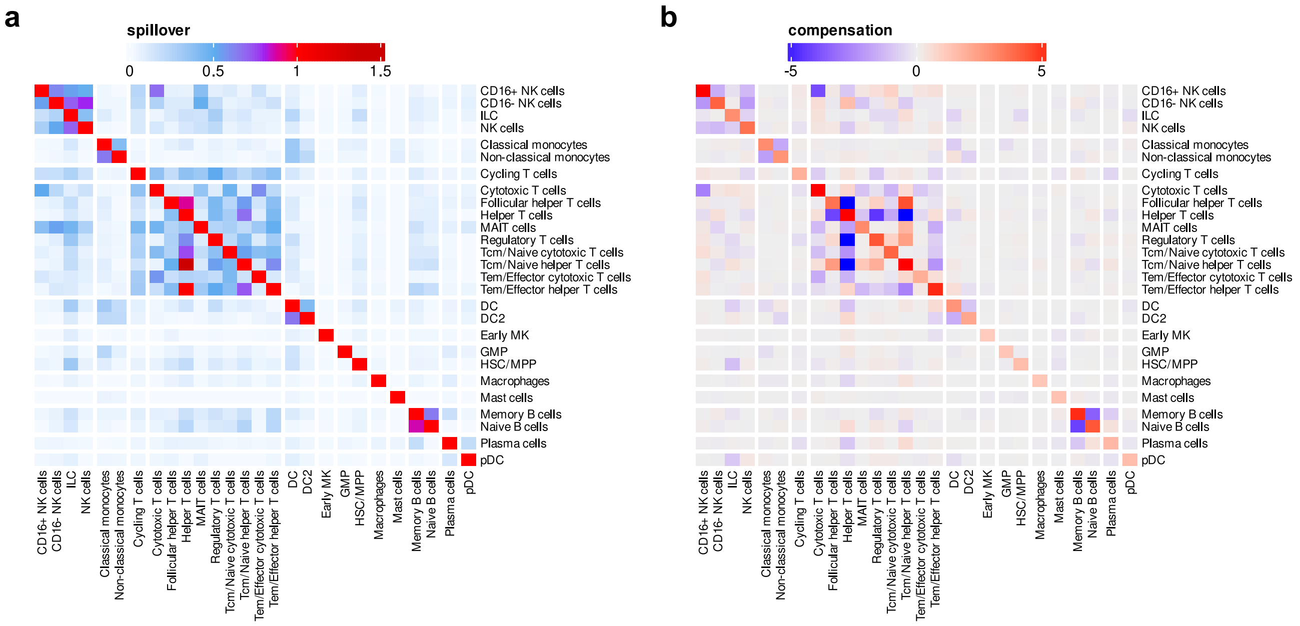


**Supplementary Fig. 3. cellGeometry spillover and compensation heatmaps for the Cell Typist dataset**
**a,** Heatmap of the spillover matrix produced by cellGeometry for the Cell Typist blood scRNA-Seq dataset[2] showing the amount of spillover of gene signatures for each cell subclass vector into other subclasses.
**b,** Heatmap of the compensation matrix for the Cell Typist blood dataset cell subclasses used for deconvolution by cellGeometry.

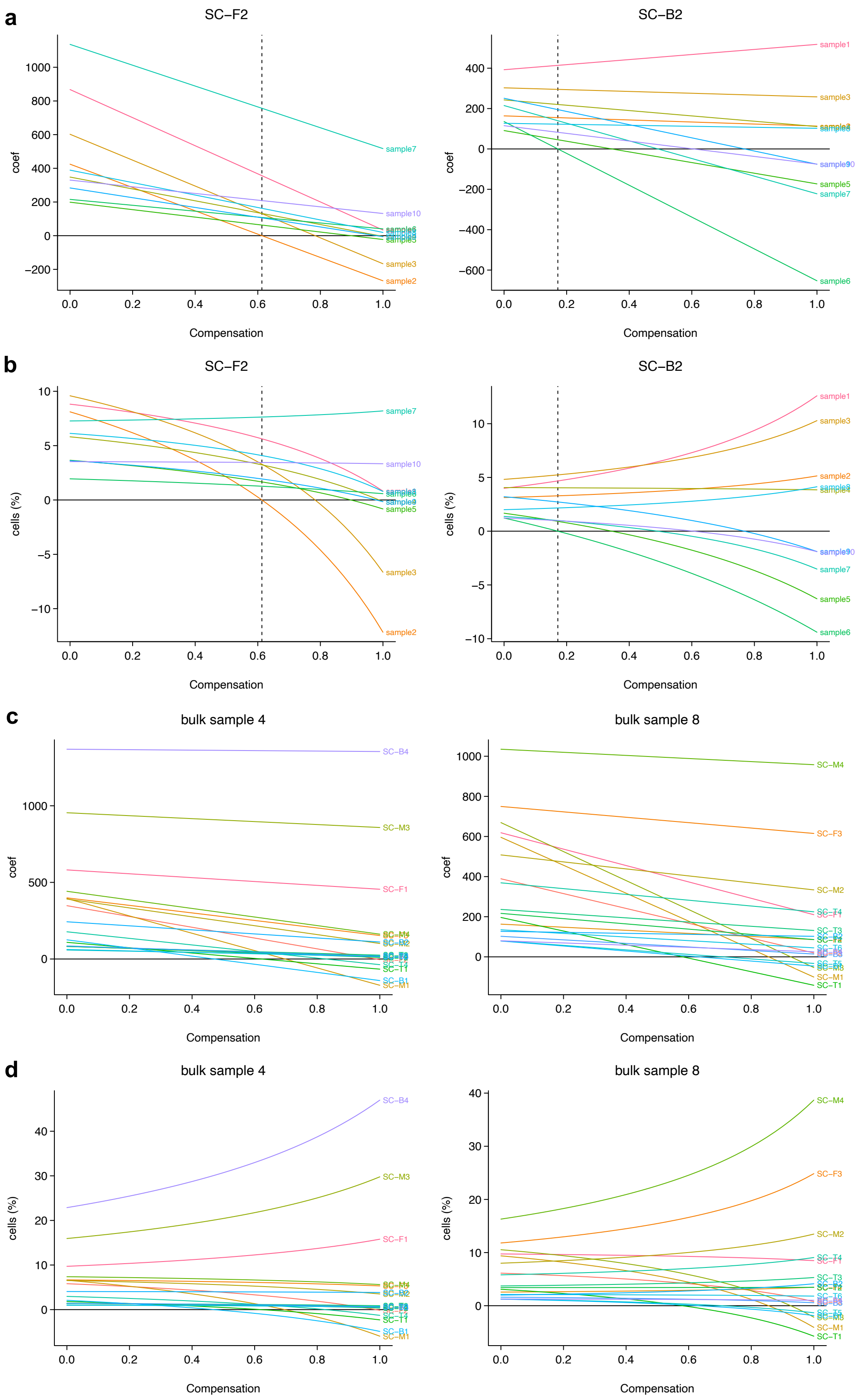


**Supplementary Fig. 4. Coefficient path plots showing the effect of varying compensation**

Example coefficient path plots generated by varying the compensation vector **w** from 0-1 for each cell subclass. Real bulk synovial RNA-Seq samples were deconvoluted using AMP scRNA-Seq as reference. **a,b)** shows coefficient paths for 10 samples for 2 individual cell subclasses, SC-F2 fibroblasts and SC-B2 B cells. Vertical dashed line shows the final optimised, chosen compensation value for each subclass. **c,d)** shows coefficient paths for two individual bulk samples, showing the paths for all 18 cell subclasses. All bulk samples are deconvoluted together to preserve the relationship for each individual cell subclass across bulk samples. **a** & **c** show cell counts, **b** & **d** show cell percentages out of total cells.

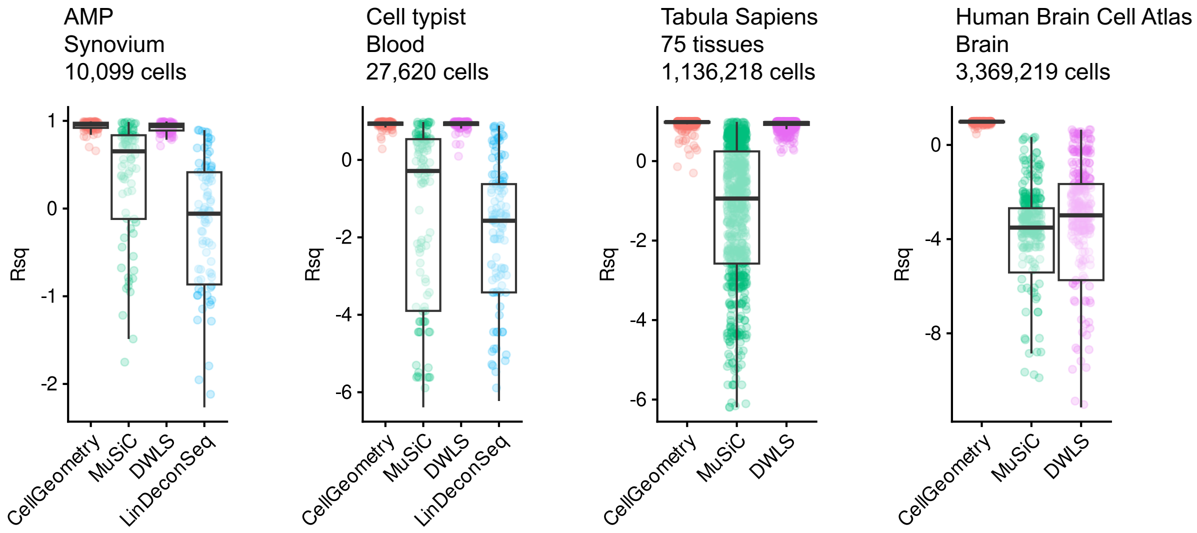


**Supplementary Fig. 5. Benchmarking deconvolution accuracy with simulated pseudo-bulk from four reference single-cell datasets**
Boxplots of the coefficient of determination ($R^2$) for cell subclass percentages from deconvolution of simulated pseudo-bulk datasets generated from Accelerated Medicines Partnership (AMP) rheumatoid arthritis synovium[1], Cell Typist blood[2], Tabula Sapiens[3] and Human Brain Cell Atlas[4] scRNA-Seq datasets. Deconvolution methods assessed were cellGeometry, MuSiC, DWLS and LinDeconSeq. Five replicates of simulated pseudo-bulk data (N = 25 samples) were generated, apart from Human Brain Cell Atlas where 3 replicates (N = 30 samples) were generated. Boxplots illustrate median, upper and lower quartiles with whiskers denoting maximal and minimal data within 1.5× interquartile range. *y* axis is cropped at the minimum whisker values to reduce plot distortion by outliers.

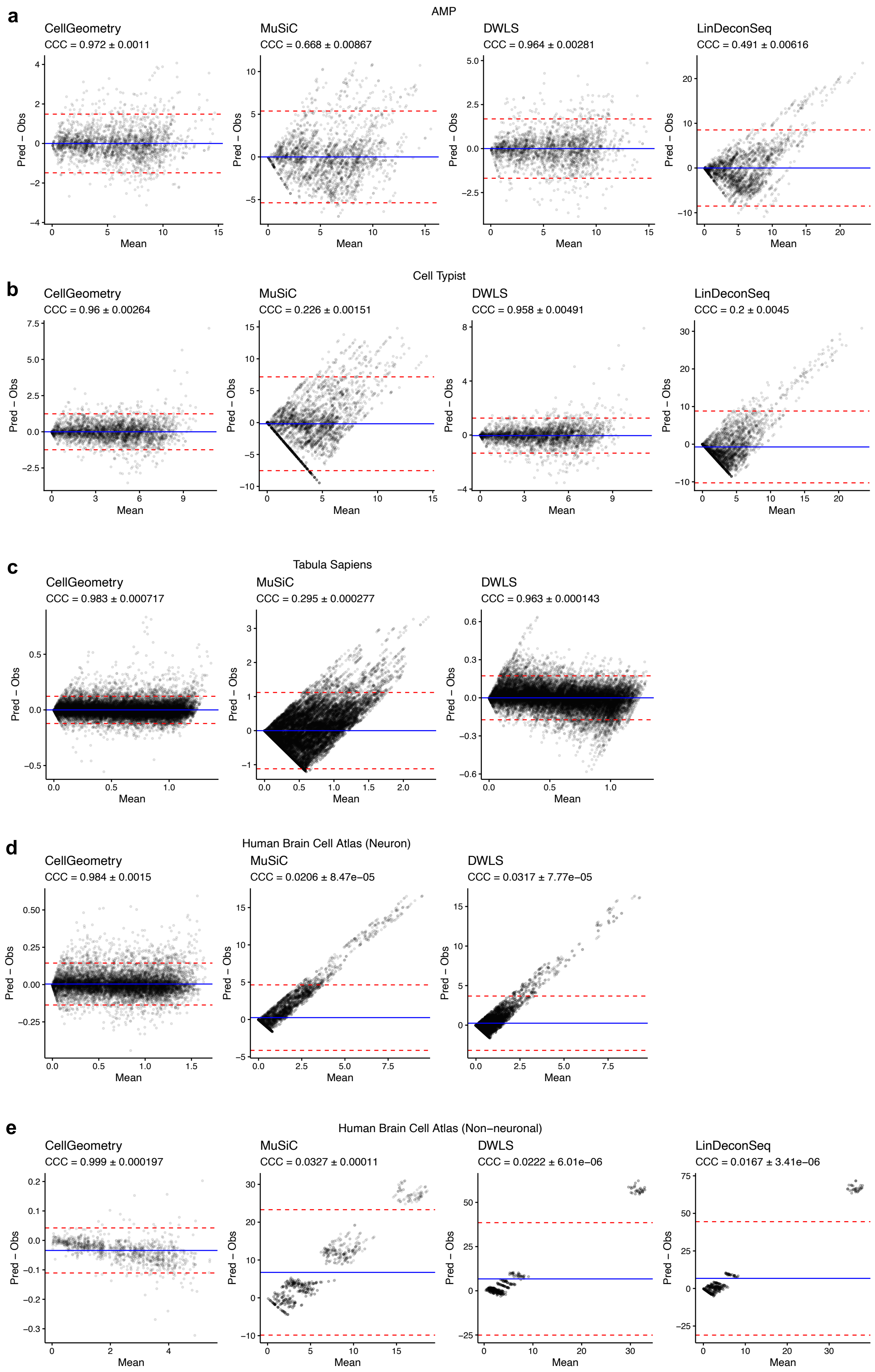


**Supplementary Fig. 6. Comparing the agreement between predicted and ground truth cell percentages with simulated pseudo-bulk from four reference single-cell datasets**

Bland-Altman plots visualising the differences between predicted and observed cell percentages against their mean from deconvolution of simulated pseudo-bulk datasets generated from **(a)** Accelerated Medicines Partnership (AMP) rheumatoid arthritis synovium[1], **(b)** Cell Typist blood[2], **(c)** Tabula Sapiens[3] and Human Brain Cell Atlas[4] **(d)** neuron and **(e)** non-neuronal scRNA-Seq datasets. Deconvolution methods assessed were cellGeometry, MuSiC, DWLS and LinDeconSeq. Five replicates of simulated pseudo-bulk data (N = 25 samples) were generated, apart from Human Brain Cell Atlas where 3 replicates (N = 30 samples) were generated. Blue line = mean difference (bias). Red lines = mean difference ± 1.96 standard deviation (limits of agreement). Statistical analysis by Lin's concordance correlation coefficient (CCC) ± 95% confidence intervals.

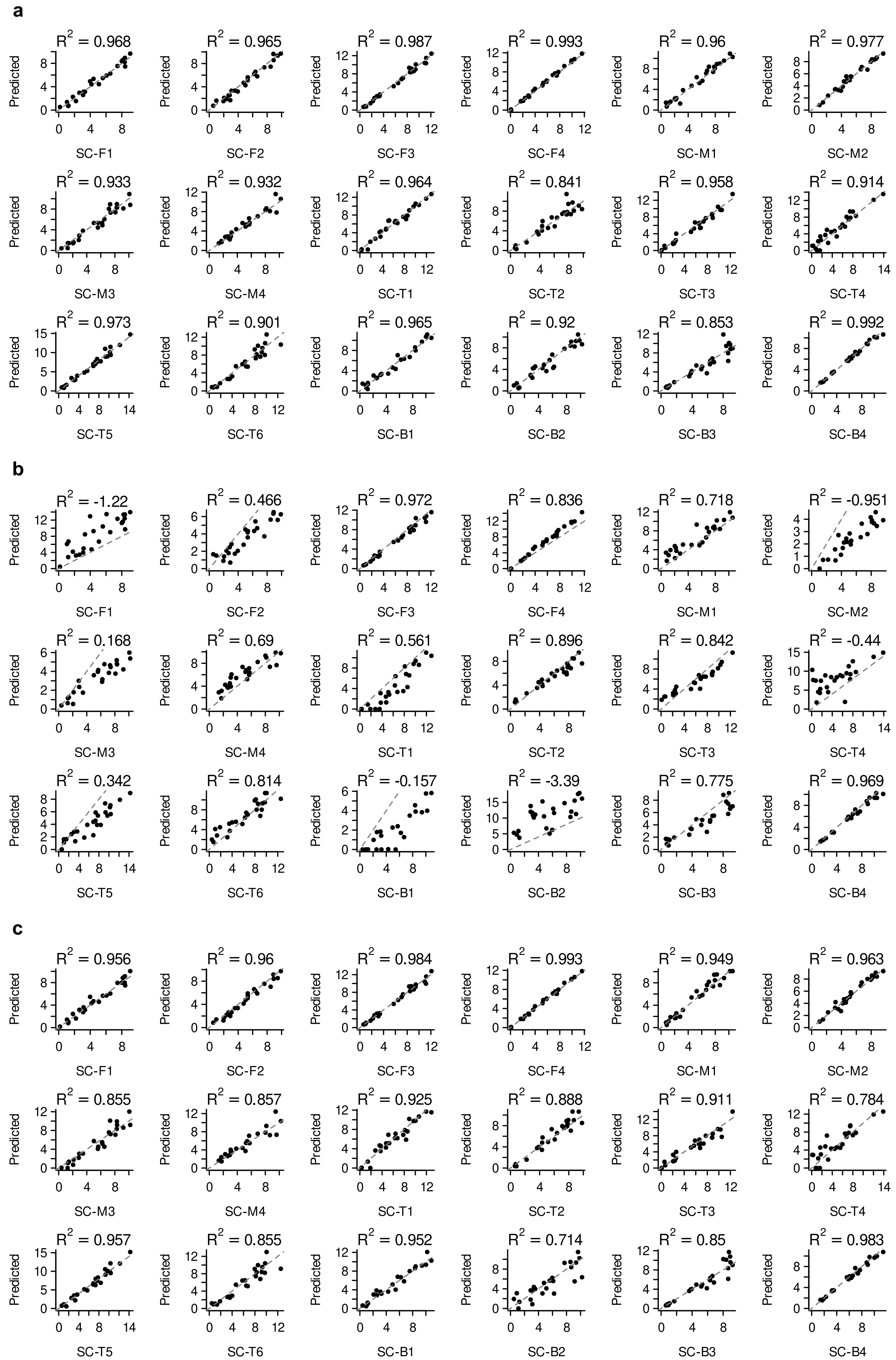


**Supplementary Fig. 7. continued overleaf**

**d**

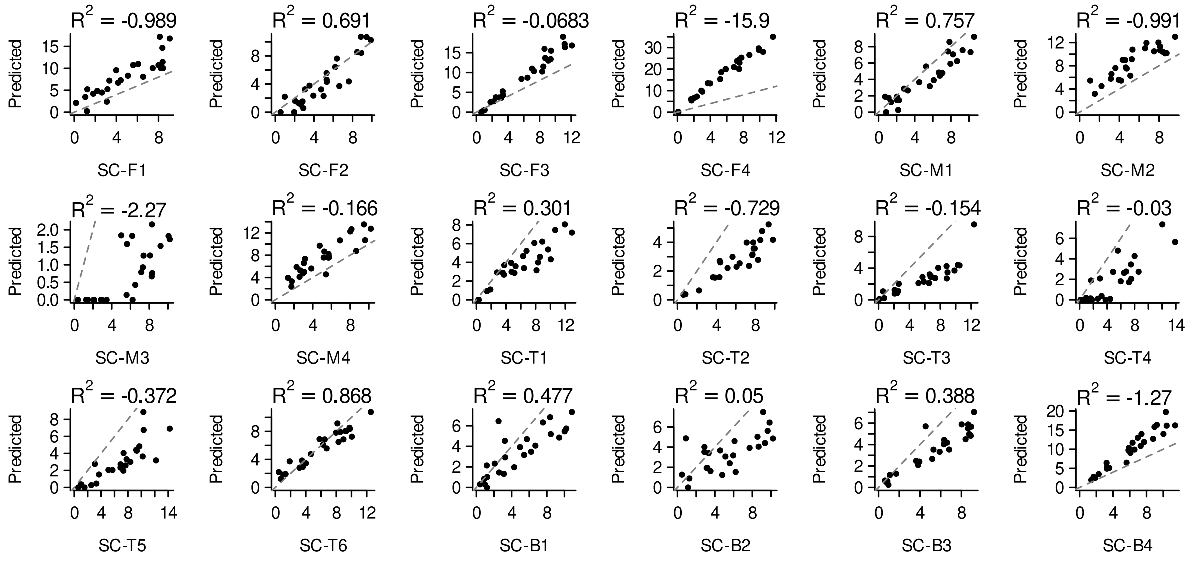


**Supplementary Fig. 7. Evaluation of different deconvolution methods with simulated synovium pseudo-bulk data showing predicted versus true cell percentages**

Deconvolution of simulated pseudo-bulk data (N = 25) from a rheumatoid arthritis synovium single-cell dataset[1] was undertaken using **(a)** cellGeometry, **(b)** MuSiC, **(c)** DWLS and **(d)** LinDeconSeq. Scatter plots show predicted versus true cell percentages for each cell subclass. The coefficient of determination ($R^2$) was used to evaluate the accuracy of predicted cell type percentages. The identity line is shown in dashed grey.

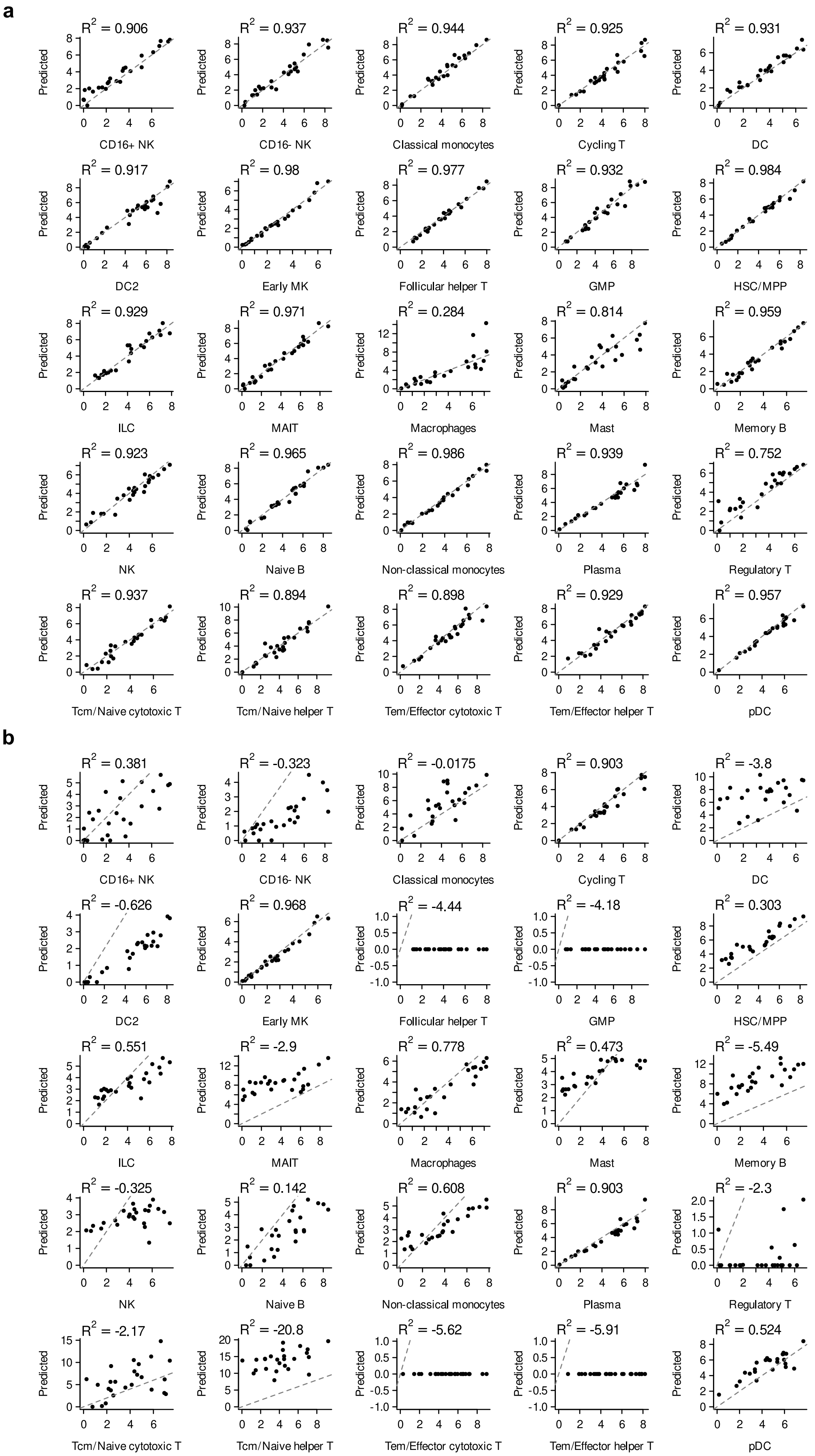


**Supplementary Fig. 8. continued overleaf**

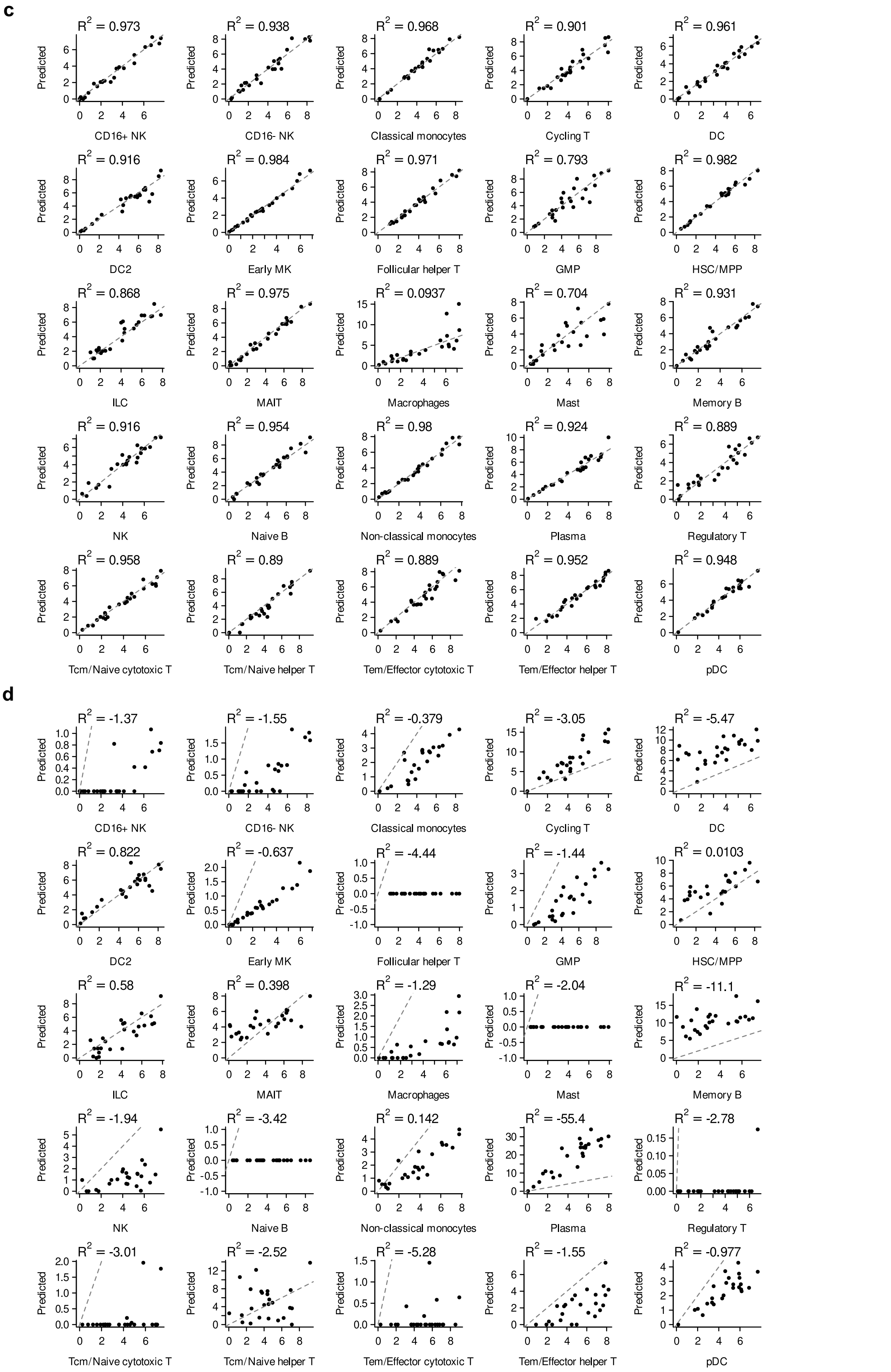


**Supplementary Fig. 8. True and estimated cell type percentages of simulated blood pseudo-bulk data using Cell Typist dataset evaluating different deconvolution methods**

Deconvolution of simulated pseudo-bulk data (N = 25) using the Cell Typist blood scRNA-Seq dataset[2] was undertaken using **(a)** cellGeometry, **(b)** MuSiC, **(c)** DWLS and **(d)** LinDeconSeq. Scatter plots show predicted versus true cell percentages for each cell subclass. The coefficient of determination ($R^2$) was used to evaluate the accuracy of predicted cell type percentages. The identity line is shown in dashed grey.

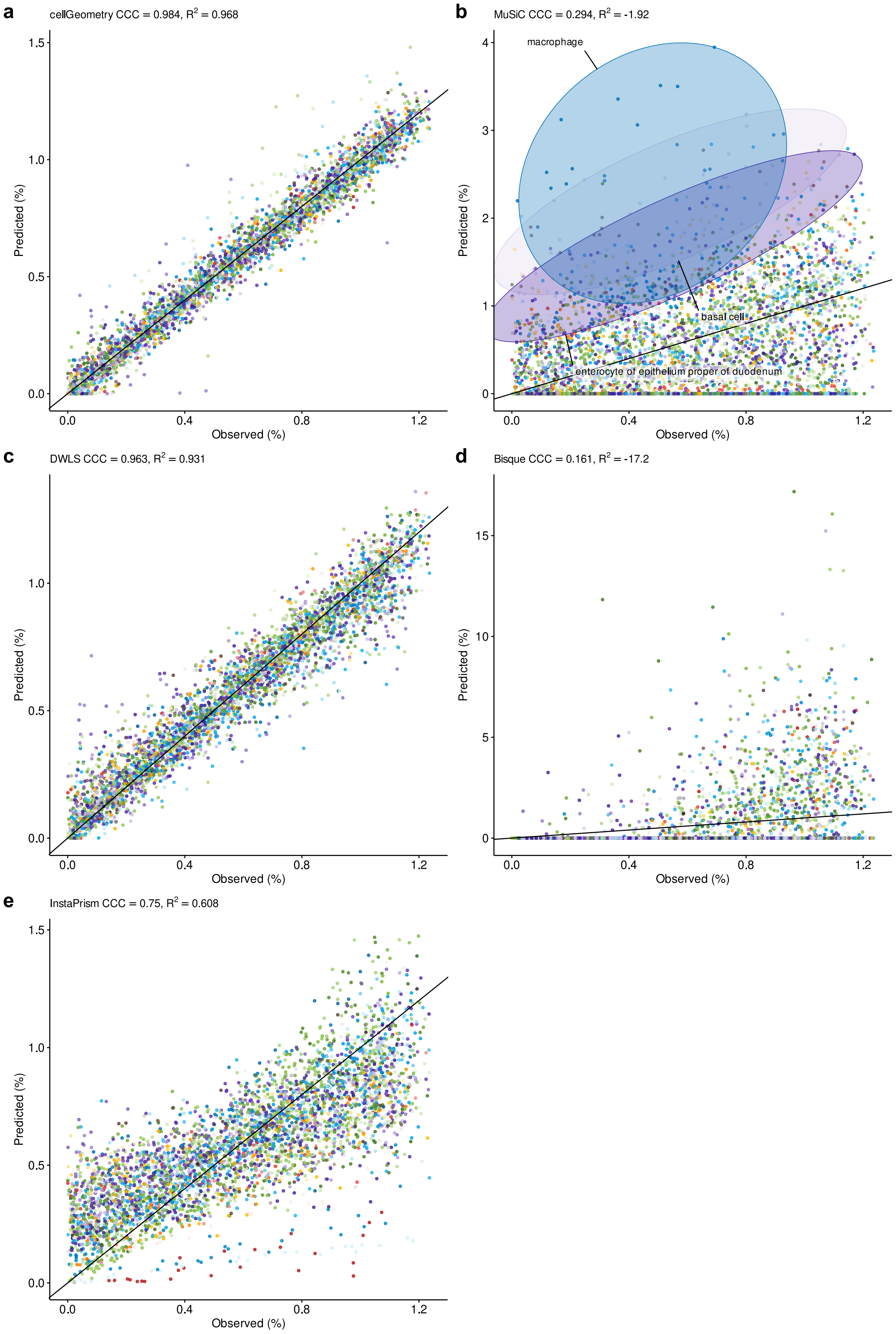


**Supplementary Fig. 9. Evaluating the accuracy of cellGeometry, MuSiC, DWLS, Bisque and InstaPrism in estimating cell type percentages of simulated multi-tissue bulk data using Tabula Sapiens dataset**
Deconvolution of simulated pseudo-bulk data (N = 25) using the Tabula Sapiens scRNA-Seq dataset[3] was undertaken using **(a)** cellGeometry, **(b)** MuSiC, **(c)** DWLS, **(d)** Bisque or **(e)** InstaPrism. Scatter plots show predicted versus true cell percentages. The coefficient of determination ($R^2$) was calculated to evaluate the accuracy of predicted cell type percentages. Point colours represent different cell subclasses. The identity line is shown.

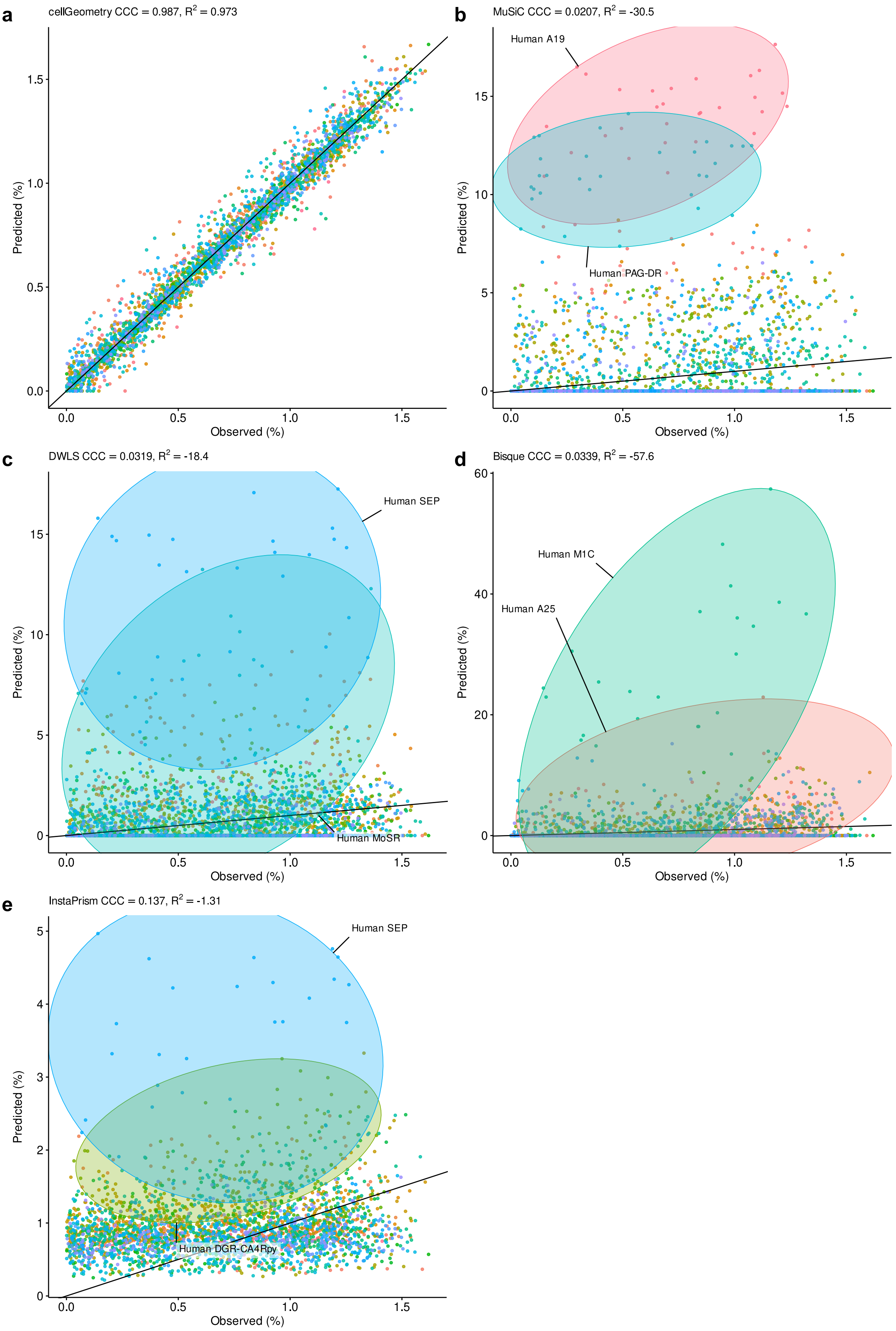


**Supplementary Fig. 10. Evaluating the accuracy of cellGeometry, MuSiC, DWLS, Bisque and InstaPrism in estimating neuron subset percentages in simulated brain pseudo-bulk**

Deconvolution of simulated pseudo-bulk data (N = 30) from the Human Brain Cell Atlas[4] was undertaken using **(a)** cellGeometry, **(b)** MuSiC, **(c)** DWLS, **(d)** Bisque or **(e)** InstaPrism. Scatter plots show predicted versus true cell percentages of neuron cell types. The coefficient of determination ($R^2$) was calculated to evaluate the accuracy of predicted cell type percentages. Point colours represent different cell subclasses. The identity line is shown.

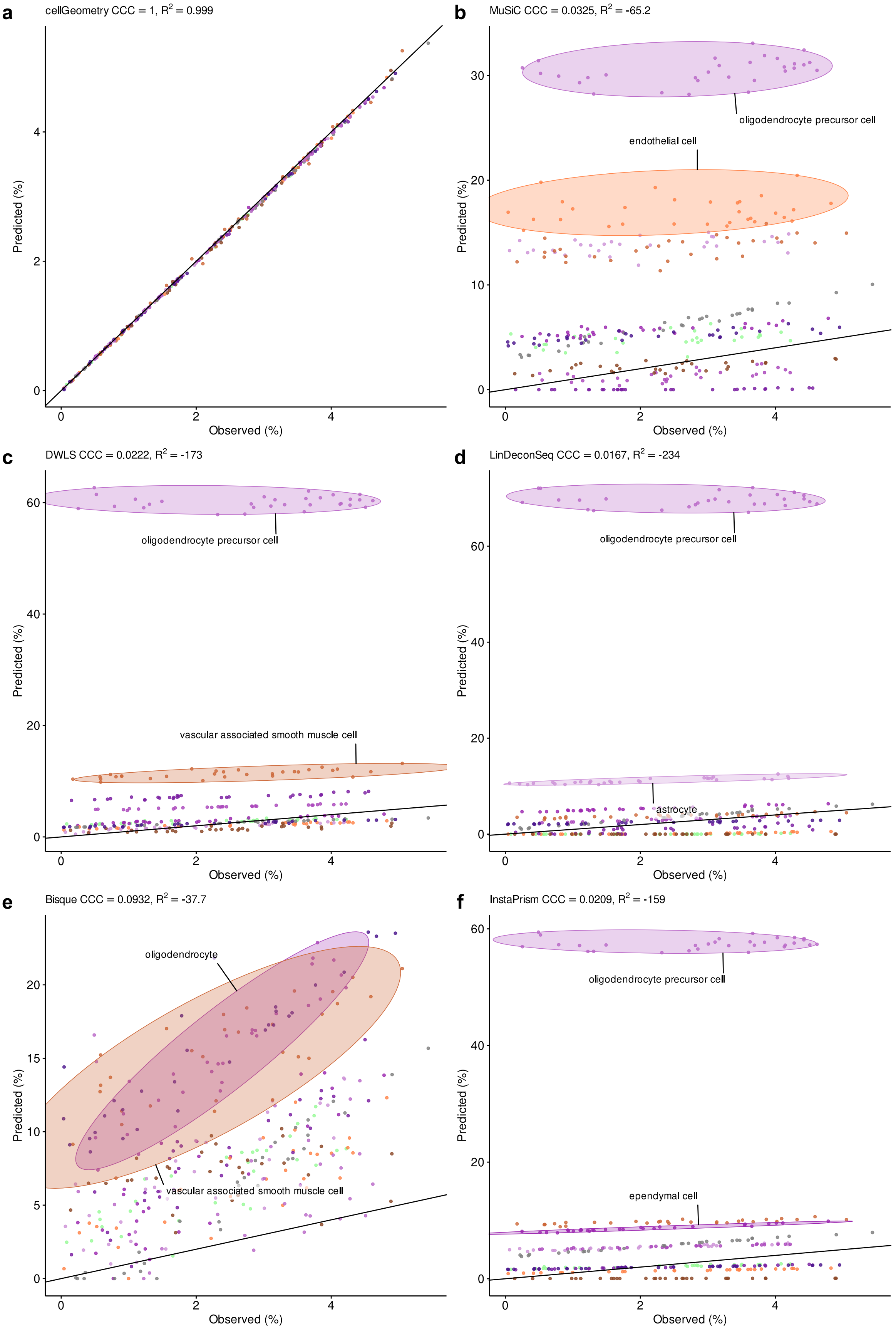


**Supplementary Fig. 11. Evaluating accuracy of estimating non-neuronal cell percentages in simulated brain pseudo-bulk samples**

Deconvolution of simulated pseudo-bulk data (N = 30) from the Human Brain Cell Atlas dataset[4] was undertaken using **(a)** cellGeometry, **(b)** MuSiC, **(c)** DWLS, **(d)** LinDeconSeq, **(e)** Bisque or **(f)** InstaPrism. Scatter plots show predicted versus true cell percentages of non-neuronal cell types. The coefficient of determination ($R^2$) was calculated to evaluate the accuracy of predicted cell type percentages. Point colours represent different cell subclasses. The identity line is shown.

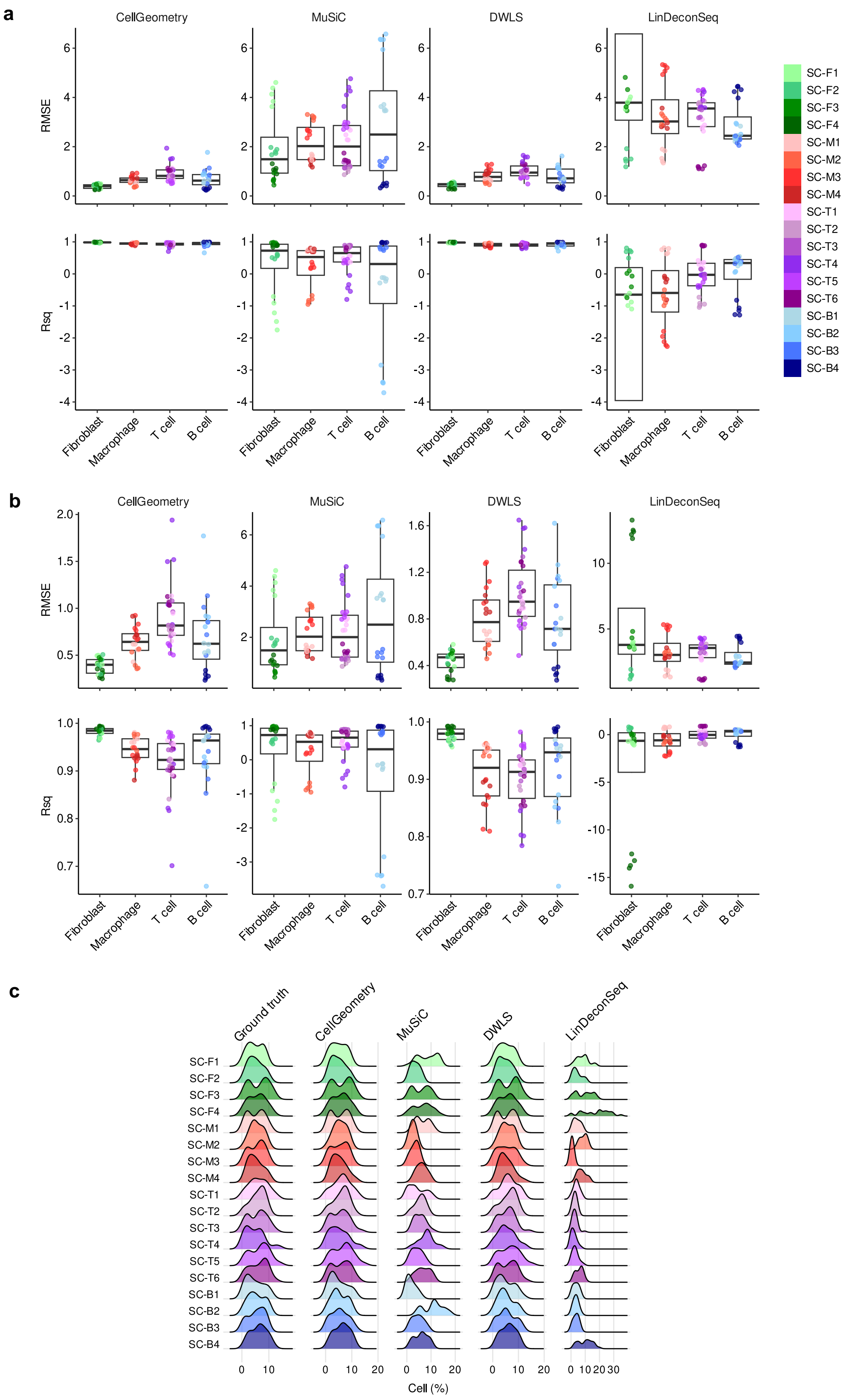


**Supplementary Fig. 12. Benchmarking the accuracy of deconvolution methods against simulated synovium pseudo-bulk data**
**a,** Boxplots of root mean square error (RMSE) and coefficient of determination ($R^2$) for cell subclass percentages from deconvolution of rheumatoid arthritis synovium[1] simulated pseudo-bulk data according to cell type group. Five replicates of simulated data (N = 25 samples) were generated. The deconvolution methods assessed were cellGeometry, MuSiC, DWLS and LinDeconSeq. Boxplots illustrate median, upper and lower quartiles with whiskers denoting maximal and minimal data within 1.5× interquartile range. *y* axis is cropped at the minimum/maximum whisker values to reduce plot distortion by outliers.
**b,** Uncropped plots from panel (**a**). **c,** Ridgeline plots of the ground truth cell proportions and predicted cell subclass proportions from deconvolution of simulated synovium pseudo-bulk data.

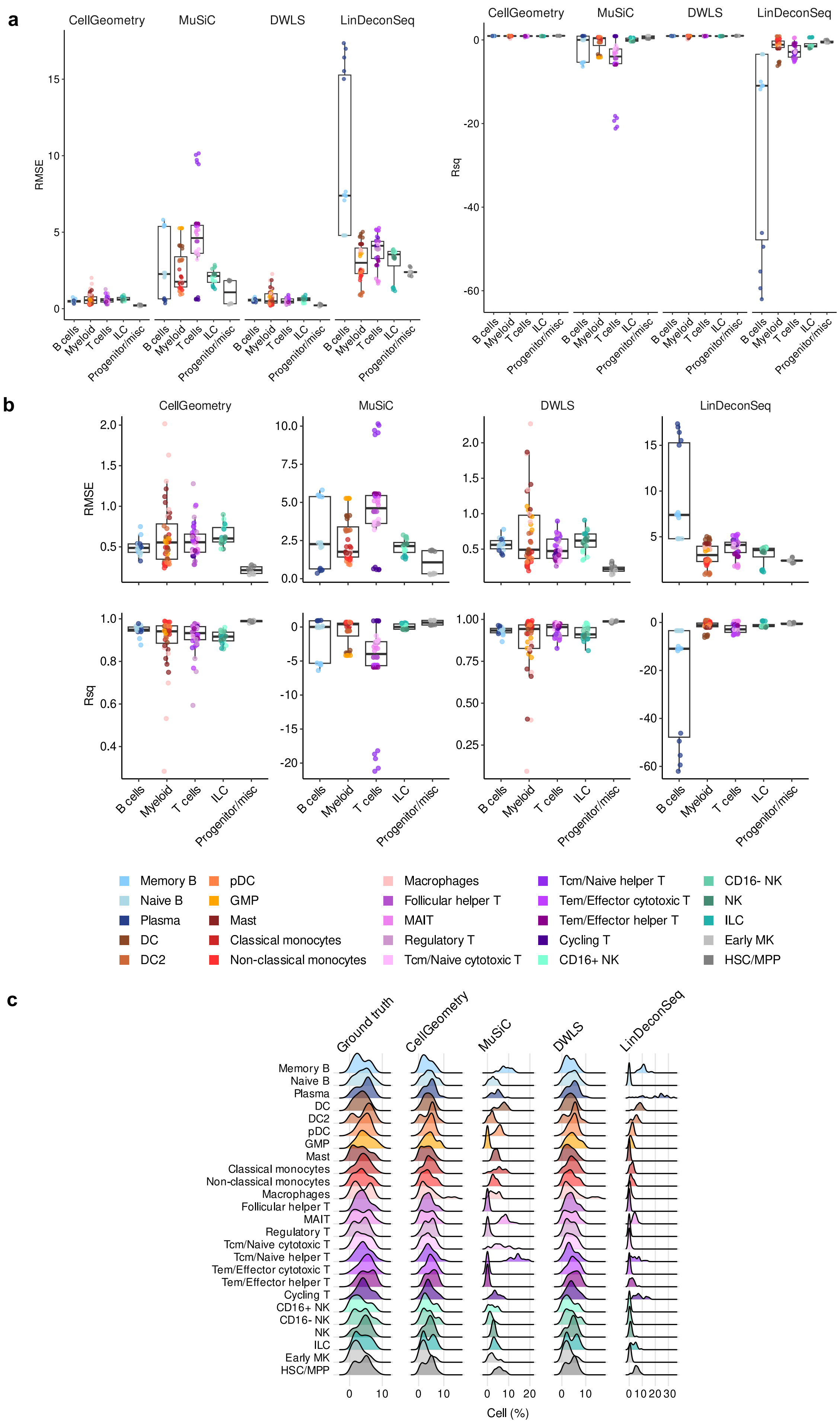


**Supplementary Fig. 13 continued overleaf**

**Supplementary Fig. 13. Benchmarking deconvolution accuracy with simulated pseudo-bulk from Cell Typist single-cell data**
**a,** Boxplot of root mean square error (RMSE) and coefficient of determination ($R^2$) for cell subclass percentages from deconvolution of Cell Typist blood[2] simulated pseudo-bulk data according to cell group. Five replicates of simulated data (N=25 samples) were generated. Deconvolution methods assessed were cellGeometry, MuSiC, DWLS and LinDeconSeq. Boxplots illustrate median, upper and lower quartiles; whiskers denote maximal and minimal data within 1.5× interquartile range. *y* axis is cropped at the minimum/maximum whisker values to reduce plot distortion by outliers.
**b,** Full boxplots of the RMSE and $R^2$ for cell subclass percentages from deconvolution of Cell Typist blood simulated data.
**c,** Ridgeline plots of ground truth and predicted cell subclass proportions from deconvolution of Cell Typist simulated pseudo-bulk.

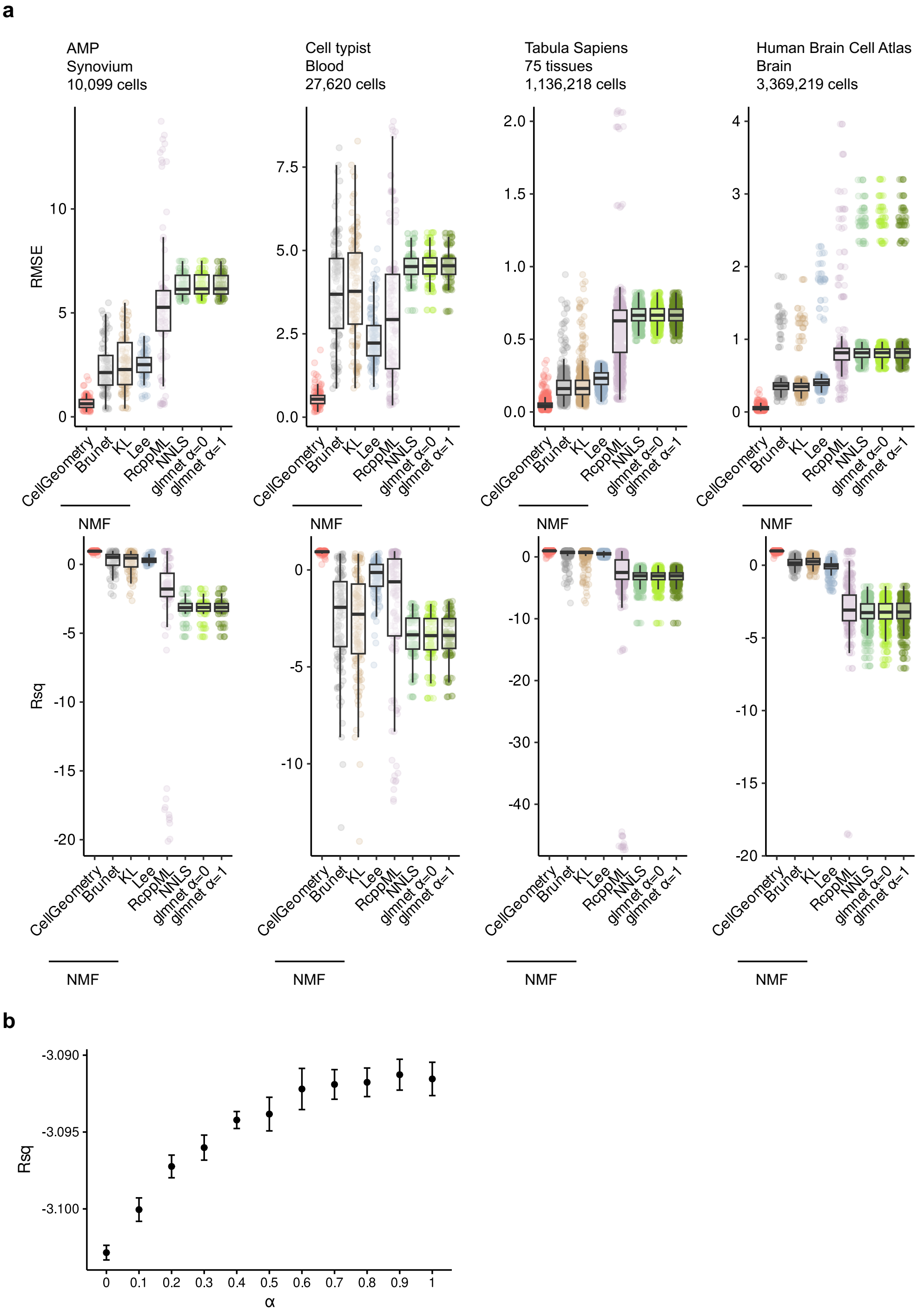


**Supplementary Fig. 14. Comparing deconvolution accuracy of cellGeometry against non-matrix factorisation (NMF), non-negative least squares (NNLS) and generalised linear modelling**

**a,** Boxplots of the root mean squared error (RMSE) and coefficient of determination (Rsq) for cell subclass percentages from the deconvolution of the different simulated pseudo-bulk datasets generated based on Accelerated Medicines Partnership (AMP) rheumatoid arthritis synovium[1], Cell Typist blood[2], Tabula Sapiens[3] and Human Brain Cell Atlas[4] single-cell datasets. CellGeometry was compared against the NMF, NNLS and generalised linear modelling. NMF was performed using NMF package (method = Brunet, Kullback-Leibler (KL) and Lee) and the RcppML package. Generalised linear modelling were performed using glmnet package with alphas 0 (ridge) and 1 (LASSO). Five replicates of simulated data (N = 25 samples) were generated, apart from Human Brain Cell Atlas where 3 replicates (N = 30 samples) were generated. The boxplots illustrate median, upper and lower quartiles with whiskers denoting maximal and minimal data within 1.5× interquartile range.

**b,** Rsq for cell subclass percentages from the deconvolution of simulated pseudo-bulk datasets generated based on Cell Typist blood using glmnet package with a range of alphas. Five replicates of simulated data (N = 25 samples). Point represents mean and error bars represent standard error of the mean (SEM).

**a**

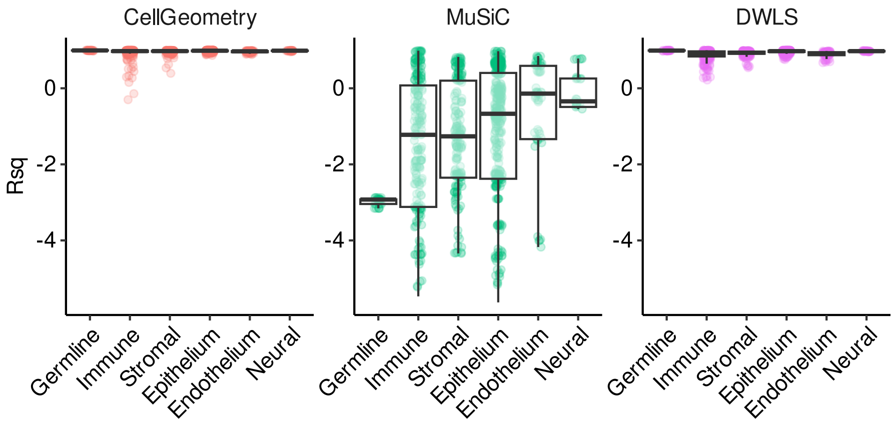


**b**

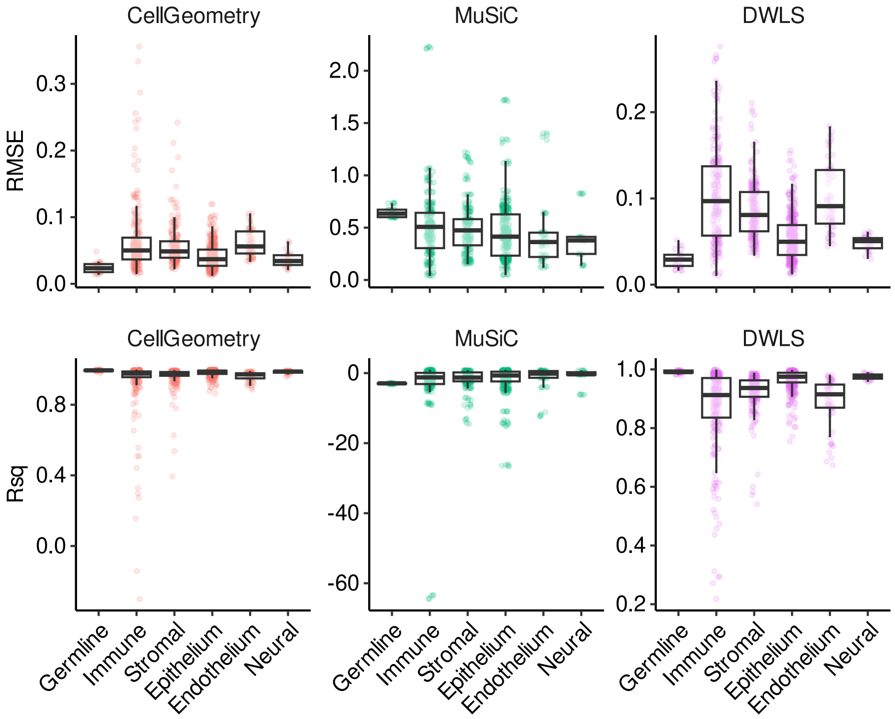


**Supplementary Fig. 15. Comparing the accuracy of cellGeometry with MuSiC and DWLS at deconvoluting simulated pseudo-bulk from Tabula Sapiens**

**a,** Boxplot of the coefficient of determination (Rsq) for the cell subclass percentages from the deconvolution of the Tabula Sapiens[3] simulated data according to the cell type group. Five replicates of simulated data (N = 25 samples) were generated. The deconvolution methods assessed were cellGeometry, MuSiC and DWLS. The boxplots illustrate median, upper and lower quartiles with whiskers denoting maximal and minimal data within 1.5× interquartile range. The *y* axis for each simulated data has been cropped just after the largest maximum whisker value so the plots are not distorted by outliers.

**b,** Full boxplots of the root mean squared error (RMSE) and Rsq for the cell subclass percentages from the deconvolution of the Tabula Sapiens simulated data according to the cell type group.

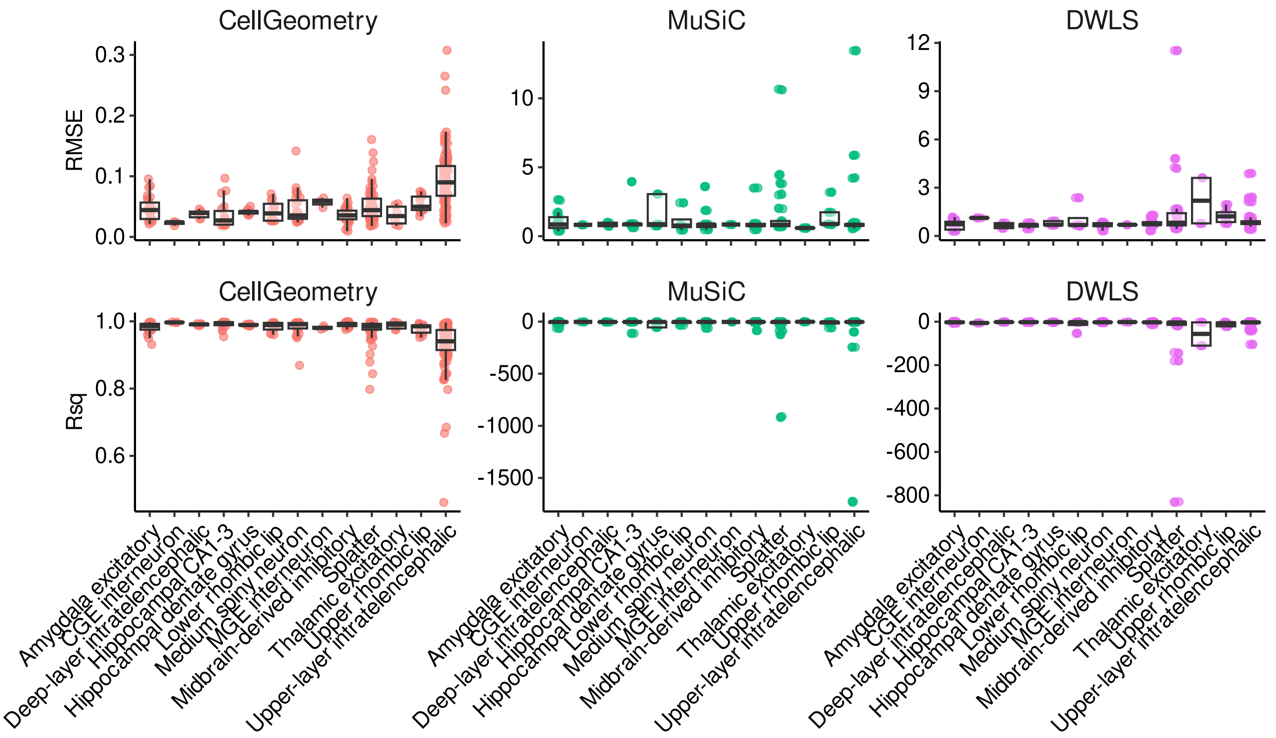


**Supplementary Fig. 16. Comparison of the deconvolution accuracy of cellGeometry with MuSiC and DWLS for neurons from simulated brain pseudo-bulk samples**

Boxplot of the root mean squared error (RMSE) or coefficient of determination (Rsq) for the cell subclass percentages from the deconvolution of neurons from Human Brain Cell Atlas[4] simulated pseudobulk data according to cell type group. Three replicates of simulated data (N = 30 samples) were generated. The deconvolution methods assessed were cellGeometry, MuSiC and DWLS. The boxplots illustrate median, upper and lower quartiles with whiskers denoting maximal and minimal data within 1.5× interquartile range.

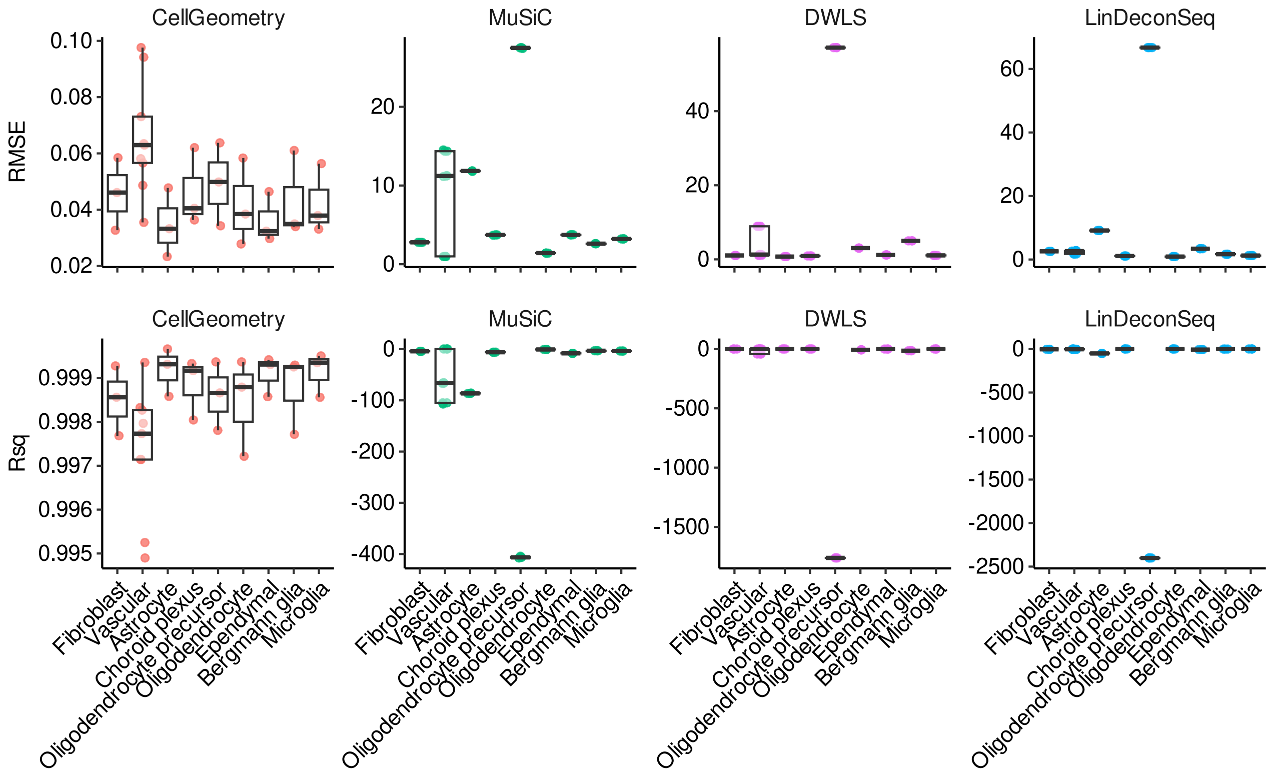


**Supplementary Fig. 17. Benchmarking deconvolution accuracy with non-neuronal cells from simulated brain pseudo-bulk samples**

Boxplot of the root mean square error (RMSE) or coefficient of determination ($R^2$) for cell subclass percentages from the deconvolution of Human Brain Cell Atlas[4] non-neuronal cells according to cell group. Three replicates of simulated data (N = 30 samples) were generated. The deconvolution methods assessed were cellGeometry, MuSiC, DWLS and LinDeconSeq. Boxplots illustrate median, upper and lower quartiles; whiskers denote maximal and minimal data within 1.5× interquartile range.

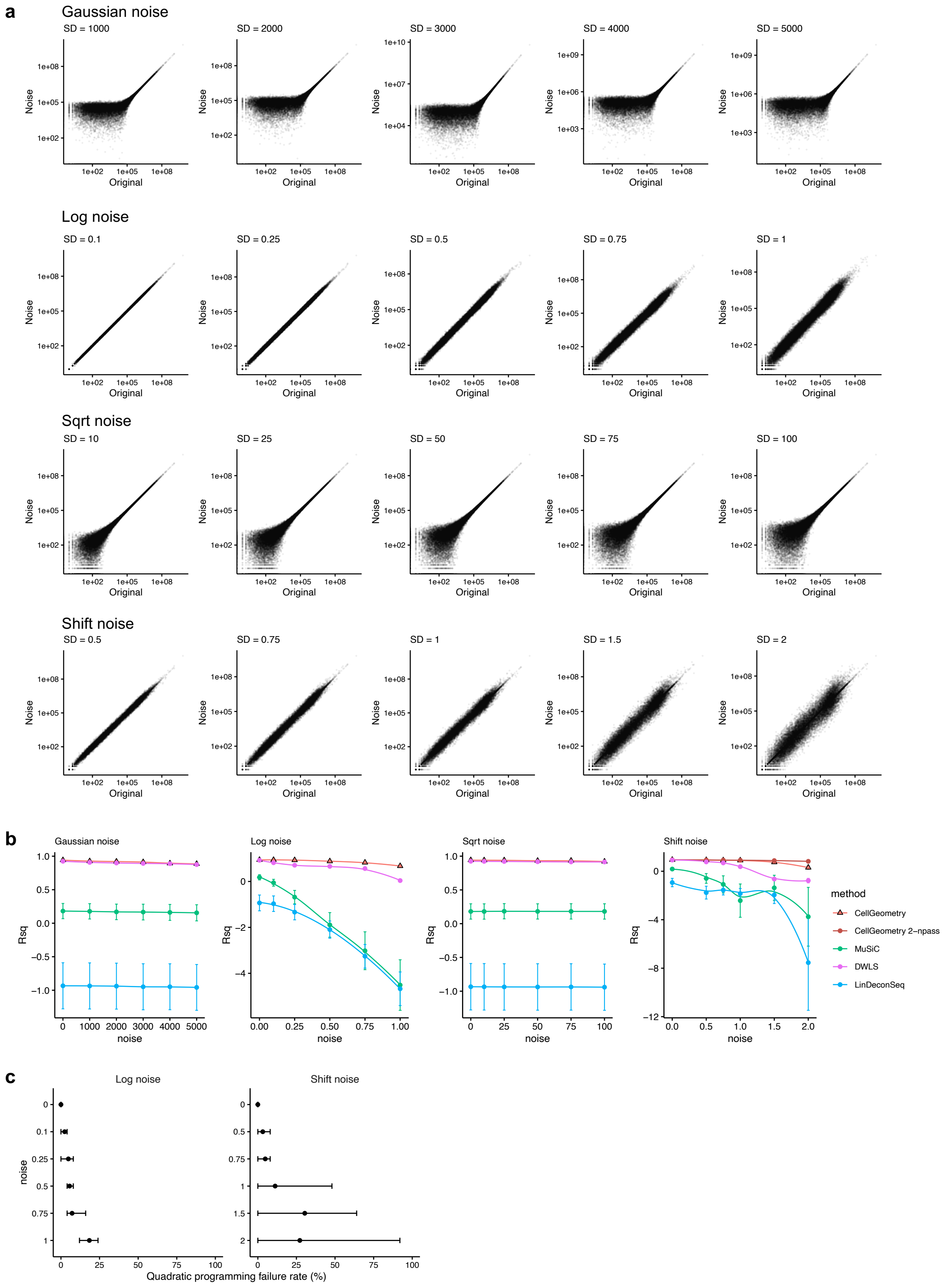


**Supplementary Fig. 18. Benchmarking deconvolution methods against simulated synovium pseudo-bulk with different types of added noise to simulate different sources of experimental assay noise**

**a,** Scatter plot illustrating the different types of noise with varying standard deviation through comparing it to the gene counts from the original simulated data of Accelerated Medicines Partnership (AMP) rheumatoid arthritis synovium[1]. Noise types: addition of Gaussian noise to counts; counts are converted to log2+1 scale before the addition of Gaussian noise and counts are converted back to count scale (Log noise); counts are square rooted before addition of Gaussian noise and then transformed back to count scale (Sqrt noise); genes are randomly selected and multiplied by a random amount (shift noise). Sqrt noise resembles experimental noise observed by Brennecke et al[5]. Shift noise mimics systematic differences in sequencing chemistries between single-cell and bulk.
**b,** Mean coefficient of determination ($R^2$) for cell subclass percentages from the deconvolution using cellGeometry and DWLS of simulated data of AMP with varying standard deviation of noise. Five replicates of simulated (N = 25) were tested. Error bars represent standard error of the mean (SEM)
**c,** Plot of the quadratic programming failure rate in DWLS. Points show mean failure rate, whiskers show range.

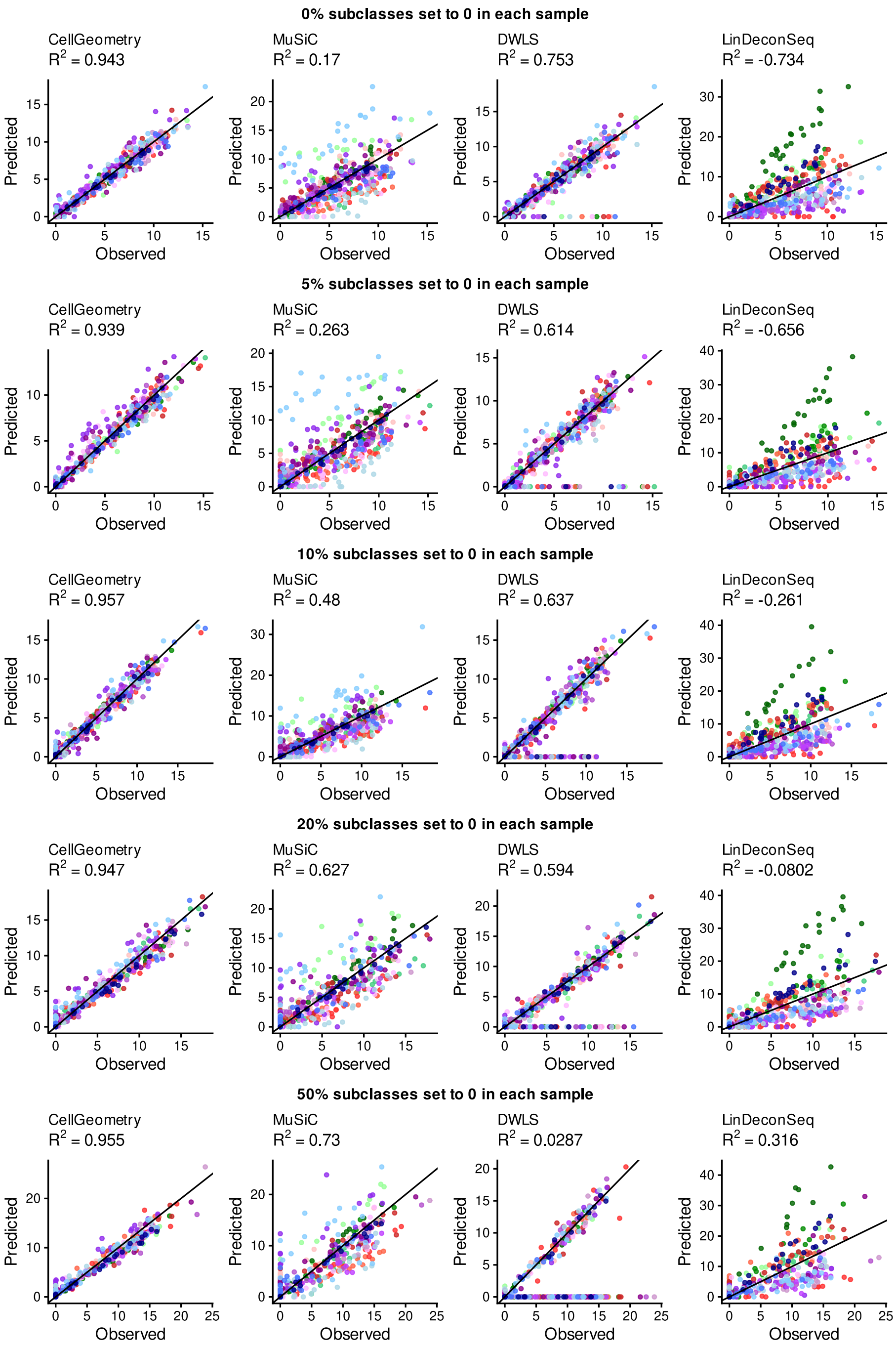


**Supplementary Fig. 19. cellGeometry accuracy is less affected by absent cell types compare to other methods**
Scatter plots showing predicted versus true cell percentages of simulated data of Accelerated Medicines Partnership (AMP) rheumatoid arthritis synovium[1] with different proportions of cell subclasses that are set to 0 in each sample. Deconvolution was undertaken using cellGeometry, MuSiC, DWLS and LinDeconSeq. The coefficient of determination ($R^2$) was calculated to evaluate the accuracy of predicted cell type percentages. Point colours represent different cell subclasses. The identity line is shown.

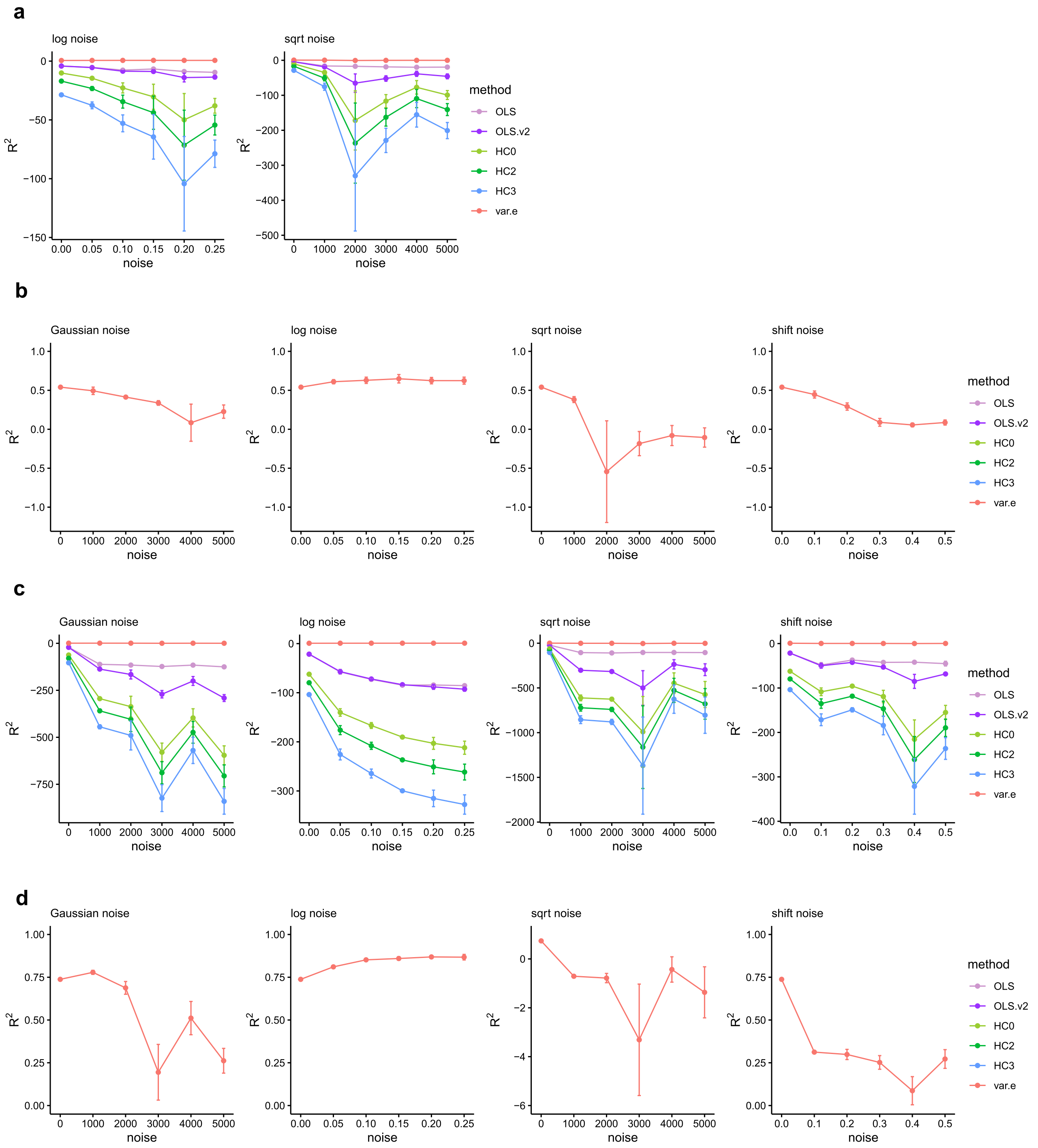


**Supplementary Fig. 20. Determining the optimal method for calculating standard error of cell counts** The mean coefficient of determination ($R^2$) for the deconvoluted cell frequency standard error and the actual error (root mean square error – RMSE) when deconvoluting simulated data of **(a, b)** Tabula Sapiens[3] or **(c, d)** the Human Brain Cell Atlas[4]. Standard error of the cell counts were determined by analogy to ordinary least squares (OLS), OLS using non-negative modified compensation matrix (OLS.v2), heteroscedasticity-consistent SE (HC0, HC2, HC3) and row variance of residuals matrix per gene (var.e) method [see Methods for details]. **(b and d)** have the *y* axis zoomed in and restricted to just the range of $R^2$ using var.e where the corresponding full boxplots are Fig. 4e/Supplementary Fig. 20a and Supplementary Fig. 20c, respectively.

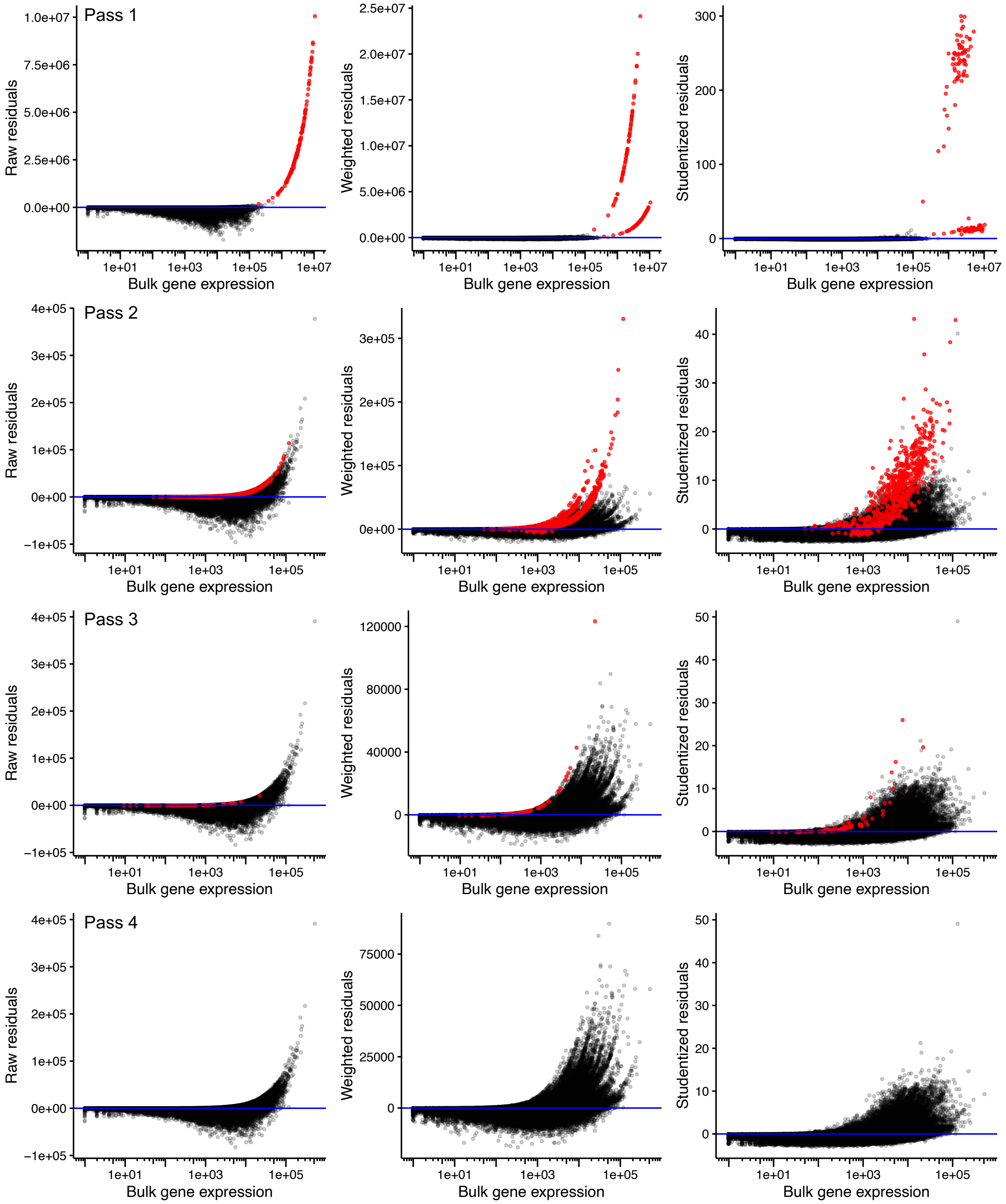


**Supplementary Fig. 21. Assessing bulk gene expression with different residuals for the number of passes of deconvolution.**

Scatter plots of the bulk gene expression and the raw (left column), weighted (middle column) or Studentized weighted residuals (right column) for the number of passes of deconvolution (1, 2, 3 or 4) by cellGeometry. With each pass, outlying genes with excess variance of the residuals (red) are removed. Deconvolution was undertaken with the whole blood RNA-Seq from RA patients in the PEAC study[6] using the Cell Typist blood[2] dataset.

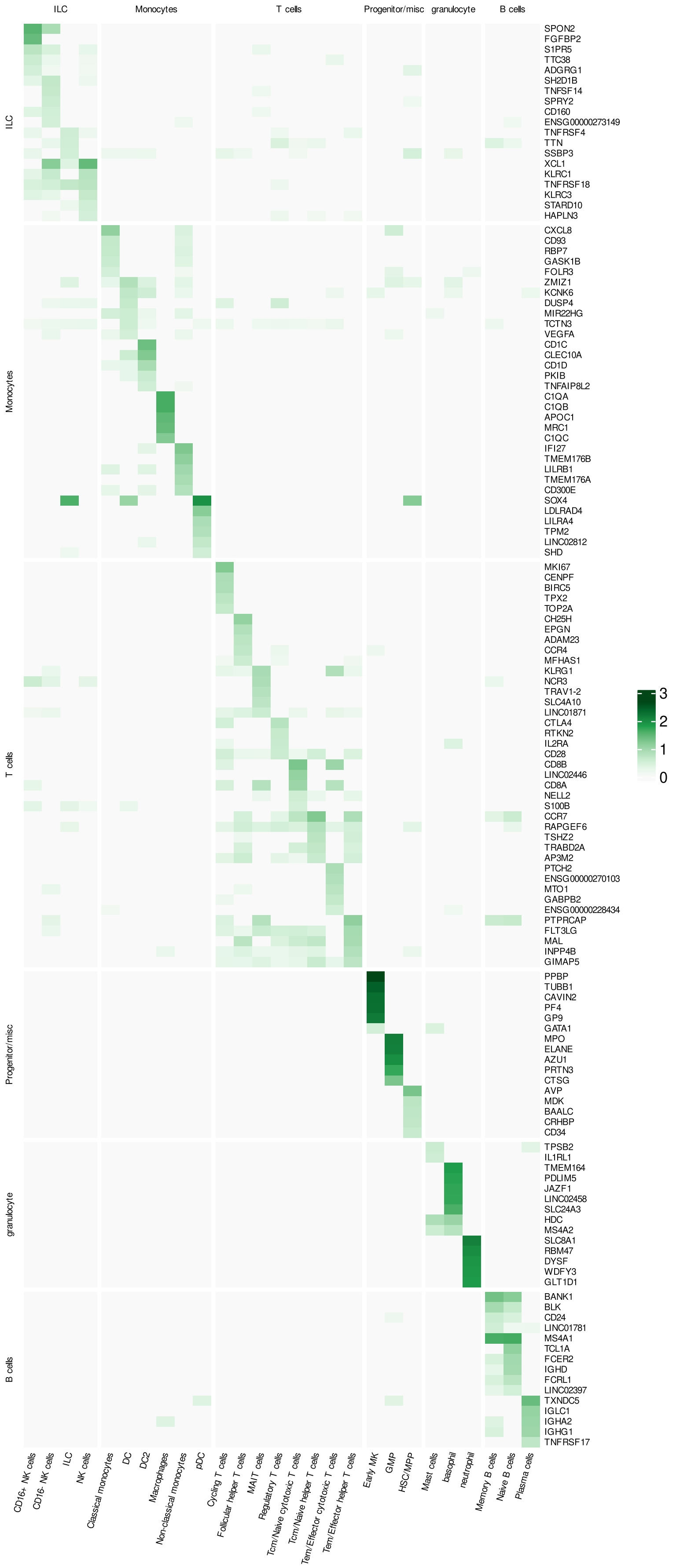


**Supplementary Fig. 22. Gene signature of merged Cell Typist blood and basophils and neutrophils from Tabula Sapiens blood**

Gene signature matrix curated by cellGeometry whereby the Cell Typist blood[2] dataset was merged with basophils and neutrophils from Tabula Sapiens blood[3] through quantile-quantile mapping of the signature matrices to adjust for the differences in the overall level of gene expression. Heatmap shows mean gene expression for top 5 most cell-specific genes per cell subclass.

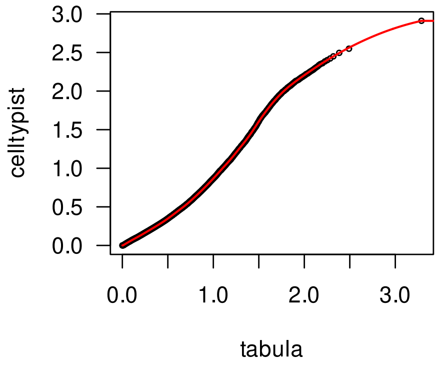


**Supplementary Fig. 23. Quantile-quantile plot mapping the CellTypist and Tabula Sapiens scRNA-Seq datasets**

Quantile-quantile plot comparing Tabula Sapiens blood[3] and Cell Typist blood[2] mean cell type gene expression on a log2+1 scale.

**a**

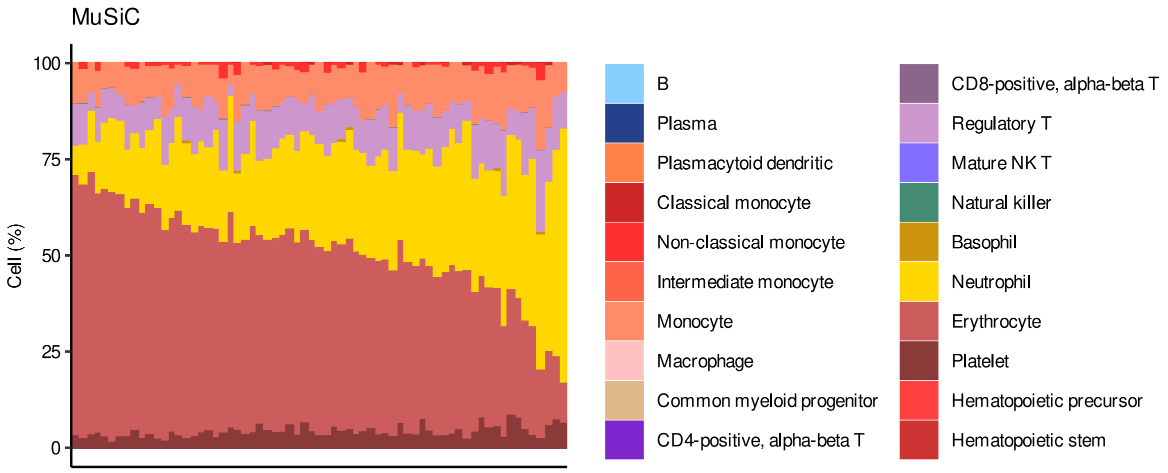


**b**

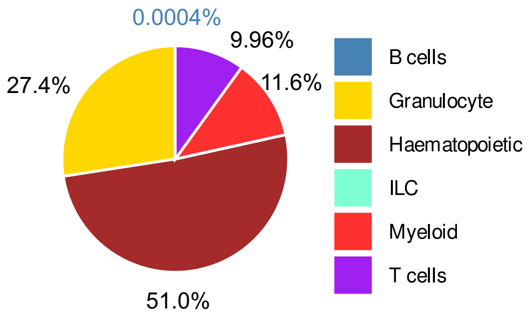


**Supplementary Fig. 24. Tabula Sapiens blood deconvolution by MuSiC in real-world whole blood samples**

**a,** Stacked barplots of the Tabula Sapiens[3] blood cell subclass proportions predicted by MuSiC in whole blood RNA-Seq from early RA patients in the PEAC study[6].

**b,** Pie chart of the mean blood cell type group proportions.

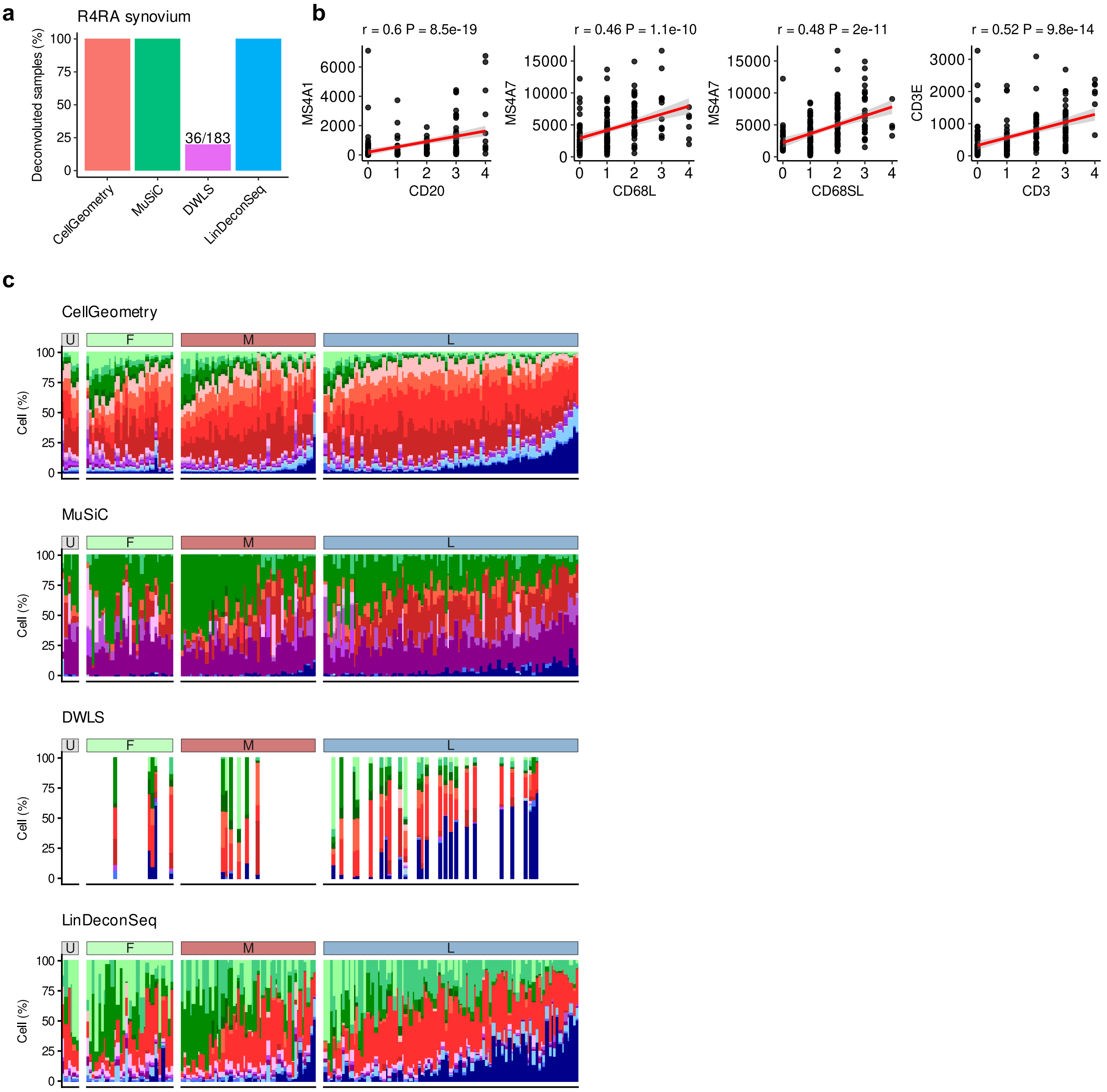


**Supplementary Fig. 25. Deconvoluting rheumatoid arthritis (RA) synovium and evaluating gene and protein cell marker expression.**

**a,** Barplot showing the completion rate for deconvolution of real bulk RNA-Seq of 183 synovial biopsies from the R4RA trial[7] comparing different methods. Rheumatoid arthritis synovium scRNA-Seq[1] was used as the reference dataset and deconvolution was undertaken using cellGeometry, MuSiC, DWLS and LinDeconSeq.

**b,** Scatter plots of gene expression from bulk RNA-Seq showing cell specific gene expression markers (*y* axis) and their respective histology cell surface markers (*x* axis) as measured by semi-quantitative immunohistology in RA synovial biopsies. *MS4A1* versus CD20 histology for B cells; *MS4A7* versus CD68L or CD68SL histology for macrophages; and *CD3E* versus CD3 histology for T cells. P value and r coefficient were calculated using Spearman's correlation test.

**c,** Stacked barplots of RA synovium cell subclass proportions in bulk RNA-seq of synovial biopsies from RA patients from the R4RA trial as deconvoluted by cellGeometry, MuSiC, DWLS and LinDeconSeq. Each stacked column corresponds to an individual synovial biopsy and grouped by histology-based pathotype class (U = ungraded, F = fibroid, M = myeloid, L = lymphoid). The order of individuals is consistent between the different deconvolution methods.

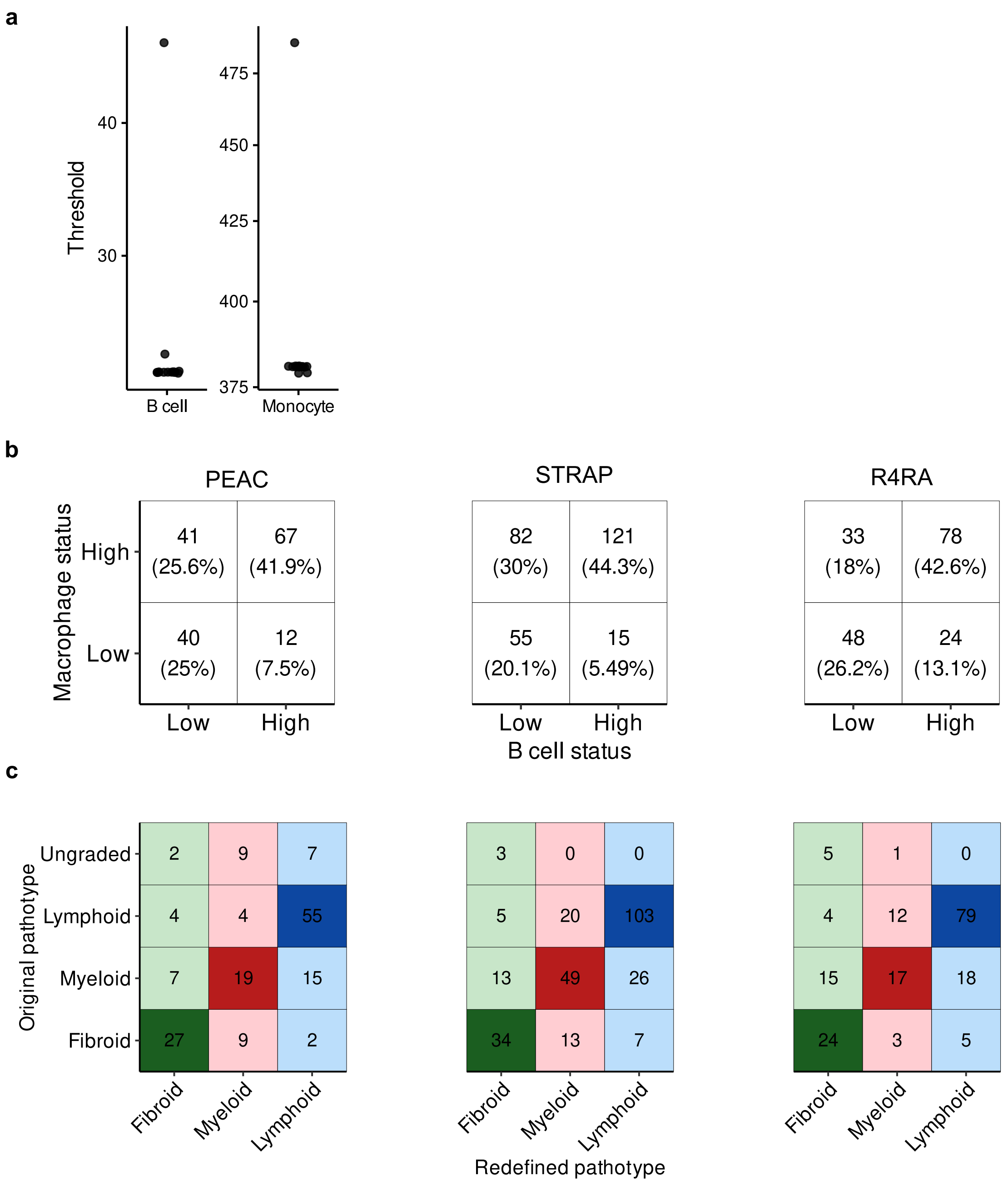


**Supplementary Fig. 26. The use of relative B cell and macrophage abundance from cellGeometry in redefining rheumatoid arthritis (RA) synovium.**

Deconvolution of RA synovium from the PEAC[6], STRAP[8] and R4RA[7] trials were undertaken by cellGeometry using the Accelerated Medicines Partnership (AMP) RA synovium scRNA-Seq dataset[1].

**a,** Thresholds from decision tree-based prediction of histological pathotypes using cellGeometry's B cell and macrophage cell frequencies to test thresholds for pathotype redefinition.

**b,** Contingency table on the macrophage and B cell status where the group abundance thresholds are 30 and 300, respectively.

**c,** Contingency table on the original pathotypes based on histology and the redefined pathotypes based on the B cell and macrophage relative abundance from cellGeometry.

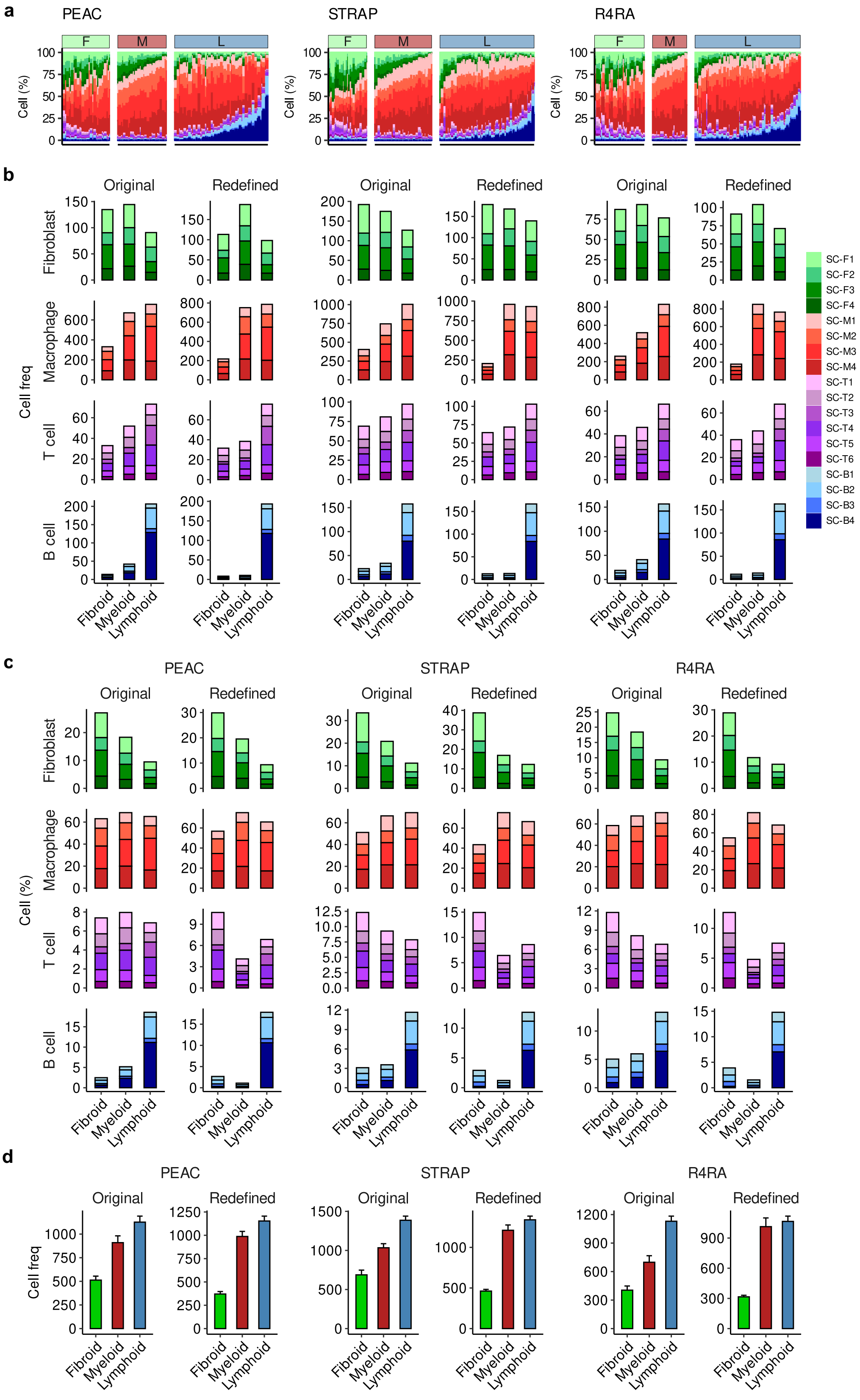


**Supplementary Fig. 27. Comparing the original histology-based pathotype and the redefined cellGeometry based pathotype**
Deconvolution of rheumatoid arthritis (RA) synovium from the PEAC[6], STRAP[8] and R4RA[7] trial were undertaken by cellGeometry using the Accelerated Medicines Partnership (AMP) RA synovium single-cell dataset as reference[1].

**a,** Stacked barplot of AMP cell subclass proportions predicted by cellGeometry. Each stacked column corresponds to a synovium biopsy, grouped by redefined pathotype based on the B cell and macrophage abundance from cellGeometry. The three pathotypes are: Lymphoid (L), high lymphocyte and myeloid cells; myeloid (M), macrophage rich; fibroid (F), low immune cells.

**b, c, d** Stacked barplots of **(b)** mean relative cell subclass frequency, **(c)** mean relative cell proportion and **(d)** mean total cell frequency comparing original and redefined pathotypes based on B cell and macrophage abundance from cellGeometry, showing that the redefined pathotypes exhibit stronger differentiation of **(b)** B/T cell, macrophage and **(c)** fibroblast amounts between pathotypes and the difference in cellularity **(d)** is clearer.

| Data | Method | RMSE | $R^2$ | CCC |
|---|---|---|---|---|
| **AMP** | CellGeometry | 0.757 ± 0.0132 | 0.943 ± 0.00198 | 0.972 ± 0.0011 |
| | MuSiC | 2.74 ± 0.0416 | 0.251 ± 0.0228 | 0.668 ± 0.00867 |
| | DWLS | 0.857 ± 0.0344 | 0.926 ± 0.00574 | 0.964 ± 0.00281 |
| | LinDeconSeq | 4.34 ± 0.046 | -0.881 ± 0.0401 | 0.491 ± 0.00616 |
| **Cell Typist** | CellGeometry | 0.632 ± 0.022 | 0.921 ± 0.00557 | 0.96 ± 0.00264 |
| | MuSiC | 3.75 ± 0.0117 | -1.79 ± 0.0174 | 0.226 ± 0.00151 |
| | DWLS | 0.658 ± 0.0403 | 0.913 ± 0.0106 | 0.958 ± 0.00491 |
| | LinDeconSeq | 4.92 ± 0.0653 | -3.81 ± 0.127 | 0.2 ± 0.0045 |
| **Tabula Sapiens** | CellGeometry | 0.0623 ± 0.00124 | 0.965 ± 0.00139 | 0.983 ± 0.000717 |
| | MuSiC | 0.571 ± 0.000341 | -1.91 ± 0.00347 | 0.295 ± 0.000277 |
| | MuSiC (downsampled) | 0.483 ± 0.000266 | -1.08 ± 0.00229 | 0.304 ± 0.00067 |
| | DWLS | 0.0882 ± 0.000161 | 0.931 ± 0.000253 | 0.963 ± 0.000143 |
| | Bisque | 1.43 ± 0.00156 | -17.2 ± 0.0396 | 0.161 ± 0.000516 |
| | InstaPrism | 0.21 ± 5.16e-05 | 0.609 ± 0.000193 | 0.751 ± 0.000136 |
| **Human Brain Cell Atlas (Neuron)** | CellGeometry | 0.0716 ± 0.00338 | 0.968 ± 0.003 | 0.984 ± 0.0015 |
| | MuSiC | 2.25 ± 0.0016 | -30.5 ± 0.0449 | 0.0206 ± 8.47e-05 |
| | MuSiC (downsampled) | 1.76 ± 0.000741 | -18.3 ± 0.0162 | 0.0236 ± 0.000106 |
| | DWLS | 1.77 ± 0.000834 | -18.4 ± 0.0183 | 0.0317 ± 7.77e-05 |
| | Bisque | 3.08 ± 0.00362 | -57.8 ± 0.138 | 0.0338 ± 8.12e-05 |
| | InstaPrism | 0.609 ± 3.6e-05 | -1.31 ± 0.000273 | 0.137 ± 0.00015 |
| **Human Brain Cell Atlas (Non-neuronal)** | CellGeometry | 0.051 ± 0.00659 | 0.998 ± 0.000382 | 0.999 ± 0.000197 |
| | MuSiC | 10.8 ± 0.00789 | -65 ± 0.0964 | 0.0327 ± 0.00011 |
| | MuSiC (downsampled) | 13.1 ± 0.0006 | -96.2 ± 0.00889 | 0.0264 ± 5.78e-06 |
| | DWLS | 17.6 ± 0.000934 | -173 ± 0.0185 | 0.0222 ± 6.01e-06 |
| | LinDeconSeq | 20.4 ± 0.000391 | -234 ± 0.00902 | 0.0167 ± 3.41e-06 |
| | Bisque | 8.27 ± 0.000435 | -37.7 ± 0.00407 | 0.0932 ± 6.2e-05 |
| | InstaPrism | 16.8 ± 0.00059 | -159 ± 0.0112 | 0.0209 ± 2.83e-06 |

**Supplementary Table 1. Evaluation metrics summary for deconvoluting simulated data**
The mean of root mean square error (RMSE), coefficient of determination ($R^2$) and Lin's Concordance Correlation Coefficient (CCC) alongside the standard error of the mean (SEM) are detailed for the agreement between predicted and true cell percentages from deconvoluting simulated pseudo-bulk datasets generated from Accelerated Medicines Partnership (AMP) rheumatoid arthritis synovium[1], Cell Typist blood[2], Tabula Sapiens[3] and Human Brain Cell Atlas[4] scRNA-Seq datasets. Deconvolution methods assessed were cellGeometry, MuSiC, DWLS, LinDeconSeq, Bisque and InstaPrism.

| Cluster | CellGeometry | MuSiC | DWLS | LinDeconSeq |
|---|---|---|---|---|
| **Memory B** | 2.02 (0.00 - 4.13) | 0.00 (0.00 - 0.00) | 0.25 (0.00 - 1.13) | 2.58 (0.00 - 10.2) |
| **Naive B** | 1.68 (0.00 - 5.15) | 0.00 (0.00 - 0.00) | 0.68 (0.00 - 2.33) | 1.57 (0.00 - 10.4) |
| **Plasma** | 1.01 (0.00 - 2.21) | 0.73 (0.05 - 2.22) | 0.08 (0.00 - 0.79) | 0.94 (0.00 - 3.40) |
| **DC** | 1.96 (0.00 - 5.19) | 0.00 (0.00 - 0.00) | 0.00 (0.00 - 0.05) | 0.00 (0.00 - 0.00) |
| **DC2** | 2.42 (0.00 - 3.80) | 0.00 (0.00 - 0.22) | 0.00 (0.00 - 0.11) | 2.24 (0.00 - 4.04) |
| **pDC** | 1.09 (0.00 - 2.11) | 0.00 (0.00 - 0.00) | 0.15 (0.00 - 0.76) | 0.52 (0.18 - 0.94) |
| **Classical monocytes** | 9.4 (0.00 - 20.2) | 26.8 (0.14 - 76.9) | 0.44 (0.00 - 19.1) | 14.4 (1.27 - 37.1) |
| **Non-classical monocytes** | 10.5 (6.2 - 18.5) | 27.0 (1.56 - 45.2) | 68.7 (0.00 - 97.6) | 0.32 (0.00 - 3.19) |
| **Macrophages** | 1.11 (0.00 - 1.79) | 0.22 (0.00 - 8.30) | 0.69 (0.00 - 25.7) | 0.00 (0.00 - 0.00) |
| **Follicular helper T** | 2.39 (0.00 - 4.98) | ND*** | 0.35 (0.00 - 1.56) | 0.00 (0.00 - 0.00) |
| **MAIT** | 1.96 (0.32 - 9.78) | 0.00 (0.00 - 0.06) | 0.39 (0.00 - 4.43) | 1.60 (0.00 - 5.09) |
| **Regulatory T** | 4.18 (1.79 - 13.1) | 0.00 (0.00 - 0.00) | 0.55 (0.00 - 2.85) | 1.00 (0.00 - 6.36) |
| **Tcm/Naive cytotoxic T** | 4.80 (0.00 - 7.42) | 0.00 (0.00 - 0.00) | 0.05 (0.00 - 0.54) | 1.13 (0.00 - 5.86) |
| **Tcm/Naive helper T** | 3.16 (0.00 - 7.07) | 0.00 (0.00 - 0.00) | 0.01 (0.00 - 0.25) | 26.4 (0.32 - 42.4) |
| **Tem/Effector cytotoxic T** | 1.66 (0.63 - 2.75) | ND*** | 0.33 (0.00 - 10.2) | 0.00 (0.00 - 0.00) |
| **Tem/Effector helper T** | 5.71 (0.16 - 8.86) | 0.07 (0.00 - 0.26) | 0.67 (0.00 - 2.42) | 0.12 (0.00 - 3.22) |
| **Cytotoxic T** | ND* | 0.06 (0.00 - 3.48) | 1.24 (0.00 - 5.30) | 14.9 (8.80 - 25.7) |
| **Helper T** | ND* | 0.00 (0.00 - 0.00) | 0.00 (0.00 - 0.00) | 0.81 (0.00 - 14.9) |
| **Gamma delta T** | ND** | ND*** | 0.01 (0.00 - 0.08) | 0.00 (0.00 - 0.00) |
| **Cycling T** | 0.67 (0.00 - 1.42) | 0.01 (0.00 - 0.60) | 0.00 (0.00 - 0.01) | 0.00 (0.00 - 0.00) |
| **CD16+ NK** | 4.27 (0.43 - 11.0) | 0.37 (0.00 - 7.14) | 0.47 (0.00 - 5.44) | 0.00 (0.00 - 0.00) |
| **CD16- NK** | 1.61 (0.00 - 2.45) | 0.00 (0.00 - 0.00) | 0.13 (0.00 - 0.92) | 6.66 (1.38 - 18.1) |
| **NK** | 1.89 (0.00 - 3.15) | 23.5 (6.41 - 43.6) | 0.00 (0.00 - 0.00) | 0.00 (0.00 - 0.00) |
| **ILC** | 2.45 (0.07 - 7.49) | 0.00 (0.00 - 0.00) | 0.00 (0.00 - 0.00) | 3.00 (1.54 - 4.03) |
| **Mast** | 2.87 (0.00 - 4.32) | 0.00 (0.00 - 0.00) | 0.37 (0.00 - 2.19) | 1.43 (0.00 - 4.65) |
| **Basophil** | 1.37 (0.48 - 2.21) | | | |
| **Neutrophil** | 23.1 (10.4 - 37.6) | | | |
| **Early MK** | 4.81 (1.23 - 20.5) | 21.2 (9.15 - 36.8) | 24.2 (0.43 – 100) | 0.71 (0.19 - 2.19) |
| **MEMP** | ND** | 0.00 (0.00 - 0.00) | 0.00 (0.00 - 0.02) | 3.28 (0.00 - 6.66) |
| **Promyelocytes** | ND** | ND*** | 0.03 (0.00 - 1.67) | 16.3 (5.40 - 26.3) |
| **HSC/MPP** | 0.53 (0.00 - 1.28) | 0.00 (0.00 - 0.00) | 0.00 (0.00 - 0.00) | 0.00 (0.00 - 0.00) |
| **GMP** | 1.38 (0.00 - 4.12) | ND*** | 0.26 (0.00 - 3.13) | 0.14 (0.02 - 0.68) |

**Supplementary Table 2. Cell subclass percentages from deconvolution of rheumatoid arthritis whole blood bulk RNA-Seq comparing deconvolution methods**

67 whole blood bulk RNA-Seq samples from individuals with rheumatoid arthritis in the PEAC study[6] were used. cellGeometry deconvoluted the bulk blood RNA-Seq using the Cell Typist blood scRNA-Seq[2] signature merged with basophil and neutrophil signatures from Tabula Sapiens[3]. MuSiC, DWLS and LinDeconSeq deconvoluted bulk blood RNA-Seq with Cell Typist blood dataset alone. Cell percentages are shown as mean and range, limited to 2 decimal places. Cell subclasses with outputs of zero (<0.005%) across all samples are highlighted in red. * These subclasses were removed from the Cell Typist blood signature due to their similarity with other closely related CD4 and CD8 T cell subset clusters. ** Gene signatures were not generated for these subclasses because of insufficient (<10) cell numbers in the scRNA-Seq. *** These subclasses were not outputted from the deconvolution algorithm.

| Cluster | MuSiC |
|---|---|
| **B** | 0.00 (0.00 - 0.03) |
| **Plasma** | 0.00 (0.00 - 0.00) |
| **Plasmacytoid dendritic** | 0.00 (0.00 - 0.00) |
| **Classical monocyte** | 0.06 (0.00 - 0.26) |
| **Non-classical monocyte** | 0.55 (0.00 - 3.86) |
| **Intermediate monocyte** | 0.00 (0.00 - 0.00) |
| **Monocyte** | 10.97 (5.71 - 18.69) |
| **Macrophage** | 0.00 (0.00 - 0.00) |
| **Common myeloid progenitor** | 0.00 (0.00 - 0.00) |
| **CD4-positive, alpha-beta T** | 0.00 (0.00 - 0.00) |
| **CD8-positive, alpha-beta T** | 0.00 (0.00 - 0.00) |
| **Naive thymus-derived CD4-positive, alpha-beta T** | 0.07 (0.00 - 0.29) |
| **Regulatory T** | 9.89 (2.32 - 21.20) |
| **Mature NK T** | 0.00 (0.00 - 0.02) |
| **Natural killer** | 0.00 (0.00 - 0.00) |
| **Basophil** | 0.00 (0.00 - 0.14) |
| **Neutrophil** | 27.44 (8.30 - 66.71) |
| **Erythrocyte** | 46.46 (9.79 - 67.62) |
| **Platelet** | 4.55 (1.88 - 8.90) |
| **Hematopoietic precursor** | 0.00 (0.00 - 0.00) |
| **Hematopoietic stem** | 0.00 (0.00 - 0.00) |

**Supplementary Table 3. Deconvolution of whole blood bulk RNA-Seq by MuSiC using Tabula Sapiens scRNA-Seq as reference**
67 whole blood bulk RNA-Seq samples from individuals with rheumatoid arthritis in the PEAC study were used[6]. MuSiC deconvoluted the whole blood bulk samples using Tabula Sapiens blood scRNA-Seq[3] as the reference dataset. Cell percentages are shown as mean and range.

| Cluster | CellGeometry | MuSiC | DWLS | LinDeconSeq |
|---|---|---|---|---|
| **SC-F1** | 4.50 (0.00 - 20.64) | 0.03 (0.00 - 5.70) | 7.94 (0.00 - 58.43) | 12.20 (0.00 - 74.74) |
| **SC-F2** | 3.15 (0.00 - 15.50) | 1.64 (0.00 - 16.45) | 3.46 (0.00 - 13.43) | 15.32 (0.00 - 57.70) |
| **SC-F3** | 4.79 (0.00 - 25.95) | 31.73 (0.05 - 91.89) | 10.78 (0.00 - 51.15) | 14.08 (0.00 - 81.18) |
| **SC-F4** | 2.38 (0.00 - 9.39) | 0.81 (0.00 - 42.54) | 6.75 (0.00 - 39.45) | 1.48 (0.00 - 29.65) |
| **SC-M1** | 9.79 (0.00 - 16.77) | 0.00 (0.00 - 0.05) | 0.69 (0.00 - 15.81) | 0.65 (0.00 - 15.96) |
| **SC-M2** | 12.96 (0.00 - 34.92) | 6.51 (0.00 - 23.89) | 9.07 (0.00 - 35.32) | 1.45 (0.00 - 22.63) |
| **SC-M3** | 22.63 (0.00 - 43.66) | 0.78 (0.00 - 18.72) | 28.93 (1.20 - 67.95) | 32.52 (0.06 - 82.19) |
| **SC-M4** | 21.88 (0.00 - 43.86) | 18.37 (0.00 - 66.64) | 5.05 (0.00 - 43.34) | 0.11 (0.00 - 4.49) |
| **SC-T1** | 2.05 (0.00 - 10.08) | 4.33 (0.00 - 61.58) | 0.29 (0.00 - 3.62) | 5.23 (0.00 - 20.30) |
| **SC-T2** | 1.39 (0.00 - 16.04) | 0.15 (0.00 - 6.44) | 0.04 (0.00 - 0.68) | 0.00 (0.00 - 0.00) |
| **SC-T3** | 0.90 (0.00 - 7.79) | 6.86 (0.00 - 30.09) | 0.23 (0.00 - 2.20) | 0.56 (0.00 - 8.83) |
| **SC-T4** | 1.48 (0.00 - 8.61) | 0.00 (0.00 - 0.25) | 0.36 (0.00 - 4.29) | 0.00 (0.00 - 0.00) |
| **SC-T5** | 1.55 (0.00 - 9.73) | 0.70 (0.00 - 23.47) | 0.54 (0.00 - 6.05) | 0.31 (0.00 - 4.24) |
| **SC-T6** | 1.01 (0.00 - 7.67) | 25.64 (5.84 - 51.31) | 0.08 (0.00 - 0.70) | 2.67 (0.00 - 11.80) |
| **SC-B1** | 1.51 (0.00 - 5.45) | 0.00 (0.00 - 0.10) | 0.17 (0.00 - 5.05) | 0.29 (0.00 - 8.01) |
| **SC-B2** | 2.93 (0.00 - 18.98) | 0.07 (0.00 - 13.06) | 0.12 (0.00 - 4.15) | 3.11 (0.00 - 29.48) |
| **SC-B3** | 1.10 (0.00 - 5.61) | 0.10 (0.00 - 3.77) | 0.44 (0.00 - 5.65) | 0.51 (0.00 - 3.66) |
| **SC-B4** | 4.01 (0.04 - 37.05) | 2.28 (0.00 - 22.00) | 25.08 (0.00 - 71.12) | 9.52 (0.00 - 55.24) |

**Supplementary Table 4. Cell subclass percentage estimates from deconvolution of bulk RNA-Seq of rheumatoid arthritis (RA) synovial biopsies comparing different methods**

183 rheumatoid arthritis (RA) synovium bulk RNA-Seq samples from the R4RA trial[7] were deconvoluted using the Accelerated Medicines Partnership (AMP) rheumatoid arthritis synovium scRNA-Seq dataset[1] as reference, comparing devolution methods cellGeometry, MuSiC, DWLS and LinDeconSeq. DWLS was only able to deconvolute 36 samples successfully (80.3% failure rate). Cell percentages are shown as mean and range.

| | | | CellGeometry | MuSiC | DWLS | LinDeconSeq |
|---|---|---|---|---|---|---|
| Advantages | Repository | Github | ✓ | ✓ | ✓ | ✓ |
| | | CRAN | ✓ | | ✓ | |
| | Usability | Vignette with dataset examples | ✓ | ✓ | | ✓ |
| | Data class input | Matrix | ✓ | ✓ | ✓ | ✓ |
| | | Sparse | ✓ | ✓ | | |
| | | Delayed | ✓ | ✓ | | |
| | | Seurat | ✓ | | | |
| | Features | Separate class and subclass information incorporation | ✓ | | | |
| | | Signature generation is separate from deconvolution | ✓ | | ✓ | ✓ |
| | | Can update and refine the gene signature | ✓ | | | |
| | | Internal plotting for visualisation and evaluation | ✓ | ✓ | | |
| | | Includes functions to generate simulated pseudo-bulk data | ✓ | ✓ | | |
| | | Outputs cell abundance as well as proportions | ✓ | | | |
| | | Ability to merge 2 or more scRNA-Seq signatures | ✓ | | | |
| | | Internal parallelisation option | ✓ | | | |
| | Error mitigation | Non-negative proportions | ✓ | ✓ | ✓ | ✓ |
| | | Regularly maintained (last Github push within 2 years) | ✓ | ✓ | | ✓ |
| Disadvantages | Issues | Inaccurate with simulations | | X | | X |
| | | Zero cell proportions in some subclasses reported across all samples | | X | | X |
| | | Failure rate of up to 80% with real world bulk samples | | | X | |

**Supplementary Table 5. Comparison of deconvolution methods**
